# Replicated hybrid zones reveal genomic patterns of local adaptation and introgression in spruce

**DOI:** 10.1101/2025.08.08.669264

**Authors:** Gabriele Nocchi, Janek Sendrowski, Andy Shi, Brianne Boufford, Manuel Lamothe, Nathalie Isabel, Samuel Yeaman

## Abstract

Hybridization between species can occur along repeated zones of contact, providing a natural laboratory for studying the interplay between migration and selection, and identifying loci involved in adaptation and reproductive isolation. However, interpreting how evolutionary processes shape genomic patterns can be challenging: repeatability of genotype-environment association alone is not strong evidence for selection, as hybrid zones derived from the same parental species are not evolutionarily independent. Conversely, processes that operate within the middle of each hybrid zone, such as selection driving directional introgression, may be more evolutionarily independent, and therefore provide stronger evidence of selection. Here we compared hybridization and local adaptation patterns between two replicated regions within the western Canada interior spruce hybrid zone: a broad latitudinal transect with gradual environmental variation and a narrow elevational transect with substantial topographical and environmental variation. We discovered a complex pattern of introgression, with strong differences in ancestry maintained even across small spatial scales at several locations along the elevational transect. Despite differences in their spatial scales, the elevational and latitudinal transects revealed strikingly similar genome-wide patterns of differentiation and adaptation, and consistent patterns of directional introgression. We explore the extent to which the evolutionary non-independence of these hybrid zones allows inferences about the role of natural selection and drift in shaping these patterns. Consistent with theory, we found longer genomic tracts in the elevational transect, likely because the steeper environmental gradients over shorter distances limit the rate of mixing by migration and recombination relative to drift and selection.

## Introduction

Introgression, which is the incorporation of alleles from one species or population into the gene pool of another through hybridization and repeated backcrossing, is a common and long-recognized phenomenon in plants (Anderson 1953; Haselhorst and Buerkle 2013; Harrison and Larson 2014), and has also been documented in many animal species (Hedrick 2013; Aguillon et al. 2022). Hybridization and subsequent introgression can have substantial impacts on adaptation, evolvability and speciation, with effects that may be beneficial or deleterious depending on the interaction between genetics and ecology. On one hand, hybridization may introduce genetic variants that enhance the fitness of the recipient population (adaptive introgression), may increase standing genetic diversity boosting a population’s adaptive potential to environmental changes, and may lead to speciation if hybrids become reproductively isolated from parental populations (Rieseberg et al. 2003; Baack and Rieseberg 2007; Becker et al. 2013; Brennan et al. 2014; Hamilton and Miller 2015; Mallet 2015). On the other hand, hybridization may counteract local adaptation with the introduction of maladapted alleles, may lead to unfit or unviable hybrids, and may homogenize differences between populations with incomplete reproductive isolation counteracting speciation (Barton and Hewitt 1985). In extreme circumstances, hybridization may lead to extinction through the complete genetic assimilation of a rare species by a more common congener, such as in the case of *Argyranthemum coronopifolium* in Tenerife (Canary Islands), which was effectively replaced by hybrids and *A. frutescens* within just a few decades from the initial contact (Levin et al. 1996). The impact of introgression therefore depends upon how the hybridizing lineages are each adapted to their local environments, and the extent to which they each carry deleterious load (that could be complemented and relieved by introgression) and globally beneficial alleles (that could introgress by a selective sweep).

Studying patterns of introgression across replicated hybrid zones can provide insights into the importance of selection vs. drift (Rieseberg et al. 1999; Simon et al. 2020; Westram et al. 2021; McFarlane et al. 2024; Semenov et al. 2025): if similar genomic regions exhibit similar patterns of introgression across multiple hybrid zones, this can suggest a consistent role for selection rather than purely stochastic processes. Outside of hybrid zones, shared selection pressures across repeated environmental gradients can lead to similar genotype-environment associations (GEAs), whereas drift will result in differing patterns of association. As long as the repeated gradients are evolutionarily independent, repeated GEA signals can provide evidence of a gene having evolved in response to selection (Whiting et al. 2024).

However, if two previously isolated species come into repeated contact across multiple hybrid zones, these zones are fundamentally non-independent, as they are repeatedly derived from the same parental species pools. If the parental species diverged in allopatry followed by secondary contact in repeated hybrid zones, loci that had differentiated by genetic drift could exhibit repeated associations with the overall environmental gradient that distinguishes the parental species. Thus, repeatability in GEA signals across multiple repeated hybrid zones is not sufficient evidence to implicate natural selection (Lotterhos and Whitlock 2014, 2015; Forester et al. 2018; Leigh et al. 2021; Booker et al. 2023). However, if the regions of the genome that exhibit strong GEA signals also exhibit abnormal introgression in the same direction across multiple hybrid zones, this would be unlikely to occur due to the above-described non-independence, and would implicate natural selection (Rieseberg et al. 1999; Buerkle and Lexer 2008; Gompert and Buerkle 2009; Bierne et al. 2013; Simon et al. 2020; Westram et al. 2021; Langdon et al. 2022). Our aim is to explore the extent of similarity in genomic patterns of introgression in two hybrid zones spanning regions with very different geography: one a broad plateau, and another a narrow and rugged mountainous region. In both regions, introgression occurs between white spruce (*Picea glauca*) and Engelmann spruce (*Picea engelmannii*), but in the former region, the environment distinguishing the two parental species transitions slowly across space, whereas in the latter region, the environmental transition is much narrower and includes considerable fine-scale variation associated with elevation and slope aspect.

White spruce (*Picea glauca*) and Engelmann spruce (*Picea engelmannii*) are wind-pollinated, long-lived commercially and aesthetically important forest tree species in Canada (Rajora and Dancik 2000; De La Torre et al. 2013). White spruce is a subarctic boreal species that can tolerate both low winter temperatures as well as summer drought but is generally restricted to low elevations (Rajora and Dancik 2000; De La Torre et al. 2013).

Engelmann spruce is a high-altitude subalpine forest tree with a more westerly and southerly distribution than white spruce, extending from Southern British Columbia and Alberta in the north, up to Arizona and New Mexico in the south (Rajora and Dancik 2000; De La Torre et al. 2013). Engelmann spruce exhibits low tolerance to elevated temperatures and drought conditions, inhabits wetter environments and can withstand short growing season and high snowfall (Rajora and Dancik 2000; De La Torre et al. 2013). White and Engelmann spruce are closely related species and are thought to have diverged in allopatry during the Pliocene (Lockwood et al. 2013; De La Torre et al. 2014). De La Torre et al. (2014) proposed that their most recent contacts are likely to have started in the Rocky Mountains of southern Colorado and Wyoming at the last glacial maximum (LGM), when both species were displaced south of their current distribution. After the glacial retreat, both species started re-expanding, causing the hybrid zone to gradually shift northwest, reaching British Columbia approximately 14,000 years ago and continuing to expand until reaching its current extent in the late Holocene (De La Torre et al. 2014).

Today, these two species hybridize extensively where their ranges overlap in western Alberta and British Columbia, so that the complex of these two species and their hybrids is commonly referred to as “interior spruce” (Rajora and Dancik 2000). As a result, it can be difficult to distinguish these two species by observation except by the morphology of their cones (Taylor 1959; Rajora and Dancik 2000). Hybrids display clinal genetic variation across the hybrid zone and tend to occupy and exhibit higher fitness in habitats that are ecologically intermediate between those of the parental species, suggesting that the interior spruce hybrid zone in western Canada is maintained according to the bounded hybrid superiority model (Horton et al. 1959; Rajora and Dancik 2000; Ledig et al. 2006; De La Torre et al. 2013, 2014). Local adaptation in interior spruce is well supported by common garden experiments as well as genomic studies, which have also revealed strong differentiation between *P. glauca* and *P. engelmannii* (De La Torre et al. 2013, 2014; Hornoy et al. 2015; Yeaman et al. 2016; Capblancq et al. 2022). However, the axis of introgression aligns with climatic gradients, making it unclear how much of the genomic divergence reflects a history of selection versus neutral drift in the parental species prior to secondary contact (Horton et al. 1959; Rajora and Dancik 2000; Ledig et al. 2006; De La Torre et al. 2013, 2014).

Here, to study hybridization and local adaptation we compare genomic patterns across two datasets: a previously studied broad latitudinal gradient spanning > 1000km throughout Alberta and British Columbia (Fig. 1a) (Yeaman et al. 2016), and a new dataset collected across a narrower elevational zone spanning ∼100km in the Canadian Rocky Mountains (Fig. 1c). Based on casual observations of the elevational zone, forests on high north-facing slopes tend to have moister soil and hold snow longer in the springtime than those on high south-facing slopes and valley bottoms, so we expected this might generate some fine-grained natural selection within each location. To test whether this relatively fine-scale environmental variation could drive adaptation, at seven locations along the valleys along the elevational zone, we sampled from three distinct environments: valley bottoms (< 1600 m), and high elevation north- and south-facing slopes (> 2000 m). To compare the latitudinal and elevational datasets, we analyse FST to characterize patterns of genomic divergence, conduct genotype-environment association (GEA) analyses to identify regions potentially driving local adaptation, and perform genomic cline analyses to detect regions showing abnormal patterns of introgression. We also explore patterns of chromosomal clustering of these statistics in each of the datasets to test whether they are consistent with expectations based on theory of hybrid zones and migration-selection balance. We aim to answer three questions: 1) does local adaptation maintain species divergence across fine-scale environmental variation within the hybrid zone; 2) how similar are genomic patterns of divergence, environmental association, and directional introgression across these very different geographies of hybridization?; and 3) to what extent can we parse evolutionary non-independence of these hybrid zones to identify regions of the genome driving adaptive divergence?

**Fig. 1.**
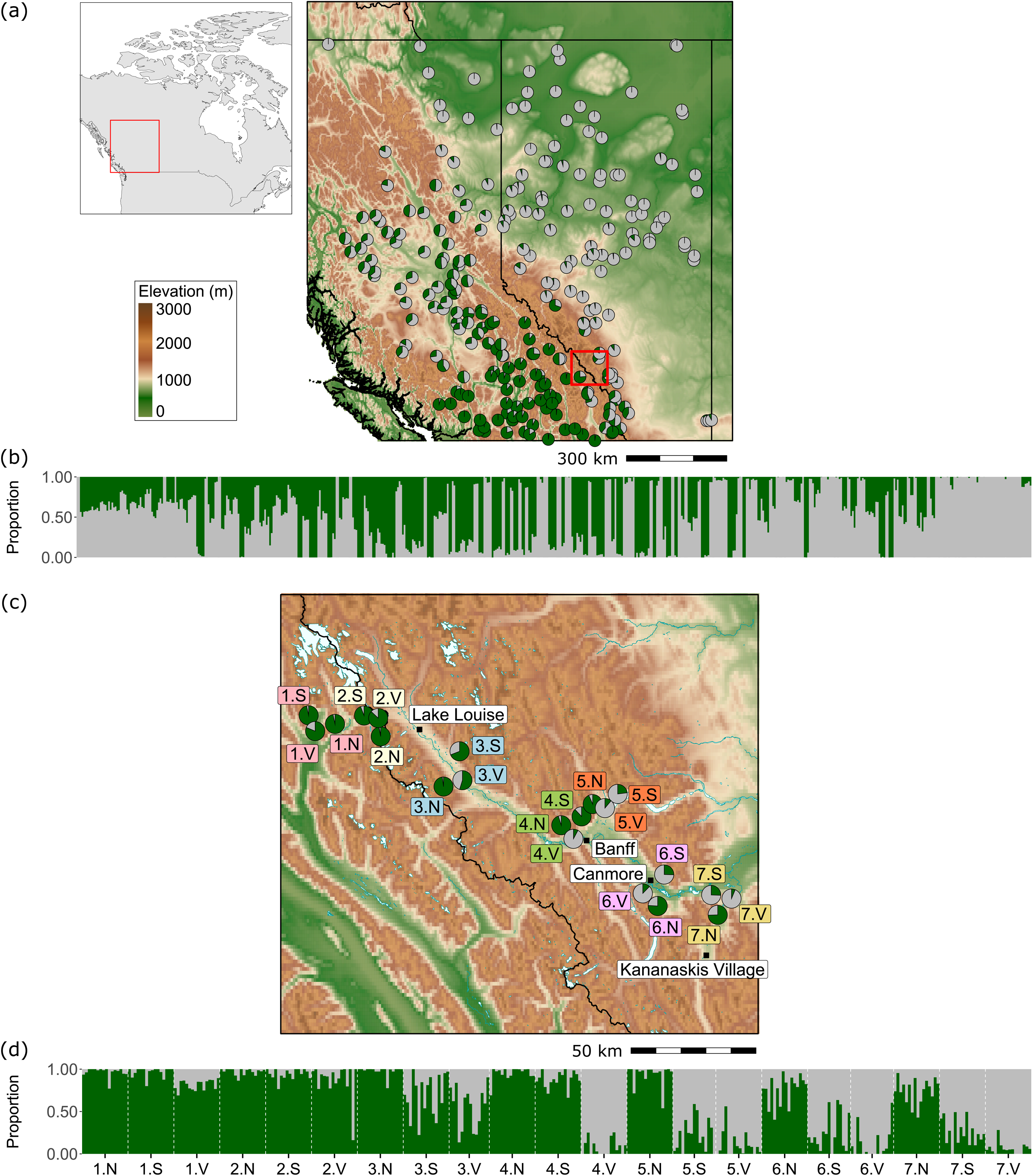
Geographic locations of populations from the (a) latitudinal and (c) elevational datasets in Canada. Pie size reflects the number of individuals per population. Grey indicates *P. glauca* ancestry; green indicates *P. engelmannii* ancestry. The small overview map in (a) shows the location of the latitudinal transect samples in Canada. The red square in (a) marks the location of the elevational transect samples within the broader latitudinal hybrid zone. Panels (b) and (d) show ancestry bar plots based on fastSTRUCTURE results for the latitudinal and elevational datasets, respectively. Individual bars in (b) and (d) are ordered by population longitude.

## Results

### Super-scaffold assembly

While the reference genome assembly of *Picea glauca* is highly fragmented (Warren et al. 2015), we used a previously constructed linkage map based on cDNA sequences to position contigs onto chromosome scale scaffolds, to facilitate study of genome-wide patterns. To maximize confidence, we identified the best scaffold match for each cDNA based on sequence coverage and identity and discarded unmapped cDNA as well as those with low-quality mapping or unresolved multi-mapping (see *Methods* for details). This process retained 21,252 cDNAs, of which 11,914 were included in the available *P. glauca* linkage map, including 14,727 genes spread across 12 linkage groups (LGs) (Pavy et al. 2017; Gagalova et al. 2022). These 11,914 cDNAs mapped to 10,475 unique scaffolds of the *P. glauca* reference genome, which we ordered along the 12 linkage groups according to the cDNAs centimorgan (cM) positions in the linkage map. The resulting super-scaffold assembly spanned 3,426,128,031 bp (∼3.42 Gbp), representing approximately 13% of the WS77111v2 *P. glauca* reference genome assembly (∼26.5 Gbp). The final size of our assembly is comparable to a recently published super-scaffold assembly for *P. glauca*, which achieved a size of ∼3.86 Gbp by using a less conservative cDNA mapping approach (Gagalova et al. 2022).

### Sampling and SNP Discovery

We sampled 331 spruce individuals, including white spruce (*Picea glauca*), Engelmann spruce (*Picea engelmannii*), and their hybrids, from 21 populations within a narrow section of the western Canada interior spruce hybrid zone which extends from central British Columbia (B.C.) into western Alberta (Fig. 1c). Samples were collected from seven locations along the Trans-Canada Highway going through Canmore, Banff, and Yoho. Each location included three populations, with 14 to 16 individuals sampled per population (supplementary fig. S1, Supplementary Material online). For each location, we sampled a north-facing high-altitude population (N), a south-facing high-altitude population (S), and a valley population (V) (Fig. 1c; supplementary fig. S1, Supplementary Material online). Using a target sequencing approach with a 10 Mb target, we identified a set of 119,664 SNPs (MAF > 0.05) following post-processing. This set included only SNPs located on the scaffolds of the *P. glauca* reference genome that we were able to anchor to the *P. glauca* linkage map (Gagalova et al. 2022) in our super-scaffold assembly construction. We termed this dataset the "elevational dataset".

Additionally, we gathered available exome capture data for 575 spruce individuals from 249 populations, derived from previous studies (Suren et al. 2016; Yeaman et al. 2016). This dataset spans both species’ geographic ranges in B.C. and Alberta and encompasses the entire western Canada interior spruce hybrid zone, with the exclusion of areas within national parks (Fig. 1a; supplementary fig. S1, Supplementary Material online). After applying the same SNP-calling pipeline and post-processing as for the elevational dataset to ensure consistency, we identified a set of 265,655 SNPs (MAF > 0.05) located on the scaffolds of our super-scaffold assembly. We termed this dataset the "latitudinal dataset".

### Population structure, ancestry, and fine-scale adaptation

We analysed genome-wide variation in both elevational and latitudinal transects using principal component analysis (PCA) and a variational Bayesian framework implemented in *fastSTRUCTURE* (Raj et al. 2014). In both datasets, *fastSTRUCTURE* identified K=2 as the number of clusters that maximizes the log marginal likelihood (K*ε) of the data (Figs. 1b and d; supplementary fig. S2, Supplementary Material online). Concordantly, PCA’s first principal component (PC1) revealed two main clusters of individuals present in both datasets, while capturing a large portion of the variation in the data (70% in the latitudinal dataset and 63% in the elevational dataset) as indicated by the sharp drop in eigenvalue scores after PC1 (supplementary figs. S3 and S4, Supplementary Material online).

To assign the two clusters identified with *fastSTRUCTURE* and PCA to their respective species, we added publicly available genomic data of a pure individual of *P. glauca* and one of *P. engelmannii* to each dataset, retrieved from the European Nucleotide Archive (ENA) (see *Methods* for details). By repeating the *fastSTRUCTURE* analysis at K=2, we could clearly identify which species ancestry each cluster represented based on the pure individuals admixture coefficients (supplementary fig. S5, Supplementary Material online). Overall, PCA and *fastSTRUCTURE* effectively distinguished the different genomic background of the two spruce species and produced highly concordant results: the distribution of spruce individuals along PC1 reflected the arrangement of the individuals based on the ancestry coefficients estimated with *fastSTRUCTURE* at K=2, with a nearly perfect correlation (0.99) in both datasets (supplementary fig. S6, Supplementary Material online). In PCA, the remaining principal components (PC2, PC3) explained considerably less variation than PC1 in both the elevational and latitudinal transects and separated a few second- and third-degree relatives found within some populations from the rest of the individuals (supplementary figs. S3 and S4, Supplementary Material online). Overall, genomic relatedness levels were low according to our estimation, with only two potential half-sibling relationships (kin ∼0.25) in the elevational dataset and ten in the latitudinal dataset (supplementary fig. S7, Supplementary Material online). Both datasets showed few third-degree relationships (supplementary fig. S7, Supplementary Material online).

The elevational transect exhibited two dominant patterns of covariation between ancestry and environment: 1) ancestry varied from almost pure Engelmann to almost pure white spruce along the broad NW to SE transect across the sampling region (spanning ∼100 km of distance along the valley); and 2) within several locations through the middle of the elevational hybrid zone (particularly locations 4, 5, and 6; Fig. 1c), the proportion of Engelmann ancestry was highest in high elevation N-facing sites (N), lowest in valleys (V), and intermediate on high elevation S-facing sites (Figs. 1c and 1d; Fig. 2; supplementary fig. S8, Supplementary Material online). Within each location, the N, S, and V sites were never separated by more than 10km, and each location was typically separated by ∼5-30 km.

**Fig. 2.**
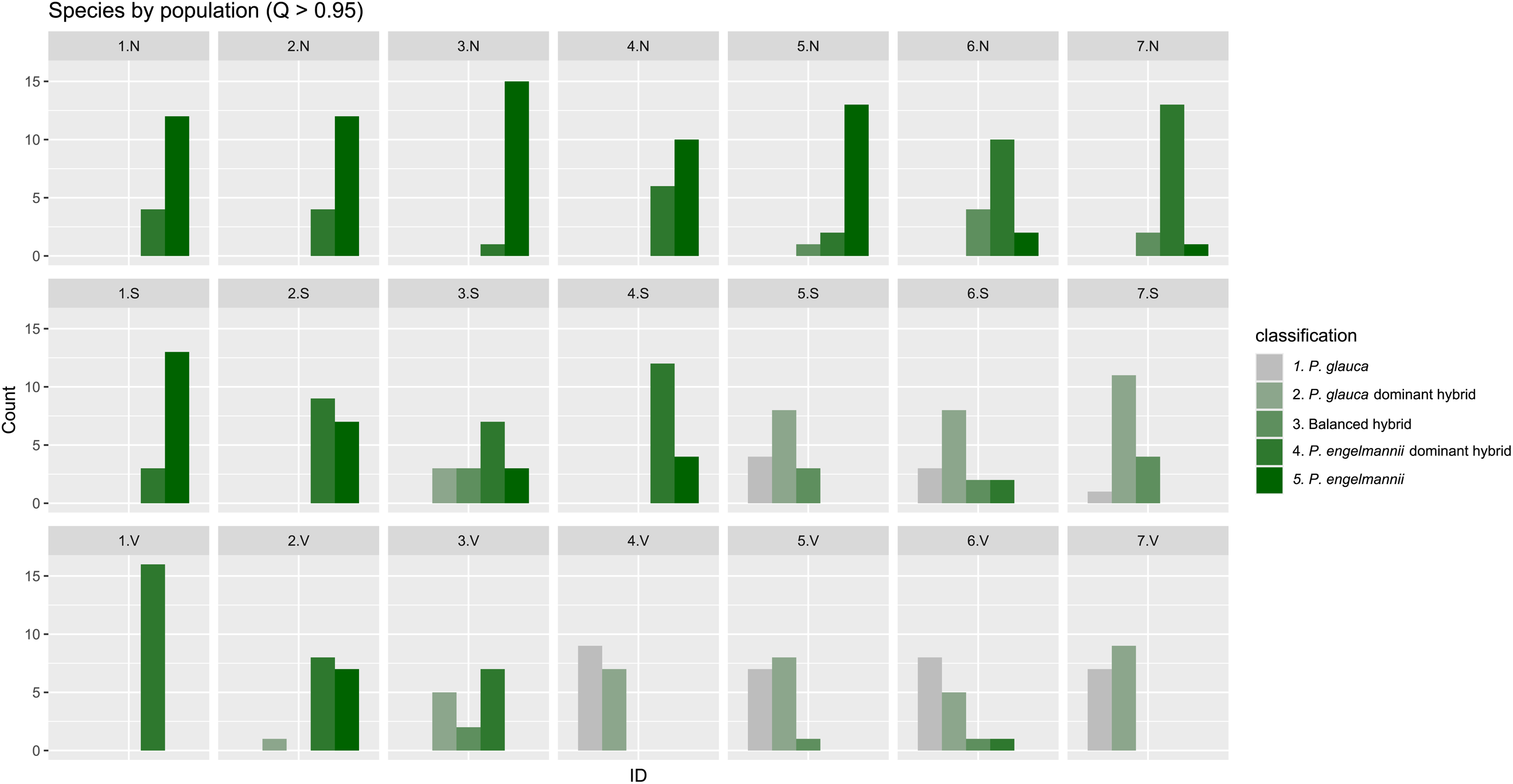
Distribution of individual’s ancestry category across locations and environments in the elevational datasets (N: north-facing slopes, S: south-facing slopes, V: valleys). Individuals were classified by ancestry according to the admixture coefficient computed with *fastSTRUCTURE*, using the following criteria: *P. glauca* (q ≥ 0.95), *P. glauca* dominant hybrid (0.60 ≤ q < 0.95), balanced hybrid (0.40 < q < 0.60), *P. engelmannii* dominant hybrid (0.05 ≤ q ≤ 0.40), *P. engelmannii* (q < 0.05).

Thus, instead of seeing a gradual transition in hybrid ancestry across the elevational transect (with similar ancestry among N, S, and V within locations), we observed substantial fine-grained local adaptation within locations (Fig. 2; supplementary fig. S8, Supplementary Material online). Given that similar changes in ancestry are seen over ∼500km in the latitudinal transect, this represents a comparatively sharp transition in ancestry across micro-environmental variations, maintained over several sampling locations.

For the latitudinal dataset, ancestry distribution aligned well with the geographical ranges of the two species in western Canada (Figs. 1a and 1b) (Rajora and Dancik 2000; De La Torre et al. 2013, 2014; Gagalova et al. 2022). Potentially pure *P. glauca* and *P. glauca*-dominant hybrids were found predominant in the northern and eastern part of the sampling range going into Alberta, while *P. engelmannii* ancestry appeared more prevalent in south-western British Columbia. More balanced species density with varying admixture levels were observed in central and western B.C. and along the southern B.C.-Alberta border, matching with the extent of what is commonly referred to as the western Canada interior spruce hybrid zone (Fig. 1a).

### Population differentiation

We measured the extent of genetic differentiation between populations in each dataset using the FST statistic on a per-SNP basis, implemented in *VCFtools* (<u>Danecek et al. 2011</u>). In the elevational dataset, mean FST between the 21 populations sampled was 0.03 (median = 0.01, sd = 0.05), based on 119,664 SNPs (supplementary fig. S9, Supplementary Material online). In the latitudinal dataset, mean FST between 249 populations was 0.07 (median=0.06, sd=0.07), based on 265,655 SNPs (supplementary fig. S9, Supplementary Material online). For reference, mean FST between *P. glauca* and *P. engelmannii*, estimated by selecting pure species individuals based on the *fastSTRUCTURE* admixture coefficient Q (Q ≥ 0.95), was 0.10 (median = 0.04, sd = 0.15) in the elevational dataset and 0.14 (median = 0.07, sd = 0.17) in the latitudinal dataset (supplementary fig. S9, Supplementary Material online). Most SNPs were polymorphic within both pure species (Q ≥ 0.95) populations, indicating substantial shared variation despite genetic differentiation. Specifically, in the elevational dataset (119,664 SNPs), 96.7 % of SNPs (115,768 SNPs) were polymorphic in *P. glauca* and 99.7 % (119,415 SNPs) in *P. engelmannii*; in the latitudinal dataset (265,655 SNPs), 98.6% of SNPs (262,023 SNPs) were polymorphic in *P. glauca* and 96% (255,500 SNPs) in *P. engelmannii*.

Comparison of the SNPs analysed in each dataset revealed that 31,667 SNPs were shared. This shared set was relatively limited, primarily due to differences in the sequence capture designs used for the two datasets. Moreover, small population sizes and missing data in the latitudinal dataset prevented FST computation for several loci, restricting the cross-dataset FST comparisons to 11,990 SNPs (out of the 31,667 shared SNPs). For this subset of 11,990 SNPs, the correlation of FST values was high (Pearson’s r = 0.85, p < 2.2 × 10⁻¹⁶; Kendall’s tau = 0.48, p < 2.2 × 10⁻¹⁶), with little variation observed among linkage groups (Fig. 3).

**Fig. 3.**
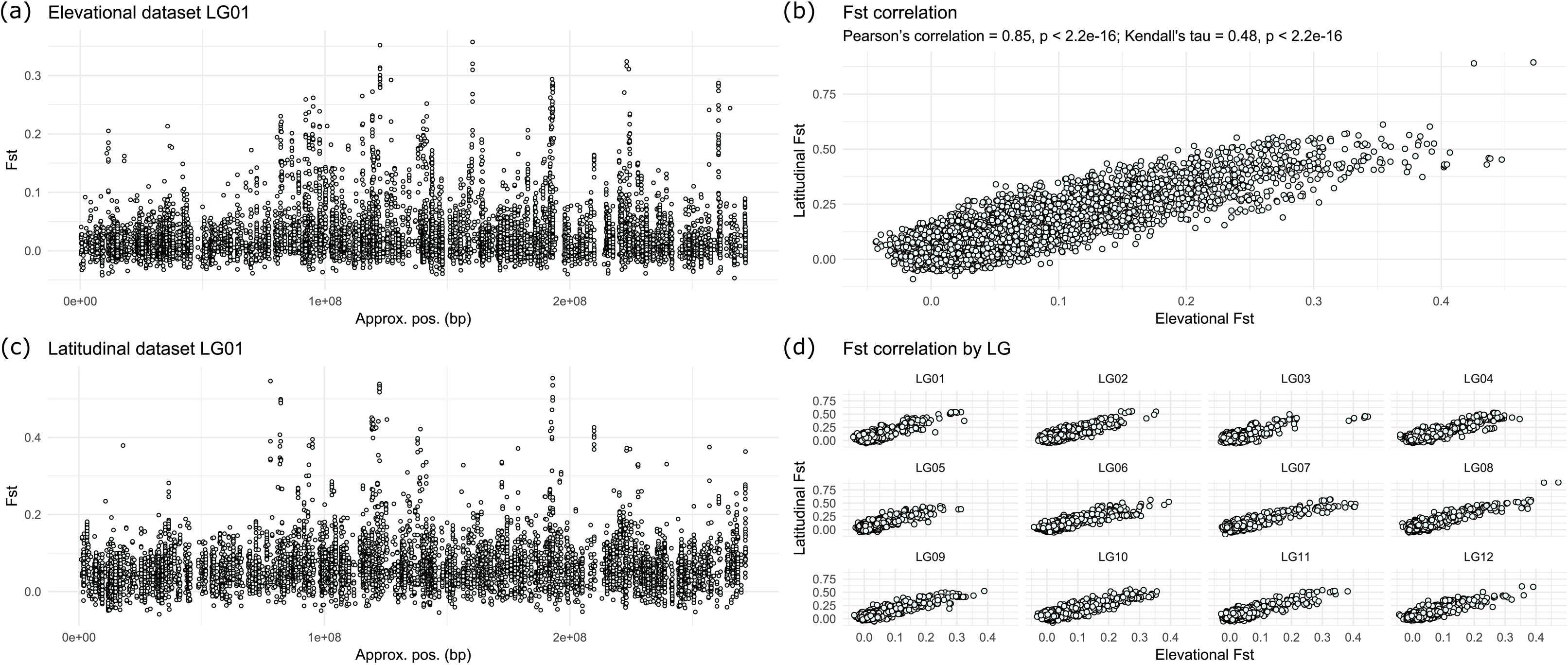
Genomic patterns of among population divergence (FST). Manhattan plots of FST values along LG01 in the (a) elevational and (c) latitudinal dataset. (b) Genome-wide correlation in FST between the shared SNPs of the elevational and latitudinal datasets. (d) Correlation in FST by linkage group (LG). Note that the significance of the p-value in (b) is overestimated as an assumption of the test is violated due to non-independence among linked SNPs.

Overall, the FST landscape between the elevational and latitudinal transects appears highly repeated, with a visually concordant distribution of peaks and valleys in the resulting Manhattan plots (Fig. 3a and c; supplementary fig. S10, Supplementary Material online).

Patterns of nucleotide diversity among parental populations (dxy) were broadly concordant with FST, and we observed relatively low correlation between nucleotide diversity within each parental species (π) and FST, with only a slight tendency for the most extreme FST outliers to have reduced diversity (supplementary figs. S11 and S12, Supplementary Material online). This suggests that background selection reducing π is not the main driver of high FST (as per Cruickshank and Hahn 2014).

### Genome-environment association analysis

To identify the particular combination of environments that covaries most strongly with variation in ancestry, we performed redundancy analysis (RDA) by regressing the individual trees’ hybrid indexes, computed with *fastSTRUCTURE*, against a set of environmental variables considered important in describing the niches of the two parental species. This approach involves a specific type of multiple regression aimed at identifying the linear combination of environmental variables that accounts for the largest portion of variation in the hybrid index, and therefore best explains the introgression patterns that we observed. RDA1 (the first axis of RDA variation i.e. the linear combination of predictors explaining most hybrid index variance) was then used as a predictor in genome-environment association analyses (GEAs). For the elevational dataset, we included six weakly correlated (Pearson’s correlation coefficient < 0.7) environmental variables: aspect (the compass direction that a terrain faces), slope (the steepness of the terrain), air temperature at 2 meters, diffuse horizontal irradiation (DIF), predicted photovoltaic electricity output (PVOUT), and snow melt day (supplementary fig. S13, Supplementary Material online). For the latitudinal dataset, we used the same variables except snow melt day. The latter, being high-resolution data, was not feasible to generate for the vast geographic area covered by the latitudinal dataset.

In the elevational dataset, the first RDA axis (RDA1) explained 73% of the variation in the hybrid index, with snow melt day and PVOUT being the most important variables, followed by air temperature at 2 meters, slope, aspect, and DIF (supplementary fig. S14, Supplementary Material online). In contrast, in the latitudinal dataset RDA1 explained 56% of the total hybrid index variation. The most influential variables were slope and PVOUT, followed by air temperature at 2 meters and DIF (supplementary fig. S14, Supplementary Material online). In this dataset, aspect contributed negligibly to the introgression patterns observed across latitude, in strong contrast with its contribution in the elevational dataset (supplementary fig. S14, Supplementary Material online).

We used the resulting RDA1 axes as predictors in the GEA, conducting separate analyses for the elevational and latitudinal datasets. This analysis should detect candidates driving local adaptation in the parental species, but it also may pick up any regions of the genome that happened to evolve strong differentiation due to drift or intrinsic reproductive isolating mechanisms. The elevational dataset analysis included 119,664 SNPs, while the latitudinal included 265,655 SNPs (MAF > 0.05). For each SNP, we performed a likelihood-ratio test comparing two models: a null model including only the intercept, where the SNP frequency was predicted using only the mean frequency; and a predictor model which used RDA1 as the environmental predictor to infer SNP frequency. The likelihood-ratio test was used to compare the goodness of fit between the predictor model and the intercept-only model. A significantly better fit by the predictor model results in a low p-value, reflecting the predictor’s importance. Conversely, if the null model is supported by the observed data, the likelihoods of the two models would not differ beyond the sampling error, resulting in higher p-values.

Similar to the FST landscape, the GEA landscape between the elevational and latitudinal transects appeared highly repeated, with a visually strikingly concordant distribution of peaks and valleys in the Manhattan plots derived from the two GEAs (Fig. 4a and c; supplementary fig. S15, Supplementary Material online).

**Fig. 4.**
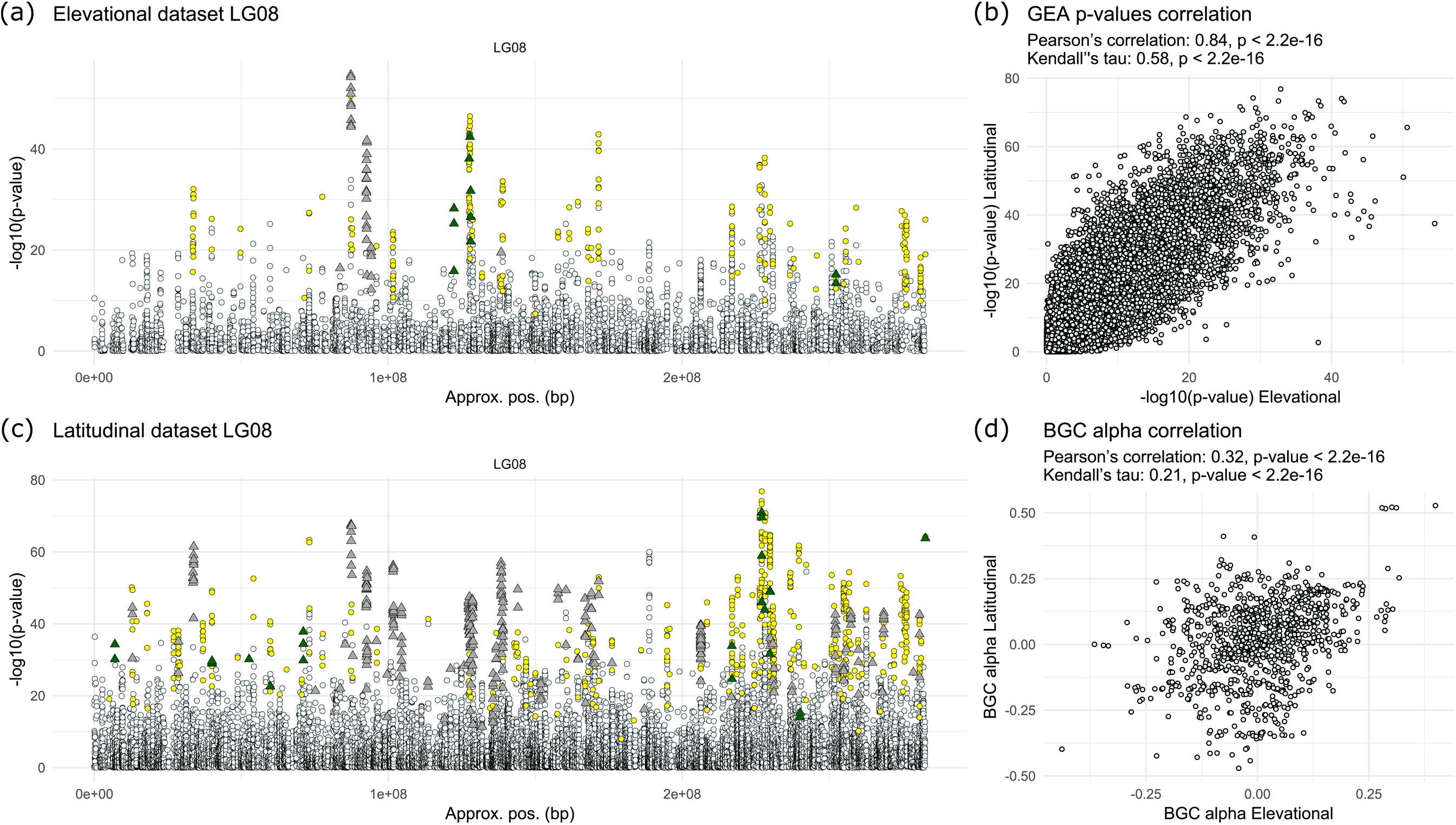
Genomic patterns of Genotype-Environment Association (GEA) and genomic clines. Manhattan plots of GEA p-values along LG08 in the (a) elevational and (c) latitudinal dataset. SNPs showing ancestry excess in the genomic clines (*BGC*) analyses are represented by triangles, coloured as follows: dark green indicates *P. engelmannii* ancestry excess, and grey indicates *P. glauca* ancestry excess. Yellow circles represent SNPs tested in the *BGC* analysis (parental frequency differential > 0.6) but not showing ancestry excess. Small pale-azure circles denote SNPs included only in the GEA analysis but not in the *BGC* analysis. (b) Genome-wide correlation of GEA p-values between the shared SNPs of the elevational and latitudinal datasets. (d) Genome-wide correlation of *BGC* α values between the shared SNPs of the elevational and latitudinal datasets. Note that the significance of p-values in panels (b) and (d) is overestimated as an assumption of the test is violated due to non-independence among linked SNPs.

Between the SNPs analysed in each dataset, 31,667 were shared, exhibiting very high correlation in p-values between the two GEA analyses (Pearson’s correlation: 0.84, p < 2.2 × 10⁻¹⁶; Kendall’s tau = 0.58, p < 2.2 × 10⁻¹⁶), with little variation between linkage groups (Fig.4b; supplementary fig. S16, Supplementary Material online). We also tested whether the inclusion of the snow melt day variable in the elevation environmental set but not in the latitudinal set contributed to our results, but found minimal impact (supplementary Text S1 and supplementary fig. S17, Supplementary Material online).

Concordant with the FST analysis, we assessed correlations between GEA p-values and nucleotide diversity within and between parental species in each dataset. We observed strong correlations with *dxy* and low correlations with nucleotide diversity within parental species, with only a slight tendency for the most extreme GEA outliers to have reduced diversity (supplementary figs. S11 and S12, Supplementary Material online).

The above results do not include any correction for population structure, so we explored the impact of adding a structure correction using PC1 from our PCA analysis (see *Methods* for details). We found that the resulting p-values appear entirely uncorrelated between the elevational and latitudinal analyses with almost no shared outliers, in strong contrast with those of our GEA results without population structure correction (supplementary fig. S18, Supplementary Material online). These observations are consistent with the expected behaviour of the methods: GEA with structure correction should appropriately control the proportion of false positives at the cost of low power to detect true positives, whereas GEA without structure correction should be more sensitive to detecting true positives at the cost of a large number of false positives (Lotterhos 2019; Booker et al. 2023). Both issues are expected to be particularly pronounced when the axis of population structure is confounded with environmental variation driving local adaptation, as is the case in this study. Given that previous studies have shown considerable signatures of local adaptation, it seems likely that GEA without correction (Fig. 4a and c; supplementary fig. S15, Supplementary Material online) is capturing some true signals, along with an unknown proportion of false positives.

### Chromosomal clustering of FST and GEA signatures

To examine chromosomal clustering of signatures of divergence and environmental association, we calculated Moran’s I index of spatial autocorrelation on chromosomal distance. For clustering of FST we used 11,990 shared SNPs for each scaffold and linkage group (see *Methods* for details). Although not statistically significant (scaffolds paired t-test p-value = 0.49, n = 27; LGs paired t-test p-value = 0.22; n = 12), this analysis showed that FST chromosomal autocorrelation tended to be slightly higher in the elevational than in the latitudinal transects when assessed by LG (Elevational scaffolds Moran’s I - Latitudinal scaffolds Moran’s I mean and median: 0.007 and 0.005; Elevational LGs Moran’s I - Latitudinal LGs Moran’s I mean and median: 0.006 and 0.002) (supplementary fig. S19, Supplementary Material online). However, the sample size of only 12 LGs doesn’t allow for much power, and no strong trend was detected in the scaffolds (supplementary fig. S19, Supplementary Material online). We repeated this analysis on the GEA p-values from the 31,667 shared SNPs for each scaffold and linkage group. Levels of GEA p-values chromosomal autocorrelation within scaffolds and LGs were significantly (p < 0.05) higher (scaffolds paired t-test p-value = 7.48 × 10⁻⁵, n = 77; LGs paired t-test p-value = 0.01, n = 12) in the elevational than latitudinal dataset (Elevational scaffolds Moran’s I - Latitudinal scaffolds Moran’s I mean and median: 0.037 and 0.028; Elevational LGs Moran’s I - Latitudinal LGs Moran’s I mean and median: 0.01 and 0.01) (supplementary fig. S20, Supplementary Material online). Additionally, we tested whether the differences in GEA p-values spatial autocorrelation could be driven by the distinct sampling schemes used in the elevational and latitudinal datasets (15-16 individuals per population vs. 2 individuals per population), and found no evidence that variation in populations sampling intensity contributed to these differences (supplementary Text S2 and supplementary fig. S21, Supplementary Material online).

### Genomic clines analysis of aberrant patterns of introgression

Genomic clines analysis is used to infer the movement of ancestry blocks into different genomic backgrounds and to assess patterns of introgression in hybrid zones, by modelling the ancestry probabilities of SNPs as a function of the hybrid index. Here, we aim to identify regions of the genome exhibiting extreme patterns of introgression, which might occur if some loci are more strongly introgressed into one species or the other due to their adaptive effect (hereafter, directional), or if some loci experience unusually steep or gradual transitions across the hybrid index compared to the genome-wide average (hereafter, steepness). Directional introgression (α) occurs when an allele moves from one parental species into the other more often than expected based on genome-wide patterns, which typically happens when a locus experiences uniform directional selection across both species and hybrids (e.g. a classic selective sweep). On the other hand, steepness (β) reflects how rapidly ancestry changes across the hybrid index and can arise from selection favouring heterozygotes (i.e. via complementation of deleterious recessive alleles) or disfavouring heterozygotes (i.e. due to reproductive incompatibilities), resulting in loci that are introgressed more slowly or more rapidly than the genome-wide average (Gompert and Buerkle 2009, 2010, 2011, 2012). We applied genomic clines analyses to assess the patterns of introgression between *P. glauca* and *P. engelmannii* in the western Canada interior spruce hybrid zone. We fit genomic clines using two approaches: a Bayesian genomic clines model implemented in *BGC* (Gompert and Buerkle 2011, 2012) and a logistic regression-based model implemented in *introgress* (Gompert and Buerkle 2010). We used *introgress* to have a source of comparison and verify the consistency of *BGC*, but only results from *BGC* were analysed further.

In order to define parental pure species populations for the genomic clines model, we classified the sampled individuals as either *P. glauca*, *P. engelmannii* or hybrids according to the admixture coefficient computed with *fastSTRUCTURE* (see *Methods* for details). We performed two separate genomic clines analyses, one for the elevational and one for the latitudinal dataset respectively and included in the models only SNPs with an allele frequency differential between parental “pure” species populations greater than 0.6. Limiting the number of SNPs tested in *BGC* was necessary, as using the full SNP set would lead to infeasible run times due to the computational demands of its Bayesian framework.

Therefore, we opted to include only ancestry-informative SNPs (those with substantial parental frequency differentials - 0.6), rather than a random subset that might contain many uninformative loci, as suggested in other studies (Knief et al. 2019; Chaturverdi et al. 2020; Del-Rio et al. 2021; Walsh et al. 2023) and following recommendations in Martin et al. (2021). In our *BGC* analysis, positive α values indicate excess *P. glauca* ancestry compared to the genome-wide average, while negative α values indicate excess *P. engelmannii* ancestry (directional introgression patterns). On the other hand, positive β values indicate a steeper cline and selection against introgression resulting in a faster rate of allelic-frequency transition along the hybrid index, whereas negative β values indicate the reverse.

In the elevational dataset, pre-processing resulted in a set of 3,256 SNPs for genomic clines analysis, which in turn led to the identification of 300 SNPs exhibiting excess ancestry (148 positive α and 152 negative α SNPs). These loci represent SNPs with introgression patterns different from the genome-wide average, with a 95% equal-tail probability interval (ETPI) for the posterior probability distribution of α that did not overlap with zero. No SNP showing unusual rates of allelic-frequency transition along the hybrid index was detected in the elevational dataset genomic clines analysis (β excess) (supplementary figs. S22 and S23, Supplementary Material online).

In the latitudinal dataset, pre-processing resulted in a set of 10,154 SNPs for genomic clines analysis, which in turn led to the identification of 3,845 SNPs with a 95% ETPI for the posterior probability of α that did not overlap 0 (1,906 positive α and 1,939 negative α SNPs), indicating excess ancestry (supplementary figs. S23 and S24, Supplementary Material online). Additionally, the latitudinal dataset analysis pinpointed 120 SNPs exhibiting unusual rates of allele frequency transitions in the hybrid population (52 negative β and 68 positive β SNPs) (supplementary fig. S24, Supplementary Material online). Similarly to SNPs exhibiting excess ancestry (α excess SNPs), these SNPs reported a 95% ETPI for the posterior probability distribution of β that did not overlap with zero (β excess). Among the SNPs that reported α or β excess in the latitudinal dataset analysis, 76 were in common, indicating loci exhibiting both excess ancestry as well as unusual rates of allelic-frequency transitions along the hybrid index gradient (supplementary fig. S24, Supplementary Material online).

Furthermore, we performed two additional *BGC* analyses on subsets of the latitudinal dataset, referred to as Subset 1 and Subset 2, by down sampling the data to a similar number of individuals and parental population composition as in the elevational dataset (supplementary fig. S25, Supplementary Material online). This was done to determine whether the high number of α-excess SNPs observed in the latitudinal dataset, relative to the total number of SNPs tested, was a statistical artifact of their differing sample sizes, or indicated a stronger biological signal in the latitudinal transects compared to the elevational transects. Both downsampled analyses revealed a pattern closely resembling that of the full dataset analysis, identifying 3,258 and 3,268 α-excess SNPs, respectively (supplementary fig. S26, Supplementary Material online), suggesting a stronger biological signal in the latitudinal dataset. Furthermore, the Pearson’s correlation between α values from the full latitudinal dataset analysis and those from the subset analyses was very high, at 0.91 and 0.90. For comparison, α correlation between the two subset analyses was even higher, at 0.97.

Comparing across datasets, the elevational and latitudinal dataset clines analyses shared a total of 982 SNPs. These showed significant correlation of α values between the elevational and latitudinal dataset analyses (Pearson’s correlation: 0.32, p-value < 2.2 × 10⁻¹⁶; Kendall’s tau: 0.21, p-value < 2.2 × 10⁻¹⁶) (Fig. 4d). In more detail, within the 982 shared SNPs, 438 exhibited excess ancestry in the latitudinal dataset analysis and 83 in the elevational - between these, 53 reported excess ancestry across both analyses, with 49 (92%) exhibiting excess ancestry in the same direction of introgression. Overall, *BGC* α values were generally in agreement with the p-value obtained with *introgress* (between *introgress* p-values and *BGC* absolute α values) for both datasets (supplementary fig. S27, Supplementary Material online).

By visually inspecting Manhattan plots derived from our genomic clines analyses, it is apparent that some of the strongest GEA peaks comprising our top candidate for environmental association show asymmetric introgression from one parental species to the other (excess ancestry) (Fig. 4a and c; supplementary fig. S28, Supplementary Material online). This pattern is further confirmed by the relationship between GEA p-values and *BGC* α-values, with numerous α-outlier SNPs exhibiting low GEA p-values (supplementary fig. S23, Supplementary Material online). This could indicate local adaptation associated with the genomic region’s ancestry, even though we cannot totally exclude stable genetic differentiation (i.e. genetic drift). SNPs with these asymmetric patterns tend to be organized into large blocks that all tend to be introgressed in the same direction (Fig. 4a and c; supplementary fig. S28, Supplementary Material online).

### Candidate Genes for Local Adaptation

Given that common garden experiments have shown these species are strongly locally adapted (Jaramillo-Correa et al. 2001; De La Torre et al. 2013; Liepe et al. 2016), regions of the genome harbouring extreme GEA signatures likely harbour many causal drivers of local adaptation. However, given the covariation between structure and environment, it is difficult to know what proportion of these regions are under selection vs. drift. Furthermore, repeated GEA patterns between the latitudinal and elevational transects are non-independent, so it is inappropriate to use statistical tests of whether repeated patterns exceed random expectations. Designating any SNP with a GEA p-value in the bottom 1^st^ percentile as a “GEA outlier” and any SNP with excess ancestry in the genomic clines analysis as an “ancestry outlier”, we present results showing the overlap in outlier SNPs and genes harbouring outliers, both among analyses and across the elevation and latitudinal datasets in the Supplementary Material (supplementary figs. S29 and S30, Supplementary Material online). It is more likely that the patterns of directional introgression are more evolutionarily independent (see Discussion), so overlap of extreme signatures here is more informative. A total of 18 genes were identified as candidates by all four analyses (both GEAs and both cline analyses), exceeding the random expectation (expected 9.7 genes; p = 0.0007) (supplementary Table S1 and supplementary fig. S29, Supplementary Material online).

Although this set of 18 candidate genes requires further study and validation, it comprises several protein families with key roles in plant growth and environmental stress responses (supplementary Text S3 and Table S1, Supplementary Material online), according to the available annotation (Warren et al. 2015; Gagalova et al. 2022). More broadly, the less stringent set of 638 candidates identified by either of our analyses includes several genes previously reported to contribute to adaptation in other genomic studies of North American conifers (supplementary Text S4 and Table S2, Supplementary Material online). Additionally, we examined differences between the candidates identified in the latitudinal and elevational analyses to highlight two distinct groups of SNPs: one putatively associated with elevational but not latitudinal environmental pressures, and the other showing the opposite trend (supplementary Text S3, Supplementary Material online).

## Discussion

Here, we assessed local adaptation, hybridization, and introgression patterns in two regions of the Western Canada interior spruce hybrid zone: a latitudinal gradient spanning ∼1000 km of gradual environmental change and a narrower elevational contact zone spanning ∼100km, including many regions with dramatic changes in environment over short spatial scales of < 10km. We observed dramatic changes in the proportion of white vs. Engelmann ancestry across different elevation/aspect sampling sites within several valley locations (Fig. 1c; Fig. 2). There are two main categories of explanation for the maintenance of highly differentiated populations across such small spatial scales: strong divergent natural selection and/or limited gene flow. Local adaptation between white and Engelmann spruce is well-established (De La Torre et al. 2013, 2014; Hornoy et al. 2015; Yeaman et al. 2016; Capblancq et al. 2022), but it is also possible that phenological barriers prevent mating between sites within locations, given that flowering time is known to respond plastically to temperature cues (Howe et al. 2003; De La Torre et al. 2013, 2014). Further study with transplant experiments and analysis of phenological barriers to reproduction would be needed to resolve this question. It is also unclear whether these repeated patterns of covariation between ancestry and environment represent a single colonization of all N-slopes by Engelmann and all S-slopes by white spruce, or initial colonization of the valleys followed by repeated expansion into the alpine areas by each species.

Our analyses revealed that patterns in the genomic basis of introgression are broadly similar between the elevational and latitudinal transects. We found strong repeatability in our FST and GEA analyses, with high correlation of the resulting FST and GEA p-values and a visually concordant distribution of peaks and valleys in the resulting Manhattan plots (Fig. 3; Fig. 4; supplementary figs. S10 and S15, Supplementary Material online). While some repeatability is expected given the non-independence involved when two hybrid zones arise from the same parental species, the extent to which the patterns observed at broad latitudinal scales (∼1000 km) are repeated at the very narrow spatial scales of the elevational transect (<100 km) is remarkable (Fig. 3; Fig. 4). From these results alone, however, it is not possible to ascertain whether highly divergent regions of the genome were driven by selection or drift, but they do provide a set of candidate regions that likely include many of the causal drivers of local adaptation, which has been well-studied at the phenotypic level (Jaramillo-Correa et al. 2001; De La Torre et al. 2013; Liepe et al. 2016).

The picture becomes clearer with patterns of directional introgression, which are unlikely to be repeated under a pure drift model if the populations in the middle of one transect are evolving independently of those in the other. Our observation of extensive repeatability of directional introgression (Figure 4d; supplementary fig. S28) is therefore more strongly suggestive of a role for natural selection in shaping genomic patterns. However, it is not possible from the data included here to conclusively test whether these populations are evolving independently, as the dominant signal of relatedness is driven by ancestry proportion, leaving little power to assess fine-scale patterns of gene flow. While average dispersal distances in trees tend to be under 1 km per generation (Austerlitz et al. 2004), both pollen and seed are known to travel long distances in conifers, which also exhibit high degrees of outcrossing (O’Connell et al. 2007). It does seem unlikely, however, that gene flow between populations in the middle of the two transects (most of which are separated by > 100km; Fig. 1 and supplementary fig. S25, Supplementary Material online) could be high enough to entrain their allele frequencies to drive repeated patterns, while also being limited enough to allow the dramatic divergence we see within several locations in the elevation transect (< 10km). Although further study is required to conclusively assess this evidence, the most likely explanation for observed repeated patterns of directional introgression is natural selection.

Our analysis also highlights that correcting for population structure in GEAs can be counterproductive, particularly when there is strong collinearity between the environmental gradient and the axis of introgression (supplementary fig. S18, Supplementary Material online). Methods that include structure correction may fail to detect true adaptive loci in such cases, as previously suggested (Lotterhos 2019; Booker et al. 2023); however, not accounting for population structure is very likely to lead to elevated false-positive rates, making this a particularly challenging problem without a clear solution. When the goal is to identify a long list of candidates for future study, maximizing sensitivity while allowing for inclusion of false positives is preferable to using structure correction to strictly control the false positive rate and fail to detect genomic regions driving selection.

In our analysis of the genomic distribution of regions of high differentiation between species, chromosomal autocorrelation of the resulting FST and GEA p-values shows that differences in the length of chromosomal tracts of introgression between latitudinal and elevational datasets are broadly consistent with expectations based on the theory of hybridization (Barton 1983; Sedghifar et al. 2016). In the elevational transect, there is less opportunity (fewer generations) for recombination across the hybrid zone, because of the abrupt transition across shorter spatial distances (Shchur et al. 2020), which results in larger blocks of ancestry around the loci with strong patterns of differentiation and association with environment (supplementary figs. S19 and S20, Supplementary Material online). Given that common garden experiments show substantial evidence for local adaptation between the parental species, some of these regions may be shaped by strong divergent selection, which would further contribute to these patterns (Shchur et al. 2020). Unfortunately, the power of this analysis was limited by the restricted overlap among genomic regions targeted by the different sequence capture panels used for the two datasets. In addition, the fragmented state of the white spruce genome assembly and the limited resolution of the linkage map required us to rely on the rank order of SNPs within scaffolds anchored along the LGs of our super-scaffold assembly rather than on physical or recombinational (cM) distances, reducing spatial precision. Therefore, further exploration of this hypothesis is warranted once a more contiguous reference genome becomes available.

Our results also echo previous work suggesting that the interior spruce hybrid zone is maintained by a bounded hybrid superiority model, where hybrids exhibit higher fitness than parental species in intermediate environments (De La Torre et al. 2013, 2014). In our case, this may be represented by high south-facing slopes in the middle of the elevational dataset (locations 3, 4, 5, and 6; Fig. 1c), which combine features of the longer, dry summers that favour *P. glauca* and the higher precipitation that favours *P. engelmannii*. Consistent with this, south-facing slopes reported a higher frequency of balanced hybrids (hybrids close to 50:50 ratio), a balanced distribution of *P. glauca* dominant and *P. engelmannii* dominant hybrids as well as lower frequencies of putative pure *P. glauca* individuals compared to valleys, and putative pure *P. engelmannii* individuals compared to north-facing slopes (Fig. 2; supplementary fig. S8, Supplementary Material online). The intermediate habitats provided by south-facing slopes may allow hybrids to outperform both parental species by combining traits such as faster growth with earlier bud setting and greater tolerance to snow load (De La Torre et al. 2013, 2014).

Loci that repeatedly exhibit high differentiation, strong evidence of adaptation, and consistent directional introgression constitute the strongest candidates for local adaptation and adaptive introgression (Rieseberg et al. 1999; Buerkle and Lexer 2008; Gompert and Buerkle 2009; Bierne et al. 2013; Simon et al. 2020; Westram et al. 2021; Langdon et al. 2022). The 18 strongest candidate adaptive loci identified here included several genes previously reported for their central role in plant growth and environmental adaptation (supplementary Text S3 and supplementary Table S1, Supplementary Material online). Furthermore, several of the genes present within our broader set of candidates (638 genes identified by either of our analyses), were also reported to contribute to adaptation in similar studies of local adaptation in other North American conifers (Hornoy et al. 2015; Yeaman et al. 2016; Depardieu et al. 2021; Capblancq et al. 2022) (supplementary Text S4 and Table S2, Supplementary Material online). Additionally, differences in candidate loci between the elevational and latitudinal transects revealed several genes potentially involved in the response to environmental factors specific to latitude such as photoperiod and seasonality, as well as others potentially involved in regulating factors more specific to elevational changes, such as oxygen levels, UV radiation, temperature, snow and precipitation levels (supplementary Text S3, Supplementary Material online). However, we note that our candidates for local adaptation require further validation, and it is likely that this broader set of 638 genes includes false positives due to the difficulty in completely disentangling environmentally driven variation from neutral population structure effects. One way we tried to address this was through the analysis of genomic clines, as directional introgression replicated across hybrid zones characterized by different environmental contrasts would not be expected under a neutral drift model (Rieseberg et al. 1999; Buerkle and Lexer 2008; Gompert and Buerkle 2009; Bierne et al. 2013; Simon et al. 2020; Westram et al. 2021; Langdon et al. 2022). This is what we observed: directional introgression occurred largely in the same genomic regions and often in the same direction between the elevational and latitudinal transects, as reflected by the significant correlation of *BGC* alpha scores and overall highly concordant genome-wide patterns of directional introgression (Fig. 4; supplementary fig. S28, Supplementary Material online). This supports the idea that the 18 strongest candidate genes identified by all four analyses form a robust set of adaptive loci (supplementary Table S1 and fig. S29, Supplementary Material online), unlikely to be generated by drift.

## Materials and Methods

### Super-scaffold assembly construction

We built a super-scaffold assembly of the WS77111v2 white spruce (*P. glauca*) genome to use in subsequent analyses, by conservatively anchoring scaffolds to the latest publicly available *P. glauca* linkage map including 14,727 cDNAs (Gagalova et al. 2022). For this process, we retrieved the sequences of an exhaustive set of 27,720 cDNAs, stemming from the catalogue of spruce expressed genes assembled by Rigault et al. (2011), which we filtered to retain only the sequences of the 14,727 cDNAs present in the *P. glauca* linkage map (Gagalova et al. 2022). We then used *GMAP* (Wu et al. 2016) to align the retained cDNAs sequences to the scaffolds of the WS77111v2 white spruce reference genome, and the output was processed according to the following criteria: mapped cDNAs with coverage above 50% and sequence identity above 90% were retained, while anything else was discarded to ensure retention of only high confidence hits. Then, for each cDNA that satisfied these mapping criteria, we extracted its best scaffold match based on the highest sequence identity. In case a cDNA best mapping scaffold could not be determined as it mapped equally well to multiple locations in the genome, it was discarded. This conservative approach ensured the removal of difficult to resolve multi-mapped cDNAs. Finally, the scaffolds with mapped cDNAs were ordered along the 12 spruce linkage groups according to the cDNAs centimorgan (cM) positions in the linkage map.

### Sampling and sequencing

We collected cambium tissue from 331 mature spruce individuals, including *P. glauca*, *P. engelmannii* and their hybrids, from seven locations along the Trans-Canada highway going through Canmore, Banff and Yoho in the summer of 2022 (Fig. 1c, elevational transect).

Each location included three distinct populations subjected to contrasting environmental pressures: a north-facing high-altitude population, a south-facing high-altitude population, and a valley population (Fig. 1c). Within each population, we sampled between 14 and 16 individuals (supplementary fig. S1, Supplementary Material online). Cambium tissue samples were sent to Rapid Genomics LLC (Gainesville, Florida) for DNA extraction, library preparation, and sequencing using a 10 Mb target approach.

We assembled a second dataset by downloading raw exome capture data for 575 mature spruce individuals from across 249 populations distributed throughout western Canada (Fig. 1a, latitudinal transects), derived from previous studies (Suren et al. 2016; Yeaman et al. 2016). This data is publicly available in the Short Read Archive (SRP071805; PRJNA251573).

### SNP calling

We applied a uniform SNP calling pipeline to both datasets for consistency. First, *fastp* (Chen et al. 2018) was used to remove sequencing adapters and for trimming raw paired end FASTQ files, using default settings. Processed reads were then mapped to the latest publicly available *P. glauca* reference genome (WS77111v2, https://www.bcgsc.ca/downloads/btl/Spruce/) (Warren et al. 2015) using *BWA-mem* (Li and Durbin 2009). Following mapping, we used *samtools* (Li et al. 2009) to convert the alignment files generated from SAM to sorted BAM formats and discarded any alignment with mapping quality below 10. *Picard* tools (available at https://broadinstitute.github.io/picard/) was then used to remove potential PCR duplicates and to assign read groups, followed by indel realignment with *GATK 3.8* (DePristo et al. 2011). Finally, SNP calling was performed using *BCFtools mpileup* and *BCFtools call* (Danecek et al. 2021), based only on alignments with a minimum mapping quality of 5 (−q 5). We filtered raw VCF files with *VCFtools* (Danecek et al. 2011) to retain only biallelic SNPs with the following characteristics: genotyped in at least 70% of the individuals, quality above 30 (--minQ 30), individuals genotype quality above 20 (--minGQ 20), minimum read depth above 5 (--minDP 5) and minor allele frequency greater than 0.05. Additionally, we filtered the VCF files of both datasets (latitudinal and elevational) to retain only the SNPs located on the scaffolds that we were able to anchor to the available linkage map (Gagalova et al. 2022) in our super-scaffold assembly construction.

### Population structure assessment

We used *fastSTRUCTURE* (Raj et al. 2014) to assess population structure in both datasets, using a simple prior and testing Ks ranging from 1 to 10, and then selected the K model that maximizes the log-marginal likelihood lower bound (LLBO) of the data (K*ɛ). To determine which species each *fastSTRUCTURE* cluster represented, we added whole genome sequencing data of two pure species individuals cultivated at the arboretum of the Horsholm University of Copenhagen (Denmark) and repeated the *fastSTRUCTURE* inference. This data was retrieved from ENA under accessions SAMEA51442168 (*P. glauca*) and ERX1862345 (*P. engelmannii*). Furthermore, we classified individuals as either pure *P. glauca*, pure *P. engelmannii* or hybrid according to the value of the admixture coefficient (Q) computed with *fastSTRUCTURE* at the chosen model complexity (K=2): q ≥ 0.95 for either pure species, while individuals with 0.05 < q < 0.95 were classified as hybrids. Population structure was further assessed using principal component analysis (PCA), implemented in *Plink* version 2.0 (Chang et al. 2015). Marker-based realized genomic relatedness was computed between all possible pairs of individuals according to the formula by VanRaden (2008), implemented in the kin function of the R package *synbreed* (Wimmer et al. 2012).

### Population differentiation

Weir and Cockerham FST (Weir and Cockerham 1984) was calculated on a per-SNP basis among the 21 populations of the elevational dataset and the 249 populations of the latitudinal dataset using the *VCFtools* function *weir-fst-pop* (Danecek et al. 2011), which reports the average FST of all pairwise population comparisons for each SNP. Additionally, FST between *P. glauca* and *P. engelmannii* was calculated within each dataset, selecting pure species individuals based on the admixture coefficient computed with *fastSTRUCTURE* (q ≥ 0.95). Using this assignment, nucleotide diversity π (Nei and Li 1979) was also computed within each parental species (π) and between parental species (*dxy*) using piawka (Scott et al. 2025).

Moran’s I indices of spatial autocorrelation based on FST scores were computed using the R package *moranfast* (https://github.com/mcooper/moranfast), using SNP position along both scaffold and linkage group. To measure clustering of FST values along the genome, we calculated Moran’s I using SNP index positions, assigning each SNP a sequential number (1, 2, 3, …) based on its genomic order in our super-scaffold assembly. This approach treats SNPs as equally spaced and captures how long similar values persist along scaffolds and, more broadly, along linkage groups. It is intended to reflect the blockiness of the signal, which may be related to haplotype structure. We used ordinal position of SNPs rather than genomic positions because base-pair distances between scaffolds are unknown, due to the highly fragmented reference genome. Additionally, estimation of Moran’s I using genomic position would be lower if a given set of SNPs were spread across long physical distances. As it is expected that SNP density would be lower in regions with slow LD decay and longer LD tracts, using genomic distances would erroneously bias the measurement downwards in these regions. Ideally, this analysis would use recombination units (cM), but the available linkage map lacks the resolution necessary to permit this.

### Genome environment association analysis

We downloaded elevation data at 90-meter resolution from the NASA Shuttle Radar Topography Mission (SRTM) (https://doi.org/10.5069/G9445JDF) and used it to compute slope and aspect using the *terrain* function of the *raster* R package (Hijmans and Etten 2012). Additionally, we downloaded seven solar variables from the Global Solar Atlas (https://globalsolaratlas.info): photovoltaic electricity output (PVOUT), global horizontal irradiation (GHI), diffuse horizontal irradiation (DIF), direct normal irradiation (DNI), optimum tilt-angle (OPTA), air temperature at 2 meters (TEMP) and global tilted irradiation (GTI).

After extracting measurements for this set of 11 variables for all our samples, we selected environmental variables with Pearson’s correlation coefficient < 0.7 in order to avoid overfitting. The following five variables were retained: slope, aspect, DIF, PVOUT and TEMP. To describe the selected solar variables, DIF measures the scattered sunlight that reaches a horizontal surface, excluding direct solar beams. It serves as an indicator of indirect light availability, which can influence plant adaptation in shaded or cloudy environments. In contrast, PVOUT represents the potential electricity generated by solar panels and acts as a proxy for total solar irradiance, integrating both direct and indirect light. As a result, PVOUT provides a measure of the total environmental solar energy available at a given location.

Moreover, we generated high-resolution (30 meter) snow-cover data for the region covered by the elevational dataset using the R package *SnowWarp* (Berman et al. 2018; Laurin et al. 2022), which fuses 30-meter Landsat optical imagery with 500-meter MODIS (Moderate Resolution Imaging Spectroradiometer) snow cover data using the Google Earth Engine platform (Gorelick et al. 2017). Daily fractional snow-covered area (fSCA) was computed for the time period spanning 2000 to 2021 and was used to derive the yearly date of snowmelt for each 30-meter pixel, which was calculated as the first date in the second half of a snow year where at least 15 of the previous 30 days were snow-free (Berman et al. 2019). Finally, the mean snowmelt date was obtained by averaging snowmelt dates from 2000 to 2021 for each of the 21 populations in the elevational dataset. For each population, before calculating the yearly average snowmelt date, we applied a spatial smoothing step by averaging the snowmelt dates of each pixel together with its neighbouring pixels, rather than using the exact value of each population location pixel alone. This approach aimed to capture a more representative local signal around each population. It was not feasible to generate this high-resolution data for the vast area covered by the latitudinal dataset, therefore we only included snowmelt day for the elevational dataset GEA, as we deemed it particularly well suited to discriminate between north-facing slopes, south-facing slopes and valleys (supplementary fig. S13, Supplementary Material online).

We then used redundancy analysis (RDA) (Capblancq and Forester 2021) in the R package *vegan* to identify the linear combination of the retained environmental variables that explains the greatest variation in the hybrid index. This was performed independently for the elevational (six environmental variables including snow melt) and latitudinal dataset (five environmental variables). Finally, we used R to perform the genotype environment association analyses. These involved a likelihood-ratio test for each SNP, comparing a null intercept-only model to a predictor model which used the first axis of variation (RDA1) from RDA to predict SNP frequency. In a likelihood-ratio test, a low p-value is obtained when a significantly better fit is obtained with the predictor model and it reflects the importance of the predictor: the smaller the p-value, the more important the predictor.

Following the same approach used in the FST analysis, we computed Moran’s I indexes of spatial autocorrelation based on GEA p-values using the R package *moranfast* (https://github.com/mcooper/moranfast). To provide a comparison, we repeated the log-likelihood ratio test-based GEA analysis, this time incorporating structure correction using the first principal component (PC1) from our PCA. In this analysis, we compared a null model, where SNP frequency was predicted by PC1 alone, to a predictor model that included both PC1 and the first axis of variation from RDA (RDA1 * PC1) to predict SNP frequency.

### Genomic clines analysis

We performed a genomic clines analysis to identify SNP loci exhibiting excess ancestry from either parental species relative to the average genomic background, as well as to identify SNP loci with unusual rates of allelic-frequency transitions along the hybrid index. For this analysis we used a Bayesian genomic clines model implemented in *BGC* (Gompert and Buerkle 2011, 2012) and a logistic regression-based model implemented in *introgress* (Gompert and Buerkle 2010). In *introgress*, we used both permutation and parametric tests to compute p-values. We restricted each analysis to SNP loci with a frequency differential between pure *P. glauca* and pure *P. engelmannii* of at least 0.6. Parental pure species and hybrid populations were defined for both datasets according to the admixture coefficient computed with *fastSTRUCTURE* (Q ≥ 0.95). This resulted in the following setups: 82 pure *P. engelmannii*, 163 pure *P. glauca*, and 330 admixed individuals in the latitudinal dataset, and 99 pure *P. engelmannii*, 40 pure *P. glauca*, and 192 admixed individuals in the elevational dataset. Furthermore, we conducted two additional *BGC* analyses on the latitudinal dataset, using the same SNPs but down sampling the data to obtain a similar number of individuals and parental population composition as in the elevational dataset, to allow for better comparison. Subset 1 included 70 pure *P. engelmannii*, 86 pure *P. glauca*, and 200 admixed individuals, while Subset 2 included 70 pure *P. engelmannii*, 45 pure *P. glauca*, and 194 admixed individuals. In our subsampling strategy, we prioritized removing individuals from the latitudinal dataset which were located near the area sampled in the elevational dataset, as well as populations from western British Columbia (supplementary fig. S25, Supplementary Material online), to avoid possible signals of admixture with Sitka spruce (Hamilton et al. 2012). In *BGC* we used 100,000 MCMC generations with 50,000 burn-in and retained every 24^th^ iteration. After assessing convergence, SNPs showing excess ancestry or unusual rates of allelic-frequency transitions along the hybrid population were identified as those for which the 99% equal-tail probability interval (ETPI) for the posterior probability distribution of the clines parameters α or β did not overlap zero. In *introgress*, we used default settings and ran both the parametric and permutation tests (Gompert and Buerkle 2010). Both tools were run within the *ClineHelpR* R package (Martin et al. 2021).

### Identification of candidate genes and overlap with other studies

We used bedtools (Quinlan and Hall 2010) to identify the genes intersected by our top candidate SNPs from the GEA and genomic clines analyses. We expanded the gene sequences by adding 5000 bp flanks on either side, to include possible regulatory elements.

We used the gene models provided for the WS77111v2 *P. glauca* reference genome (https://www.bcgsc.ca/downloads/btl/Spruce/). We used *Orthofinder2* (Emms and Kelly 2019) on each species proteome to identify orthology between white and red spruce, and between white spruce and lodgepole pine. For white spruce (*P. glauca*), we used the proteome for WS77111v2, available at https://www.bcgsc.ca/downloads/btl/Spruce/Pglauca_WS77111/genome_annotation/WS77111v2/. As red spruce was mapped to Norway spruce (*P. abies*) in the study by Capblancq et al. (2022), we used the proteome available for *P. abies* version 1 reference genome (https://treegenesdb.org/org/Picea-abies). For lodgepole pine (*Pinus contorta*), we first retrieved the entire transcriptome analysed in Yeaman et al. (2016) and used TransDecoder (https://github.com/TransDecoder/TransDecoder) to identify the longest open reading frames for each transcript and then translate them into protein sequences. Finally, for each species proteome we selected a single primary transcript per gene according to the longest isoform, and then *Orthofinder2* was run using default settings and independently for white spruce versus Norway spruce and white spruce versus lodgepole pine.

## Data Availability

Sequence data are deposited in the Short Read Archive under BioProject IDs PRJNA251573 (latitudinal dataset) and PRJNA1294281 (elevational dataset). Additional resources are available on Zenodo (https://doi.org/10.5281/zenodo.17474986), including the order of anchored scaffolds in the linkage map used to arrange the VCFs according to our super-scaffold assembly, VCF files for the elevational and latitudinal datasets, and metadata with individual coordinates and populations information.

## Acknowledgements

We would like to thank Sally Aitken for many conversations that helped inspire this work, Ian MacLachlan, Jason Mogilefsky, Vashti Dunham, and Tim McCready for help obtaining funding and overall planning of the project, and staff at the Canadian and Alberta parks systems for help with securing permits. Thanks also to Michael Williamson for help brainstorming the methods, to members of the Yeaman lab for feedback, and to Janek’s brother, Grant Scully, and Sai Yeaman for help with field work. This project was funded by an NSERC Alliance and FRIAA funds contributed by Weyerhauser and West Fraser Timber, with computational support from the Digital Research Alliance of Canada.

## Author contributions

GN analysed the data and wrote the manuscript. JS performed sampling and collected cambium for DNA extraction. AS helped assess snow cover data. BB generated snow cover data. ML and NI contributed to the development of genomic resources. SY conceived the project and secured funding, designed sampling and sequencing methods, provided overall supervision, and helped with writing the manuscript.

