## Supplementary Material for "Replicated hybrid zones reveal genomic patterns of local adaptation and introgression in spruce"

### Supplementary Text

##### Text S1. RDA and GEA without the snow melt day variable

To test whether the difference in environmental variables affected our elevational-latitudinal comparison, we repeated the RDA and GEA in the elevational dataset after removing snow melt day, so that both datasets used exactly the same environmental variables. Without snow melt day, RDA1 in the elevational dataset explained ~63% of the variance in hybrid index (vs. 73% when snow melt was included) but was strongly correlated with the original RDA1 axis including snow melt day (Pearson = 0.93; Kendall's tau = 0.76) (supplementary fig. S17, Supplementary Material online). Concordantly, the resulting GEA reported high correlation with the original elevational GEA including snow melt day (Pearson's = 0.99; Kendall's tau = 0.86), and with the latitudinal GEA (Pearson's = 0.83; Kendall's tau = 0.56). This confirms that including snowmelt day in the elevational dataset did not substantially alter the elevational-latitudinal comparison but it only slightly improved the RDA explanatory power in the elevational dataset. This is likely because the variation explained by snowmelt day is still largely captured by PVOUT (Pearson's = -0.49), air temperature at 2 m (Pearson's = -0.69) and slope (Pearson's = 0.44).

##### Text S2. Evaluating the influence of sampling intensity on GEA p-values spatial autocorrelation

To test whether the differences in GEA p-value spatial autocorrelation could be driven by the distinct sampling strategies used between the elevational and latitudinal datasets (2 individuals per population vs. 15-16 individuals per population), we subsampled the elevational dataset 1,000 times, retaining only 2 random individuals per population for each iteration. We repeated the GEA analysis for each subsampled elevational dataset and subsequently recalculated Moran's I based on the resulting p-values. The 1,000 GEAs from the subsampled elevational datasets showed mean Pearson correlations of 0.31 (p-values) and 0.72 (-log10 p-values) with the fully sampled elevational GEA. Moreover, recalculating the difference in Moran's I between each subsampled elevational GEA and the latitudinal GEA across scaffolds and linkage groups (LGs) revealed that the patterns observed with the original dataset strongly persisted. Mean and median differences between the subsampled elevational datasets Moran's I and the latitudinal Moran's I were almost always positive with rare exceptions, whether this was assessed by scaffolds or by LGs (supplementary fig. S21, Supplementary Material online). This confirms that the higher autocorrelation in GEA p-values that we observed in the elevational dataset is unlikely due to differences in sampling strategy with the latitudinal dataset.

##### Text S3. Functional roles of candidate adaptive genes

The more stringent set of 18 candidate genes identified across all four analyses (elevational GEA, elevational clines, latitudinal GEA, latitudinal clines) comprised some protein families with key roles in plant growth and environmental stress response. These included cytochrome P450, a superfamily of enzymes linked to both biotic and abiotic stress responses as well as developmental processes in plants (Xu et al. 2015), and recently identified as a target of repeated global selection in multiple plant species (Nocchi et al. 2024). MYB transcription factors, which are an extensively studied family of plant proteins involved in development, metabolism and responses to biotic and abiotic stresses (Biswas et al. 2023; Roy 2016). The HhH-GPD DNA glycosylase superfamily, which includes proteins involved in DNA repair caused by oxidative stress (Nota et al. 2015). And a family of UV-B-induced proteins, though not yet fully characterized, was also present within this set of 18 candidate genes.

Additionally, we examined the different candidates between the elevational and latitudinal analyses to identify genes potentially involved in environment-specific stresses. We identified three genes that appear to be involved in adaptation to elevational but not latitudinal, environmental pressures (supplementary fig. S29, Supplementary Material online). These genes belonged to the no apical meristem (NAM) and F-box protein families. NAM proteins are involved in responses to various abiotic factors and play key roles in plant development, particularly in leaf formation, shaping morphology, patterning, and size, which in turn influence critical processes such as transpiration, light absorption, and thermal regulation (Hasson et al. 2010; Hu et al. 2010; Cheng et al. 2012; Wang et al. 2022). These proteins have also been previously identified as adaptive candidates in common garden studies of local adaptation in interior spruce in British Columbia (De La Torre et al. 2014). F-box proteins are similarly involved in a wide range of responses to abiotic factors, including drought, temperature, and in particular, UV radiation (Lechner et al. 2006; Zhang et al. 2013, 2014; An et al. 2016; Wang et al. 2024). These types of abiotic factors tend to vary more abruptly with elevation than with latitude, as environmental transitions along latitude are mostly related to photoperiod and seasonality and generally shift more gradually. While this interpretation is consistent with known functional roles, we note that further validation is needed to confirm the involvement of these genes in adaptation specific to elevational gradients.

Conversely, we identified 110 genes (supplementary fig. S29, Supplementary Material online) that appear to be involved in adaptation to latitudinal, but not elevational, factors. While exploring each candidate goes beyond the scope of this work and this set requires further validation, it is worth highlighting the presence of several proteins involved in photoperiodic flowering regulation. These included the phosphatidylethanolamine-binding protein (PEBP) family (Faure et al. 2007), COL genes (Chen et al. 2021), PP2C phosphatases (Zeng et al. 2022), TCP transcription factors (Zhang et al. 2024), AP2/ERF transcription factors (Matías-Hernández et al. 2014), SPL transcription factors (Jung et al. 2011), 14-3-3 proteins (Paul et al. 2008) and the trehalose-phosphatase family (Wahl et al. 2013).

**Text S4. Comparison with adaptive loci identified in similar studies of local adaptation in conifers**

The study by Hornoy et al. (2015) identified 131 *P. glauca* genes with signals of selection along temperature and precipitation gradients in eastern Canada, of which 108 were included in our tested gene set. Between these 108 genes and the broader less stringent set of genes identified as candidates in either of our analyses (638 genes), 7 were shared (supplementary Table S2, Supplementary Material online). This study tested about a quarter of the transcriptome (7,819), but since it is not clear exactly which genes were tested, a hypergeometric expectation cannot be calculated. Nonetheless, it is unlikely that the identified overlap (7 genes) represents a strong enrichment against random expectation. The set of seven common candidate genes included some notable protein families, such as the AP2/ERF transcription factor family, which comprises essential regulators in many biological processes like plant growth, fruit/flower development, photoperiodic flowering and biotic and abiotic stress responses (Matías-Hernández et al. 2014; Jiang et al. 2022; Jiang et al. 2024). This family has also been identified as a target of repeated global selection across several plant species in a recent comparative genomics study (Nocchi et al. 2024). RieskeFeS proteins which are involved in light capture have been shown to improve the efficiency of photosynthesis and to stimulate growth in low light conditions (Simkin et al. 2017). And the PAP2 superfamily, which includes proteins that when overexpressed have been shown to increase photosynthetic rate and enhance growth (Cai et al. 2022).

The study on red spruce (*P. rubens* Sarg.) in north-eastern North America by Capblancq et al. (2022) identified 125 genes with signals of local adaptation along climatic gradients, of which 76 had orthology with *P. glauca* according to our orthology assignment. Between these 76 genes and the 638 genes identified in our study (of which 541 with orthology), 7 were in common (supplementary Table S2, Supplementary Material online). In total our analysis and their analysis tested 4,897 common orthogroups. Although the observed overlap of 7 genes does not significantly deviate from the statistical expectation based on the hypergeometric distribution (expected overlap = 8.39, observed overlap = 7, p-value = 0.75), this set of common candidates included some notable proteins, such as a longevity assurance gene (LAG1) which helps plants resist toxins and survive ER stress (Brandwagt et al. 2000), and a flowering time control gene (FPA). FPA has been shown to regulate flowering via a temperature-driven but day-length independent mechanism, suggesting its potential role in the phenological differentiation of populations experiencing identical photoperiod regimes (i.e. same latitude, different elevation) (Schomburg et al. 2001; Capblancq et al. 2022).

The study by Yeaman et al. (2016) on convergent evolution identified 258 lodgepole pine candidate genes for local adaptation along geographic and environmental gradients in north-western North America, of which 136 had orthology relationships that could be identified with spruce. Between these 136 genes and the 638 genes identified in our study (of which 489 had orthology with pine), 12 were shared (supplementary Table S2, Supplementary Material online). In total these two analyses tested 4,220 common orthogroups. This set of 12 genes, despite not significantly differing from the number of expected overlap based on the hypergeometric distribution (expected overlap = 15.75, observed = 12, p-value = 0.87), still included proteins documented to contribute to adaptation, such as TCP transcription factors which mediate plant growth and development (Yu et al. 2022), AP2/ERF transcription factors (also a candidate protein family in Hornoy et al. 2015), which are involved in growth, flowering regulation and defence responses to a variety of stresses (Matías-Hernández et al. 2014; Jiang et al. 2022; Jiang et al. 2024), and four zinc finger proteins (ZFPs). These latter proteins constitute the largest E3 ubiquitin ligase family and play important roles in plant growth, development, and response to abiotic stresses such as drought, salt, temperature, reactive oxygen species and harmful metals (Han et al. 2022). Consistent with this, the study by Depardieu et al. (2021) on drought adaptation *in P. glauca* in Quebec (Canada) found that ZFPs were among the most represented gene families in their set of adaptive candidates. More specifically, Depardieu et al. (2021) identified eight high-confidence genes associated with drought tolerance in white spruce populations, including GQ03707_G19 (DB47_00099276), a zinc finger transcription factor that was also detected in our study and in the lodgepole pine study by Yeaman et al. (2016). This gene was correlated with the summer soil moisture index in their GEA, was linked to annual fluctuations in wood density in their genotype-phenotype association (GPA) and was also shown to be upregulated under drought conditions in their gene expression analysis (Depardieu et al. 2021). Furthermore, a Kelch-repeat-containing protein was also present in our set of 18 candidate genes identified by all analyses. Kelch repeat proteins (and other Kelch-repeat subunits) act as substrate-recruiting components of E3 ubiquitin-ligases in plants (Wu et al. 2025), and, notably, E3 ligases were recently revealed as key drivers of global adaptation across several plant species (Nocchi et al., 2024).

### Figures & Tables

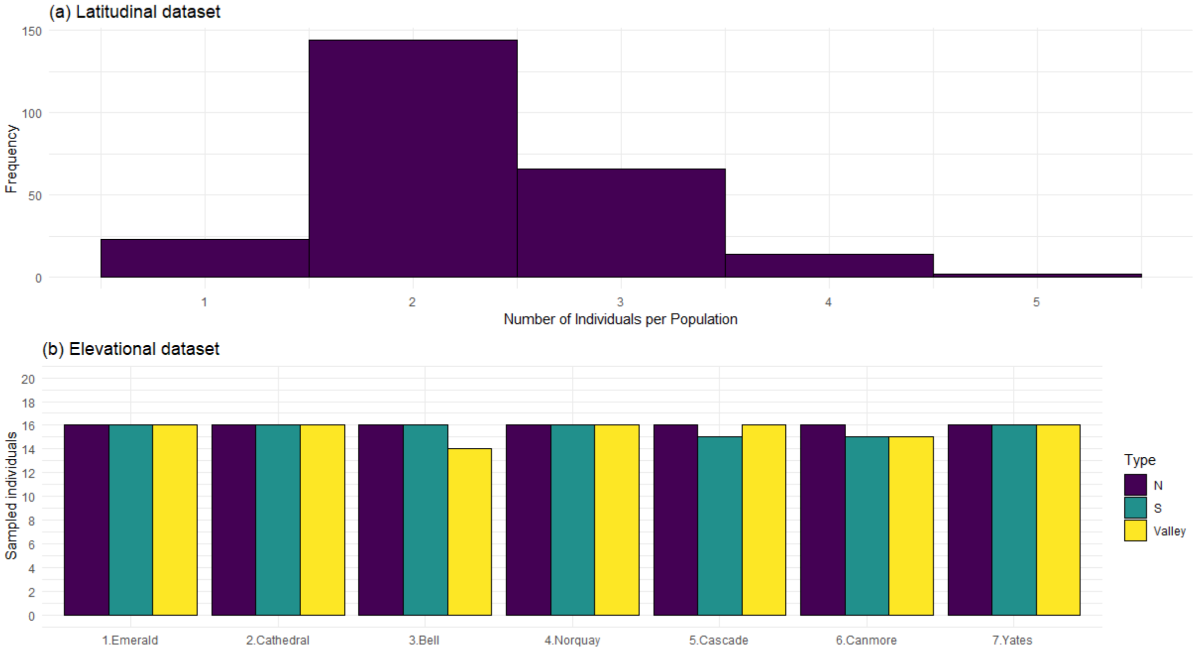

**Figure S1.** (a) Distribution of the number of individuals per population in the 249 populations of the latitudinal dataset. (b) Number of sampled individuals for each population type (north-facing high-altitude = N, south-facing high-altitude = S, Valley = V) in the 21 populations (7 locations) of the elevational dataset.

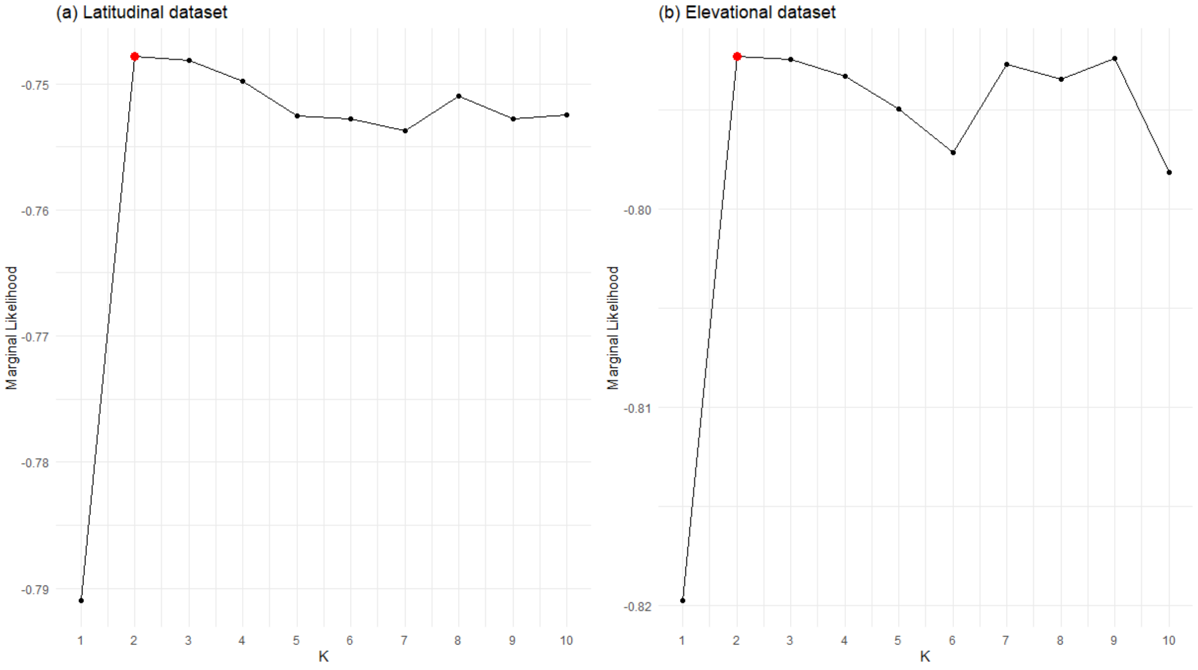

**Figure S2.** Results of the *fastSTRUCTURE* inference. Log-marginal likelihood lower bound (LLBO) of the models for Ks ranging from 1 to 10 for the (a) latitudinal dataset and (b) elevational dataset. In red the maximum LLBO value.

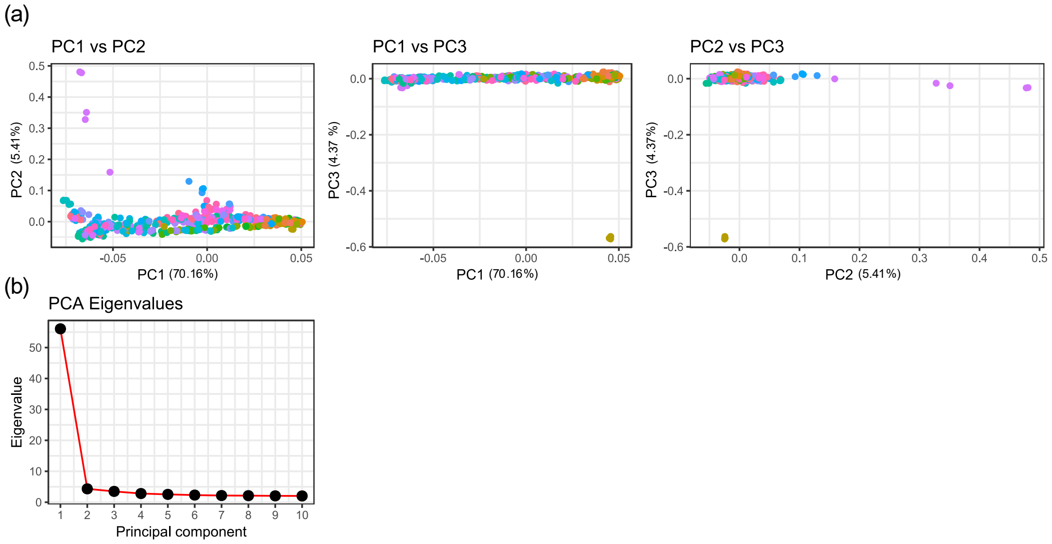

**Figure S3.** PCA of 575 spruce individuals (249 populations) of the latitudinal dataset. Colours represent populations. (a) PC1 against PC2, PC1 against PC3 and PC2 against PC3. (b) Eigenvalues of the first 10 principal components.

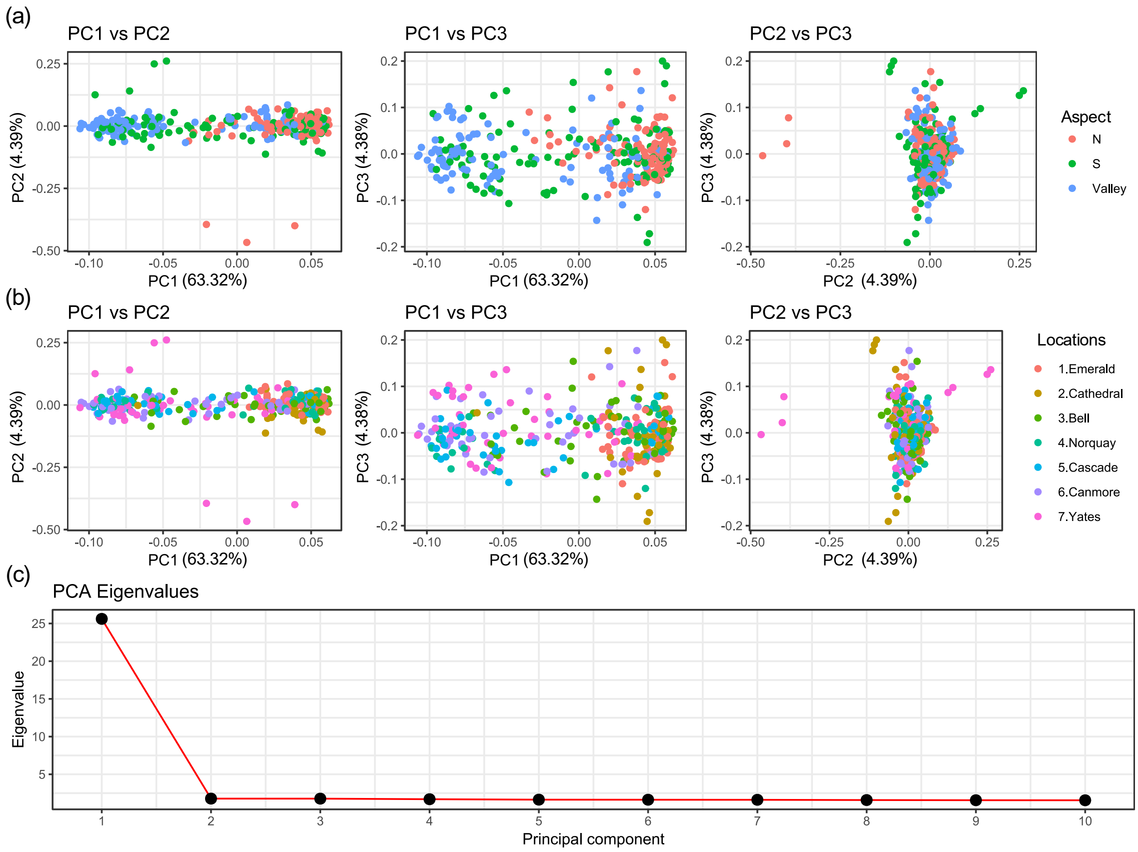

**Figure S4.** PCA of 331 spruce individuals (21 populations, 7 locations) of the elevational dataset. Colours represent populations. (a) PC1 against PC2, PC1 against PC3 and PC2 against PC3 color-coded by environment (north-facing high-altitude, south-facing high-altitude and valley). (b) PC1 against PC2, PC1 against PC3 and PC2 against PC3 color-coded by location. (c) Eigenvalues of the first 10 principal components.

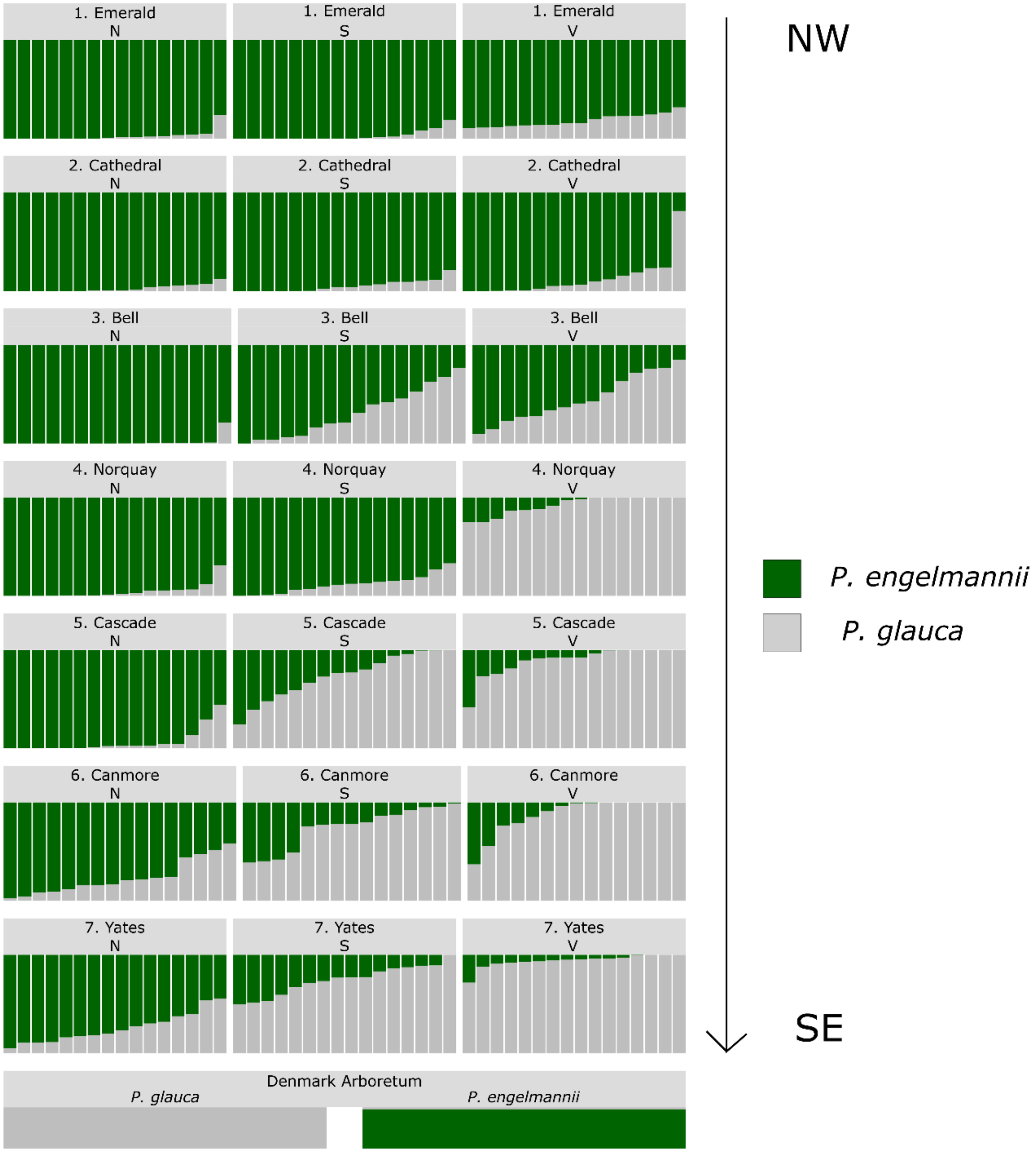

**Figure S5.** Bar plot of *fastSTRUCTURE* ancestry proportions (K = 2) for the 331 individuals of the elevational dataset, including two pure species individuals (Denmark Arboretum) used to identify the two species clusters (*P. glauca* and *P. engelmannii*).

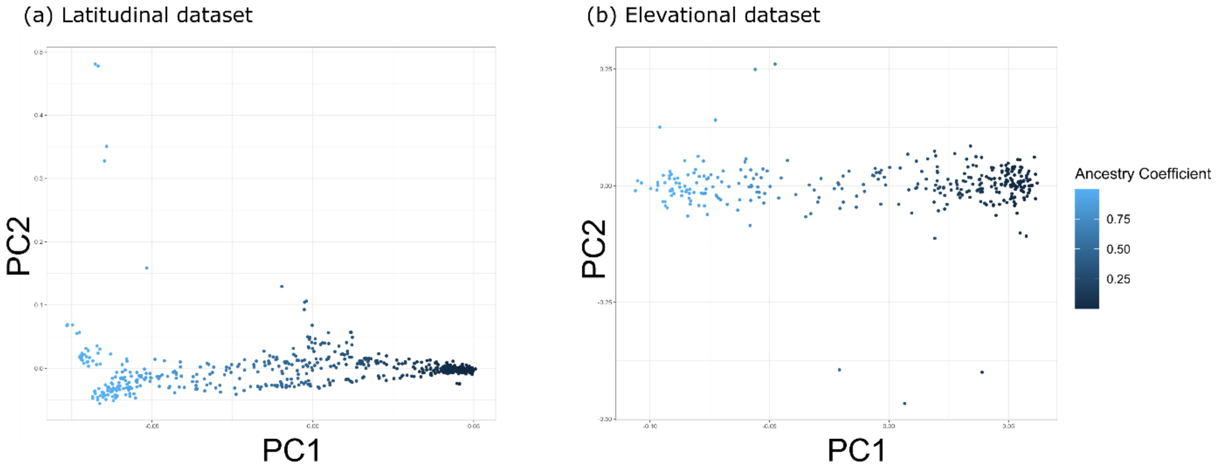

**Figure S6.** PC1 vs PC2 plots color-coded by the ancestry coefficients computed with *fastSTRUCTURE* for the (a) latitudinal dataset and (b) elevational dataset. The correlation between PC1 and ancestry coefficients was 0.999 in both datasets.

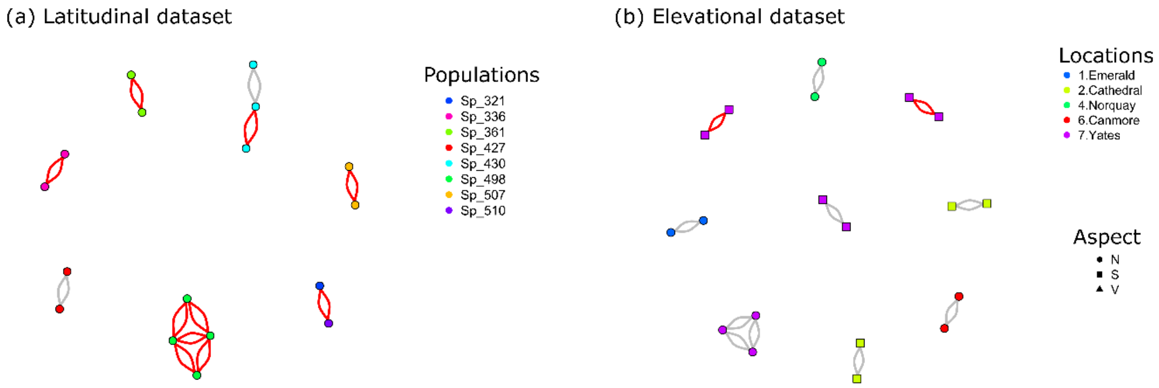

**Figure S7.** Network of individuals genomic-based relatedness showing third-degree relationships and above (kin > 0.125) for the (a) latitudinal and (b) elevational dataset. In grey third-degree relationships (0.125 < kin < 0.25), in red second-degree relationships (kin ~0.25).

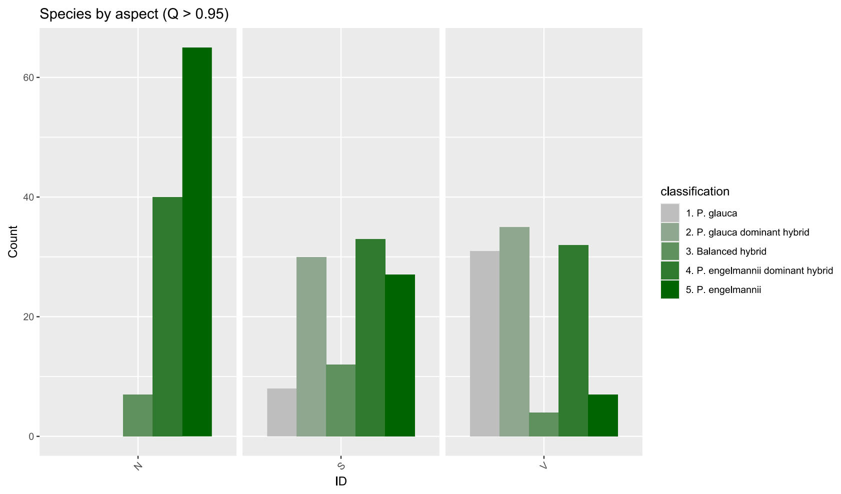

**Figure S8.** Summary of the distribution of individual ancestry categories across environments in the elevational datasets (N: north-facing slopes, S: south-facing slopes, V: valleys). Individuals were classified according to the admixture coefficient computed with fastSTRUCTURE, using the following criteria: *P. glauca* ($q \geq 0.95$), *P. glauca* dominant hybrid ($0.60 \leq q < 0.95$), balanced hybrid ($0.40 < q < 0.60$), *P. engelmannii* dominant hybrid ($0.05 \leq q \leq 0.40$), *P. engelmannii* ($q < 0.05$).

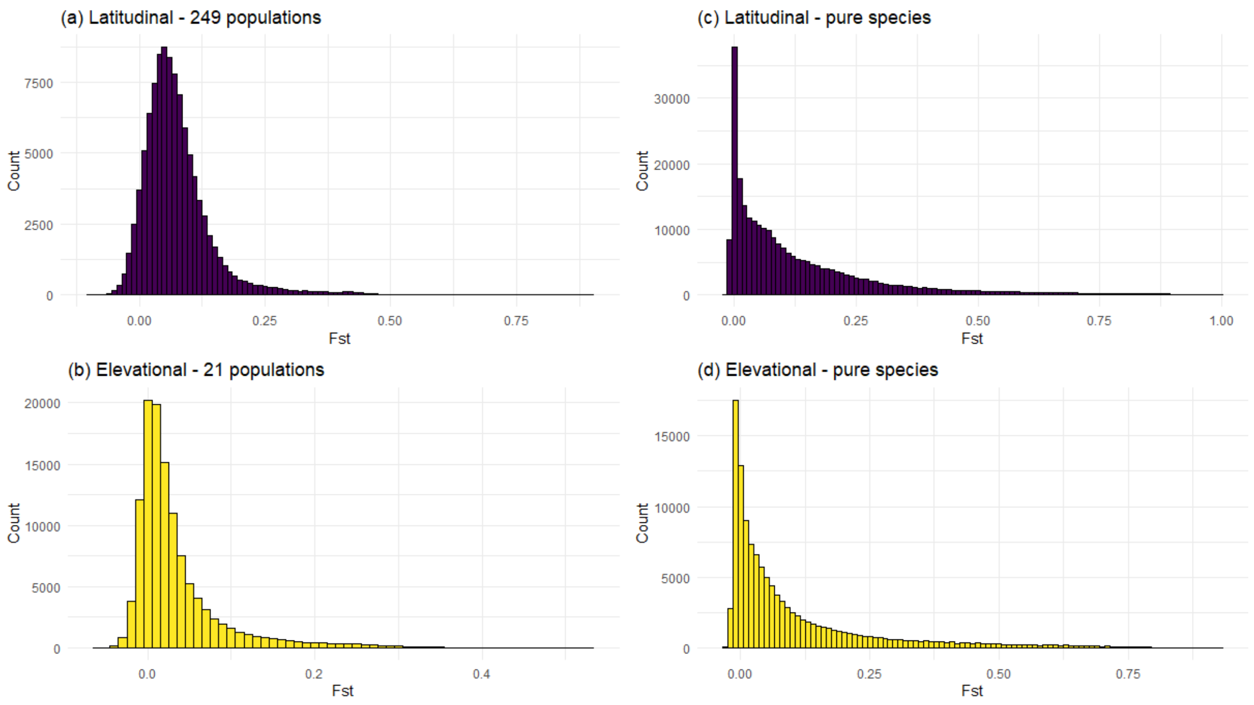

**Figure S9.** Distribution of $F_{ST}$ between: (a) the 249 populations of the latitudinal dataset. (b) pure *P. glauca* and pure *P. engelmannii* in the latitudinal dataset. (c) the 21 populations of the elevational dataset. (d) pure *P. glauca* and pure *P. engelmannii* in the elevational dataset.

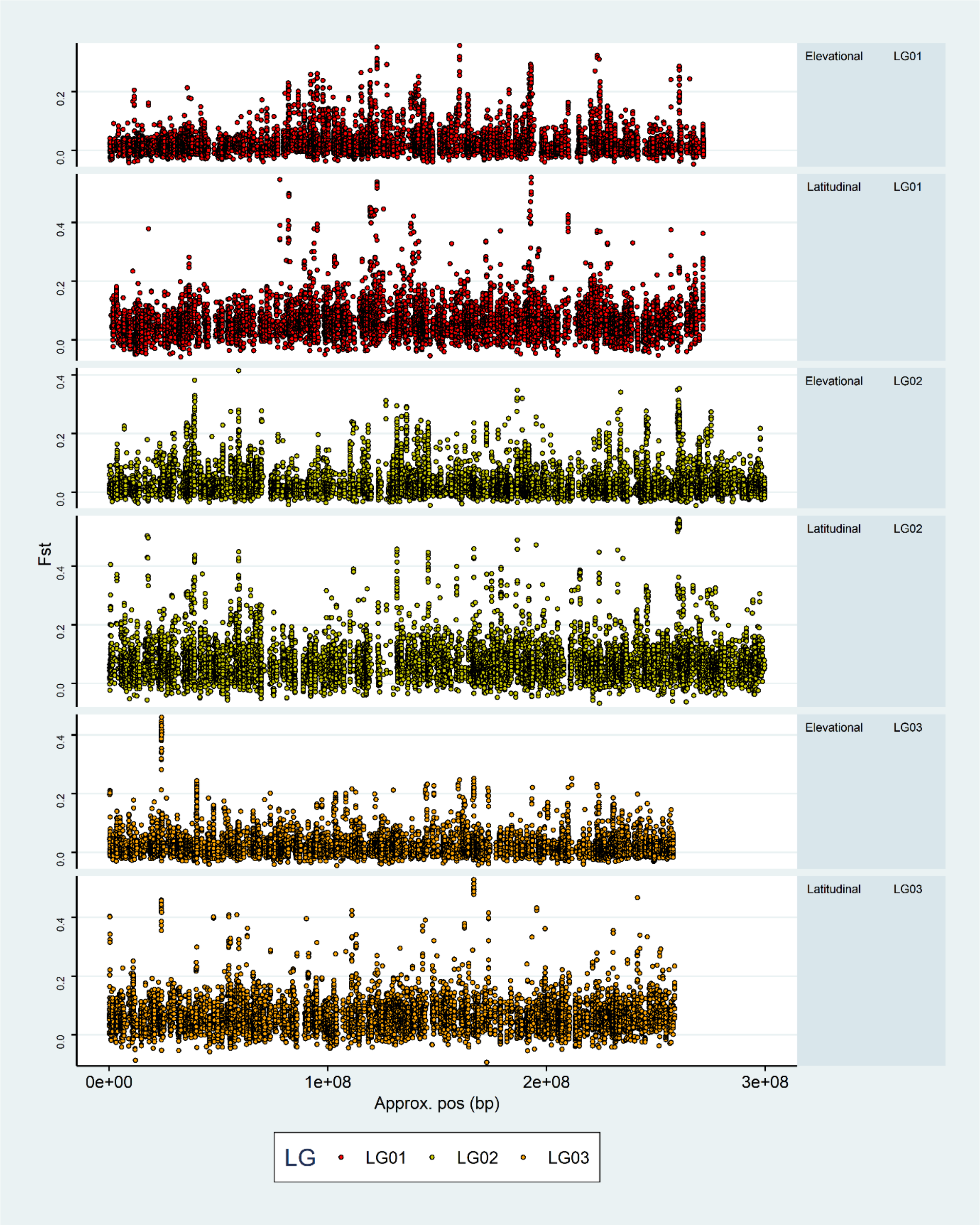
Elevational LG01
Latitudinal LG01
Elevational LG02
Latitudinal LG02
Elevational LG03
Latitudinal LG03
Fst
0.0
0.2
0.4
0e+00
1e+08
2e+08
3e+08
Approx. pos (bp)
LG
LG01
LG02
LG03

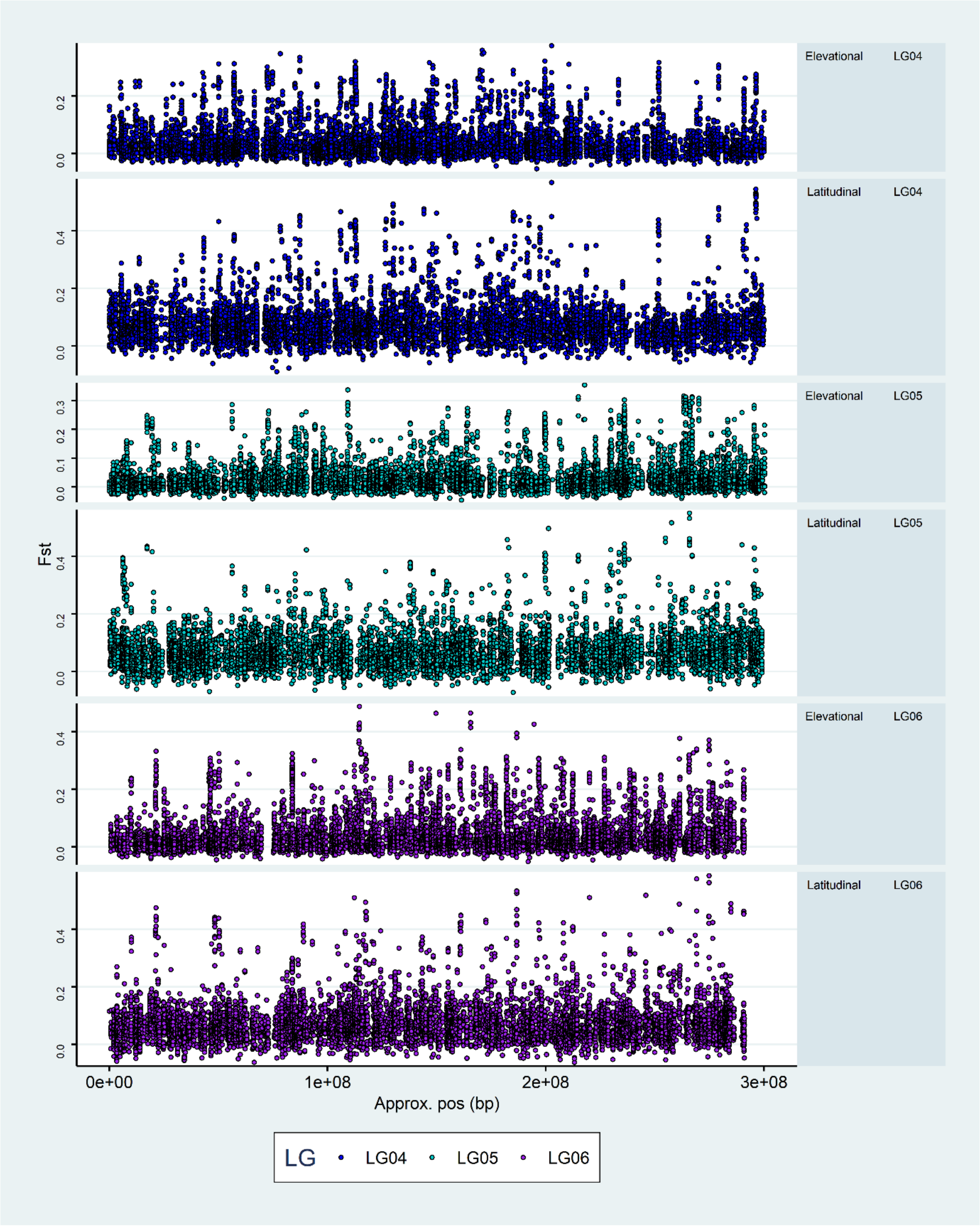

Elevational LG04
Latitudinal LG04
Elevational LG05
Latitudinal LG05
Elevational LG06
Latitudinal LG06
Fst
0.0
0.1
0.2
0.3
0.4
0e+00
1e+08
2e+08
3e+08
Approx. pos (bp)
LG
LG04
LG05
LG06

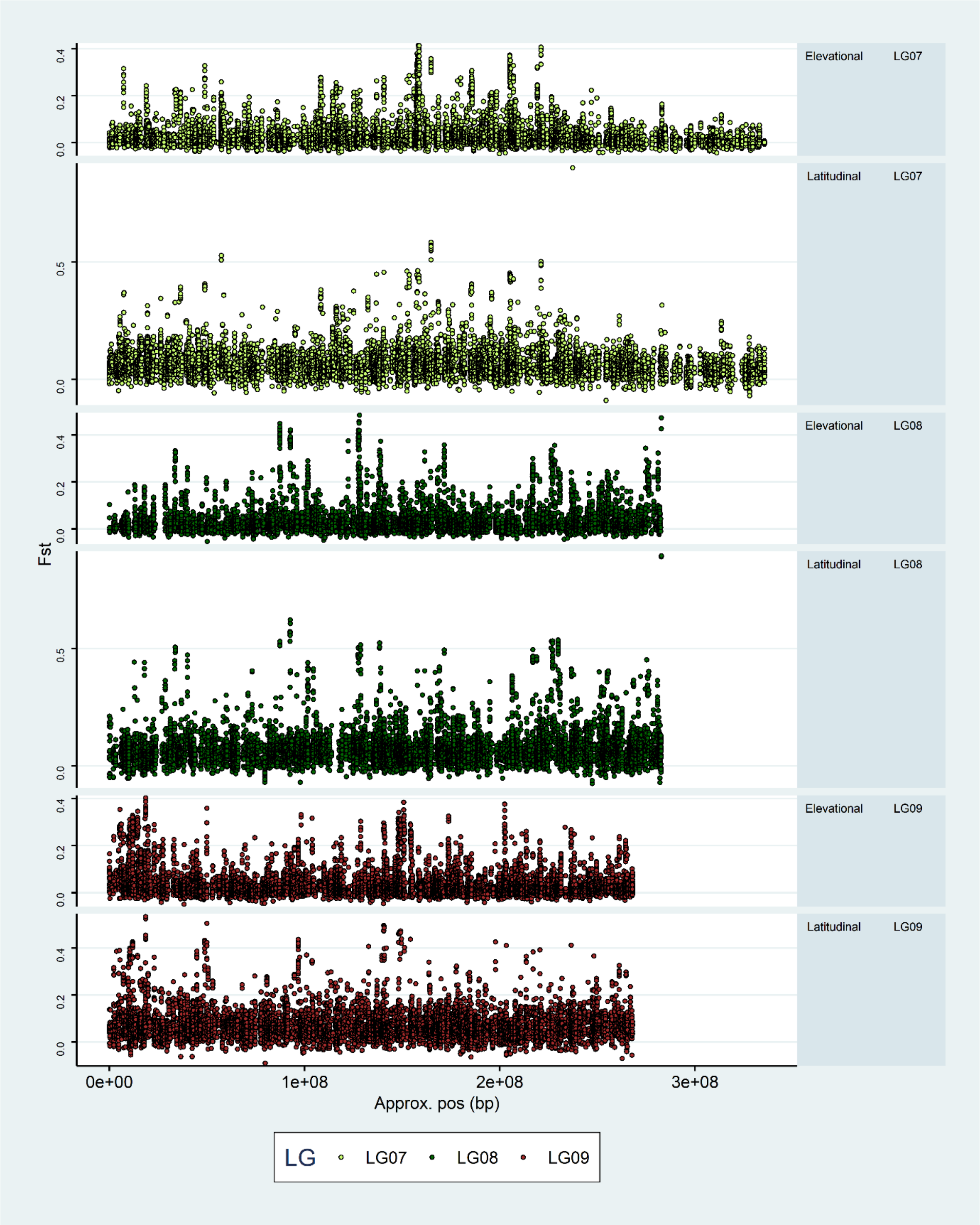

Elevational
LG07
Latitudinal
LG07
Elevational
LG08
Latitudinal
LG08
Elevational
LG09
Latitudinal
LG09
Fst
0.0
0.2
0.4
0.5
0e+00
1e+08
2e+08
3e+08
Approx. pos (bp)
LG
LG07
LG08
LG09

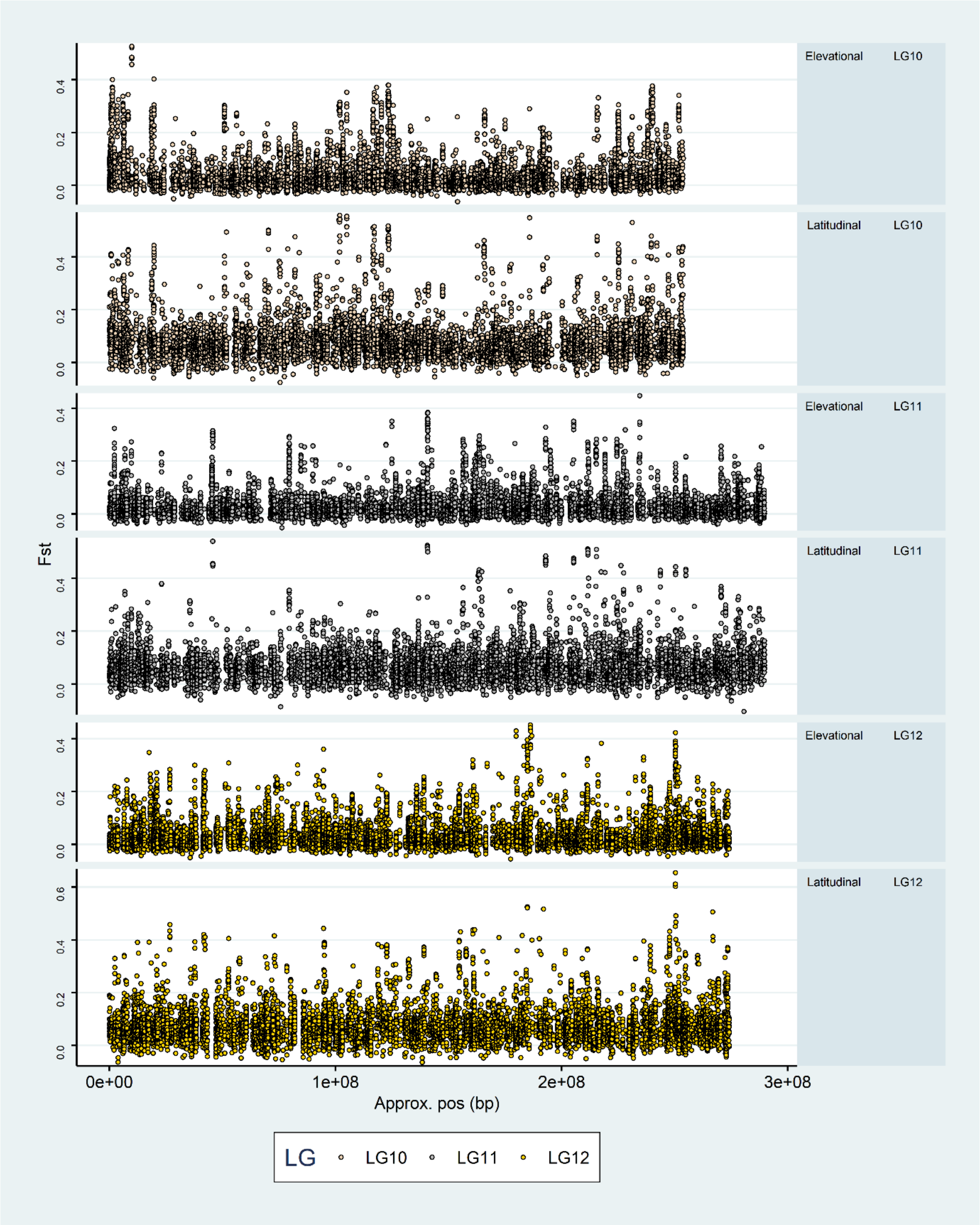

**Figure S10.** $F_{ST}$ Manhattan plots for the elevational and latitudinal datasets overlaying the same position, based on the super-scaffold assembly.

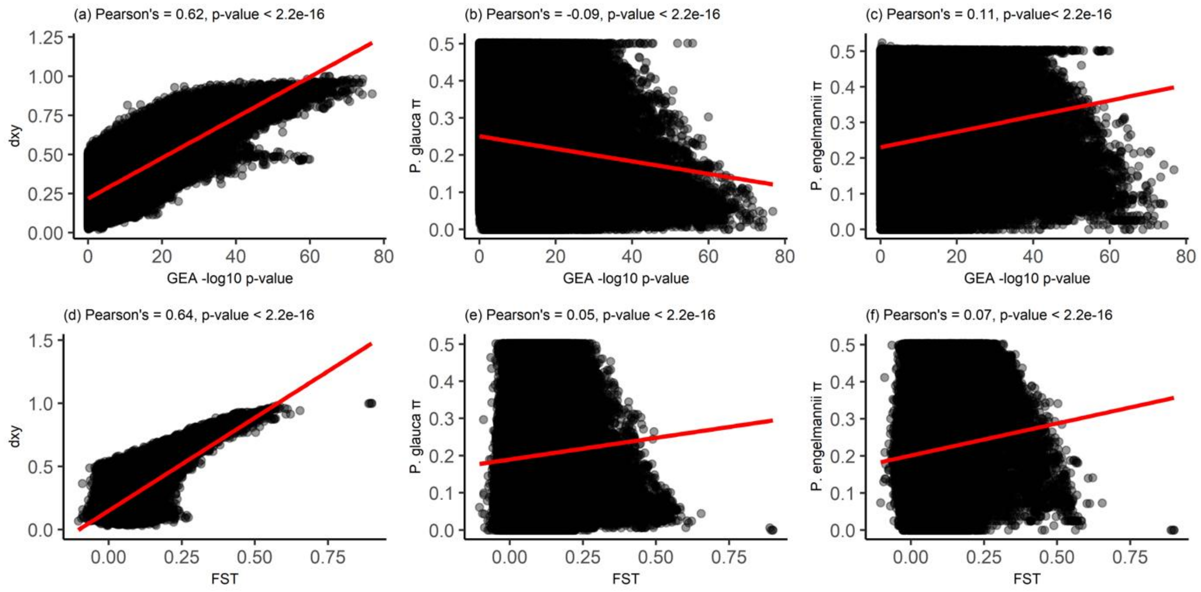

**Figure S11.** Correlation between GEA p-values and $F_{ST}$ values with nucleotide diversity (π) within pure species populations (Q ≥ 0.95) and among them (*dxy*) in the latitudinal dataset. (a) *dxy* vs GEA p-values. (b) *P. glauca* π vs GEA p-values. (c) *P. engelmannii* π vs GEA p-values. (d) $F_{ST}$ vs *dxy*. (e) $F_{ST}$ vs *P. glauca* π. (f) $F_{ST}$ vs *P. engelmannii* π.

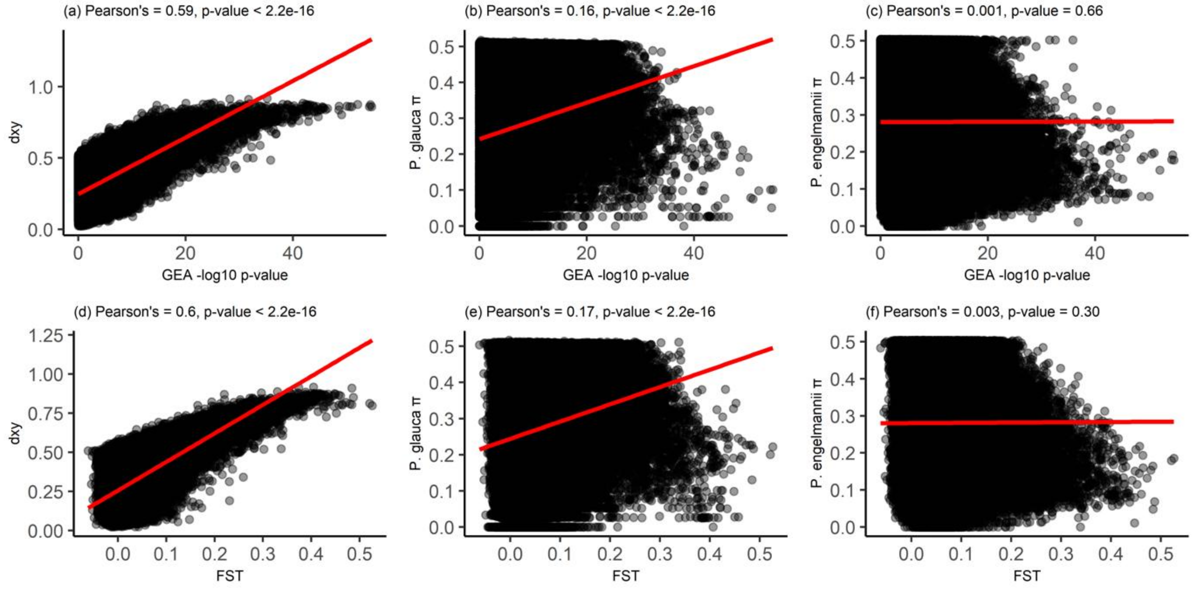

**Figure S12.** Correlation between GEA p-values and FST values with nucleotide diversity (π) within pure species populations (Q ≥ 0.95) and among them (*dxy*) in the elevational dataset. (a) dxy vs GEA p-values. (b) *P. glauca* π vs GEA p-values. (c) *P. engelmannii* π vs GEA p-values. (d) FST vs dxy. (e) FST vs *P. glauca* π. (f) FST vs *P. engelmannii* π.

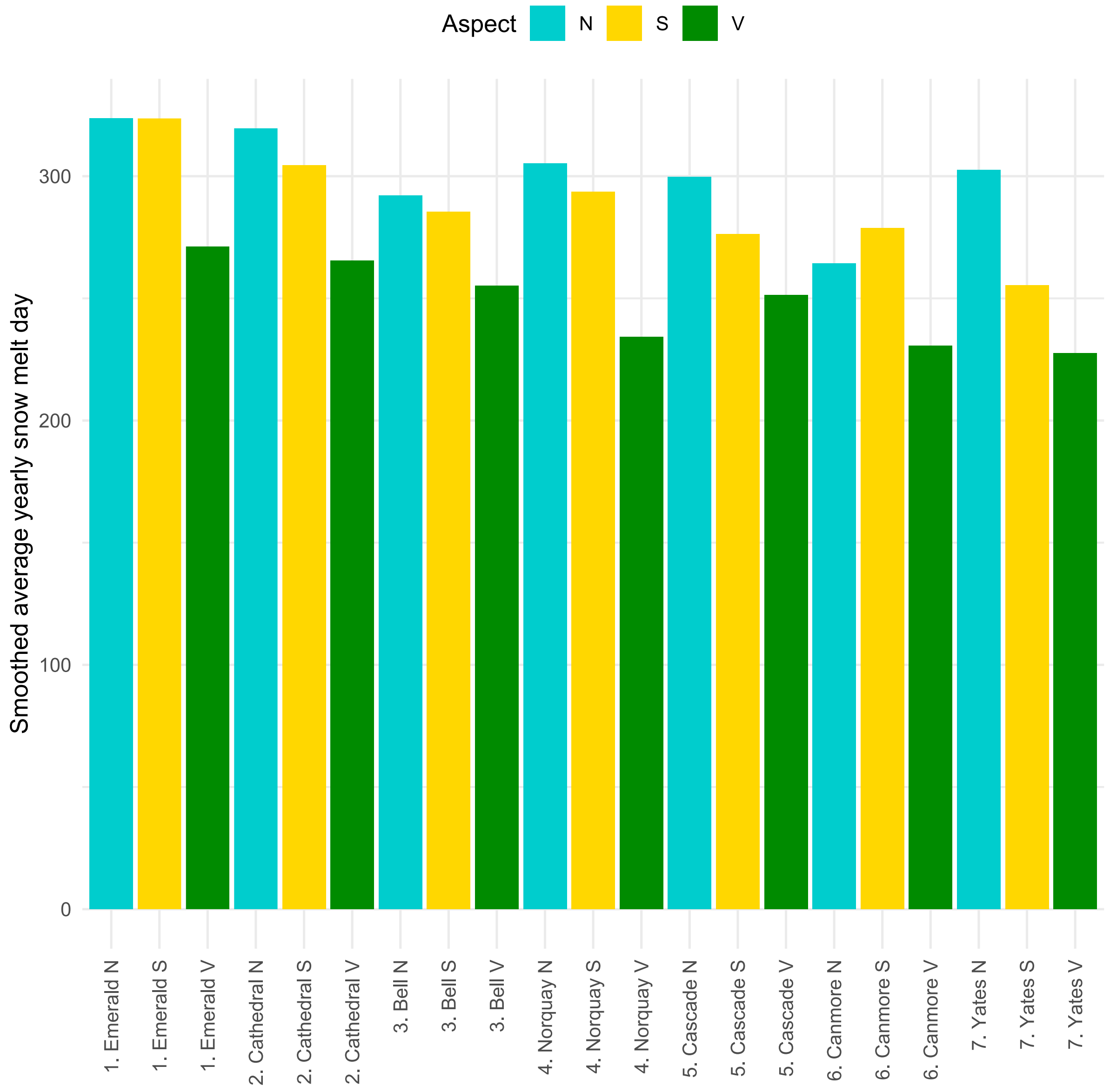

**Figure S13.** Distribution of snowmelt date across the locations and environments of the elevational dataset.

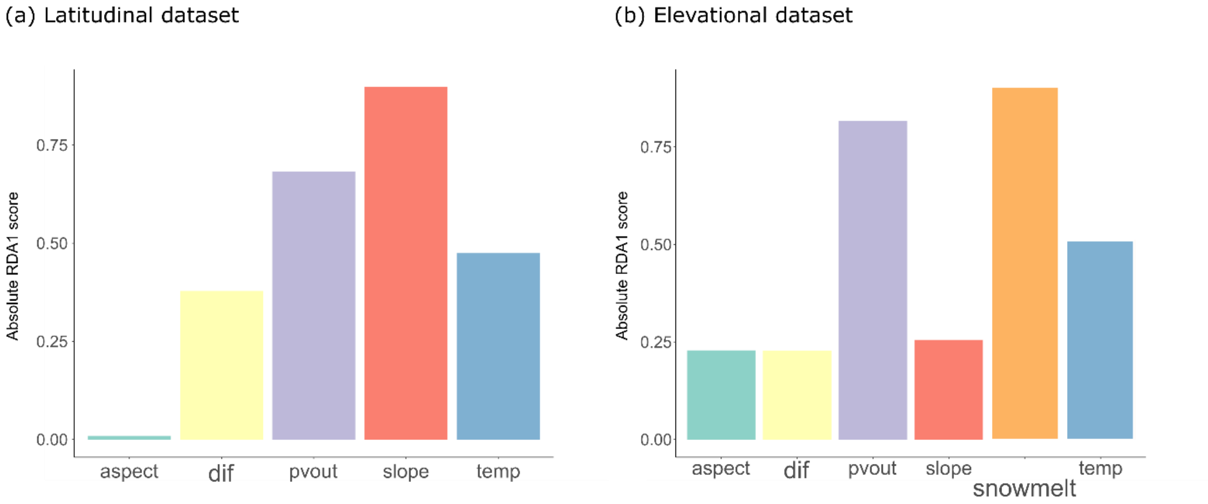

**Figure S14.** RDA-based contribution of the selected environmental variables in explaining h-index variation in the (a) latitudinal and (b) elevational dataset.

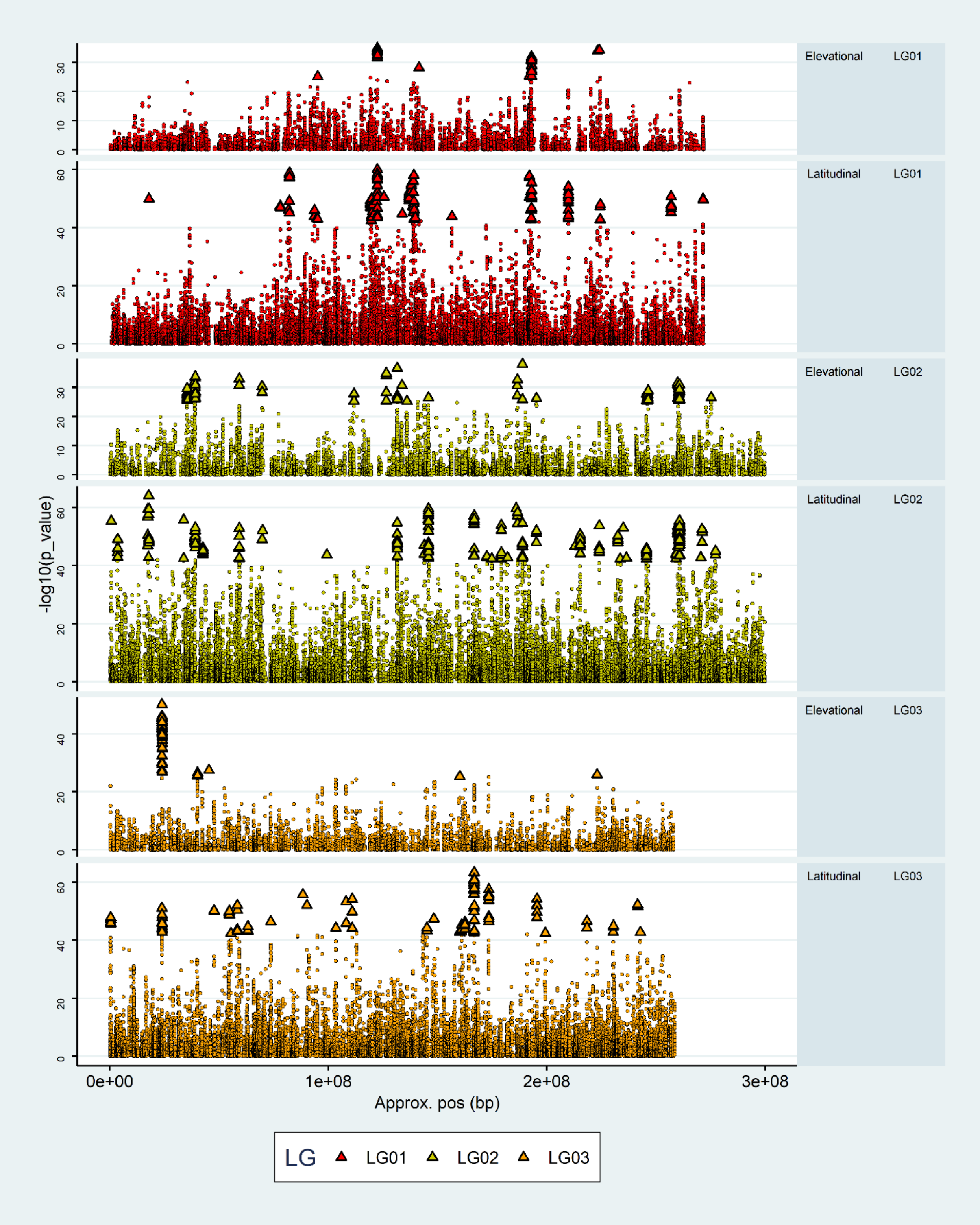
Elevational LG01
Latitudinal LG01
Elevational LG02
Latitudinal LG02
Elevational LG03
Latitudinal LG03
-log10(p_value)
Approx. pos (bp)
0e+00
1e+08
2e+08
3e+08
LG
LG01
LG02
LG03

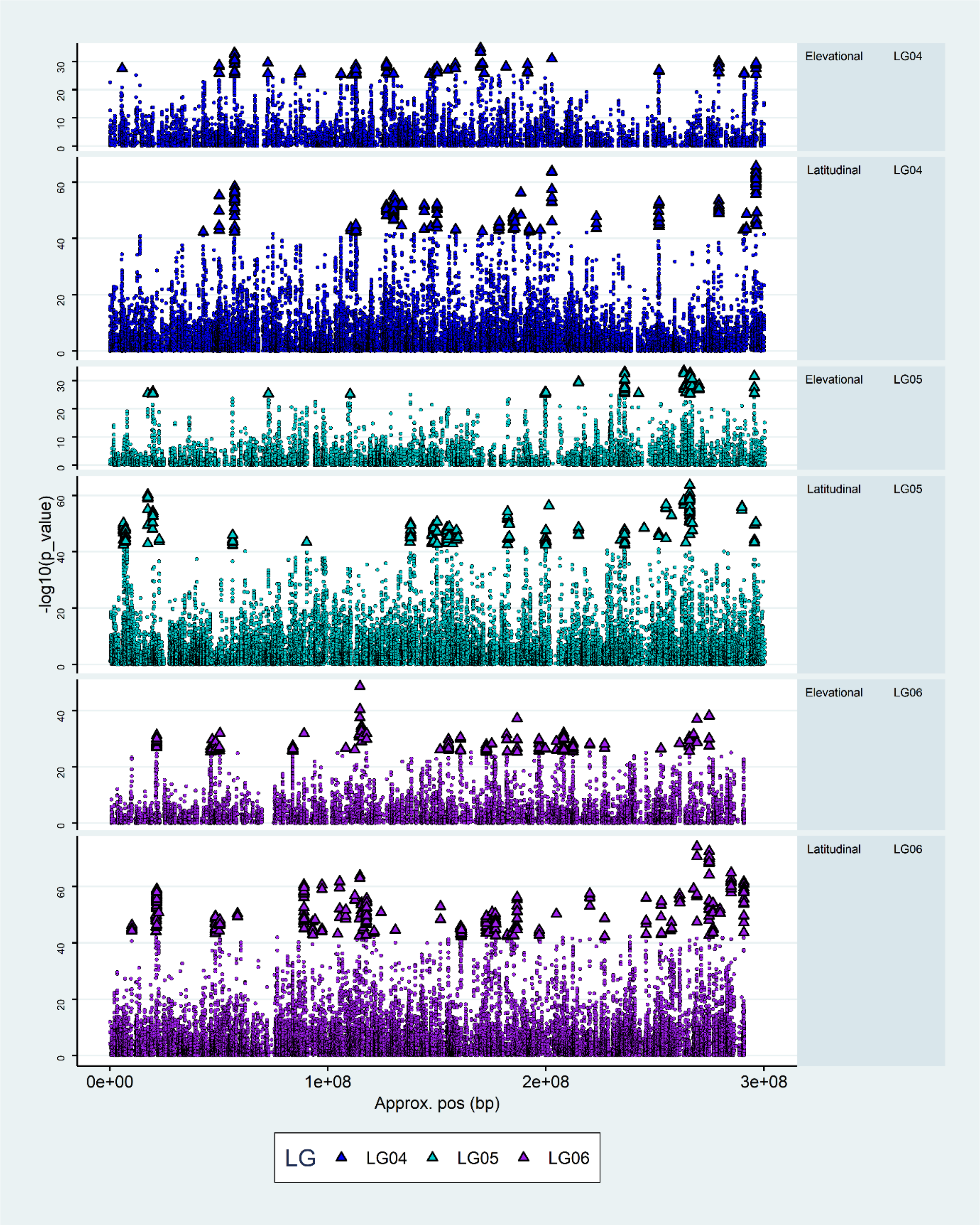
Elevational LG04
Latitudinal LG04
Elevational LG05
Latitudinal LG05
Elevational LG06
Latitudinal LG06
-log10(p_value)
0e+00
1e+08
2e+08
3e+08
Approx. pos (bp)
LG
LG04
LG05
LG06

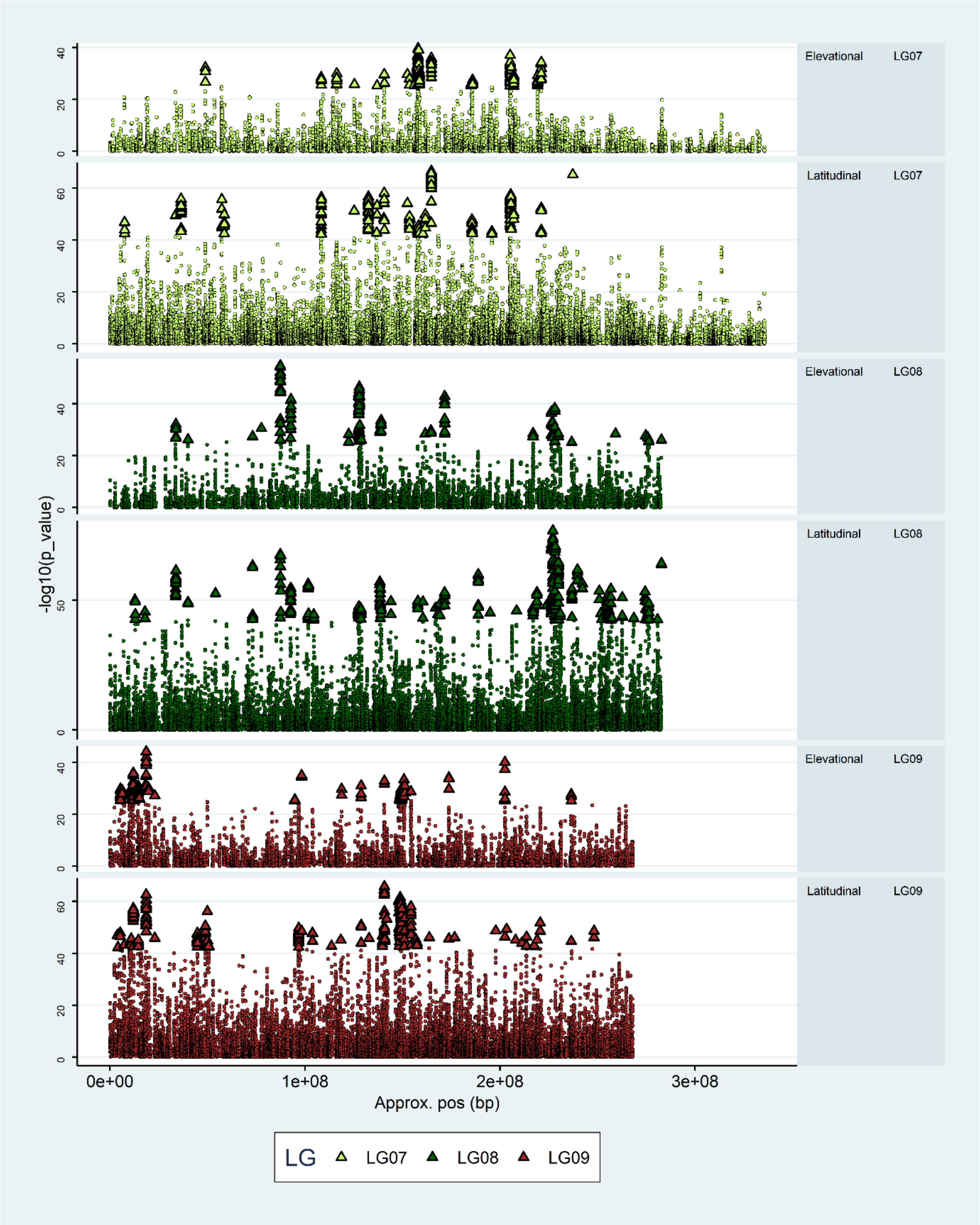

-log10(p_value)
Elevational LG07
Latitudinal LG07
Elevational LG08
Latitudinal LG08
Elevational LG09
Latitudinal LG09
0e+00
1e+08
2e+08
3e+08
Approx. pos (bp)
LG LG07 LG08 LG09

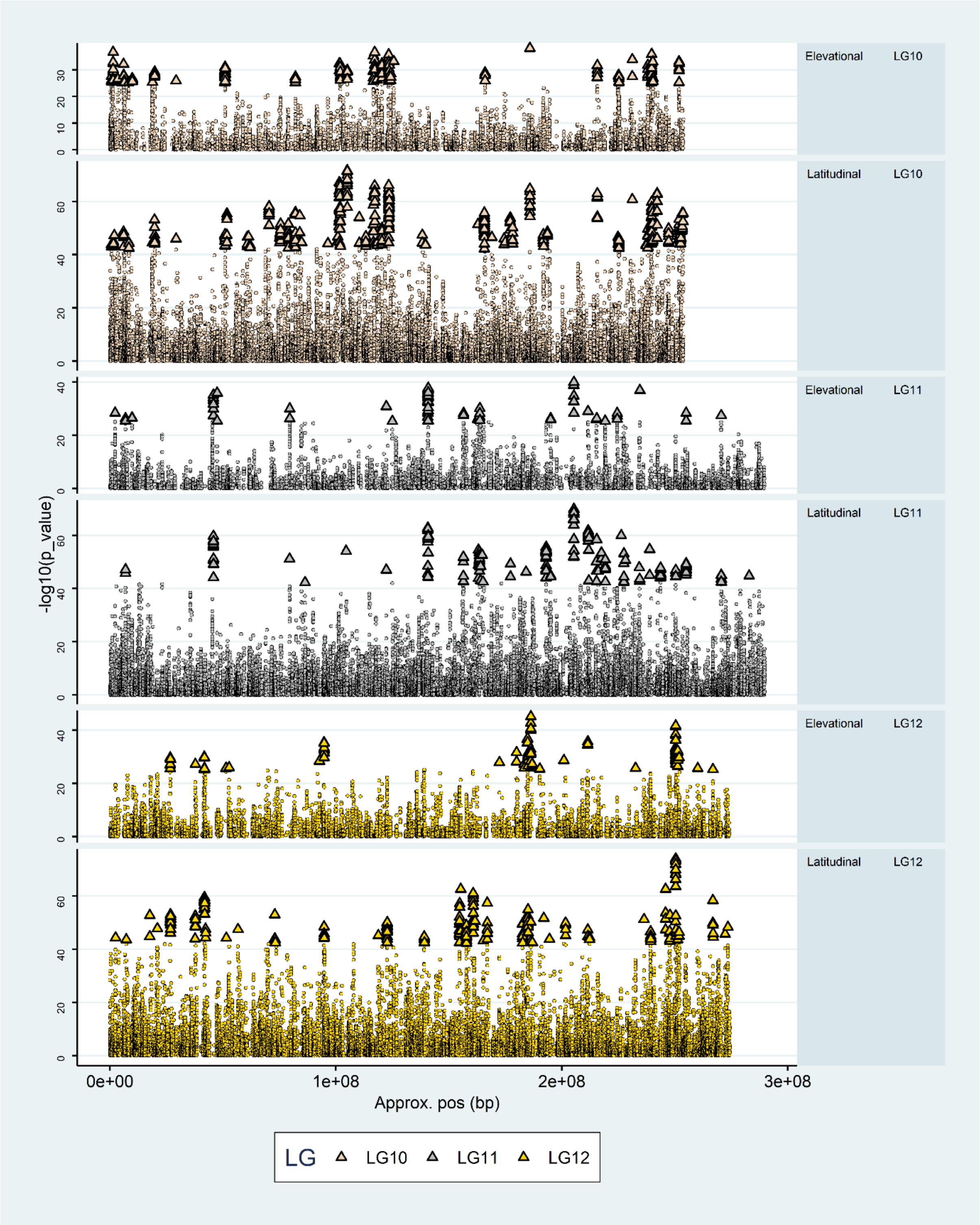

**Figure S15.** GEA p-values Manhattan plots for the elevational and latitudinal datasets overlaying the same position, based on the super-scaffold assembly. Triangles represent the 1% lowest GEA p-values SNPs in each dataset.

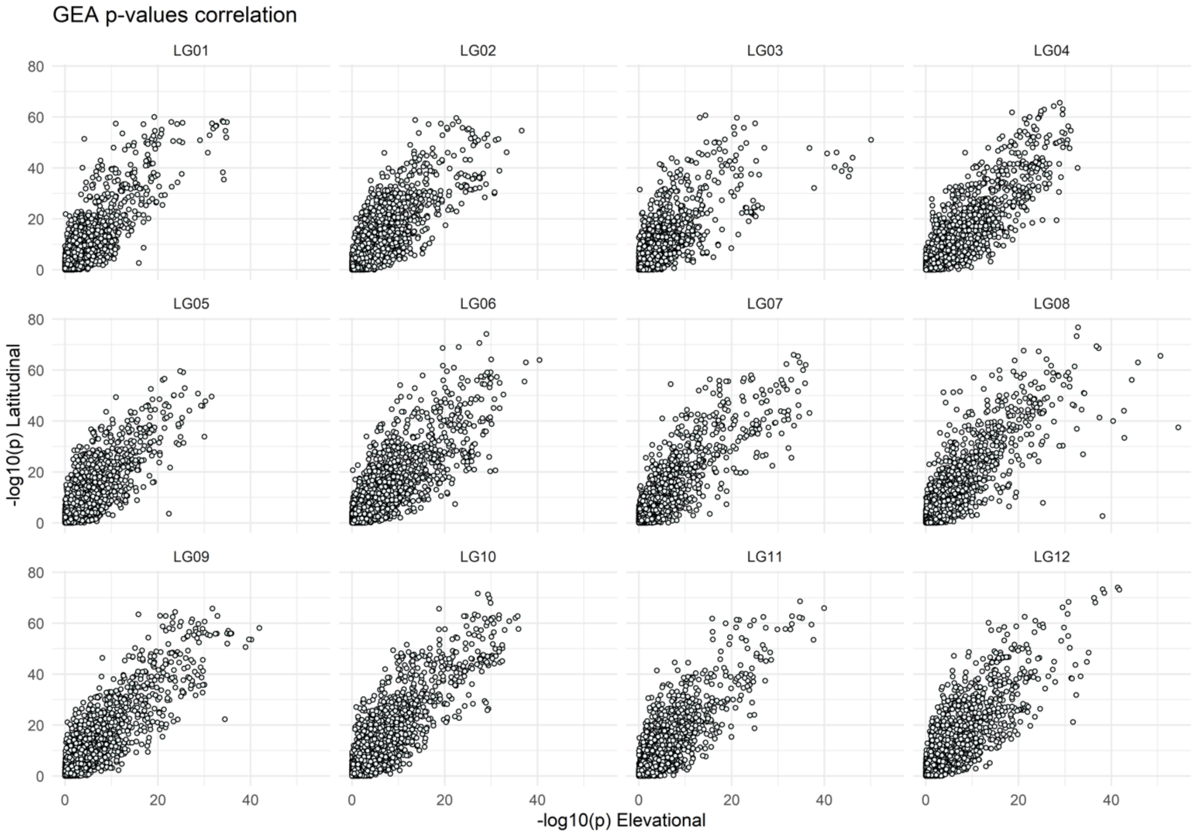

**Figure S16.** Correlation of GEA p-values of the 31,667 shared SNPs between the elevational and latitudinal dataset by linkage group.

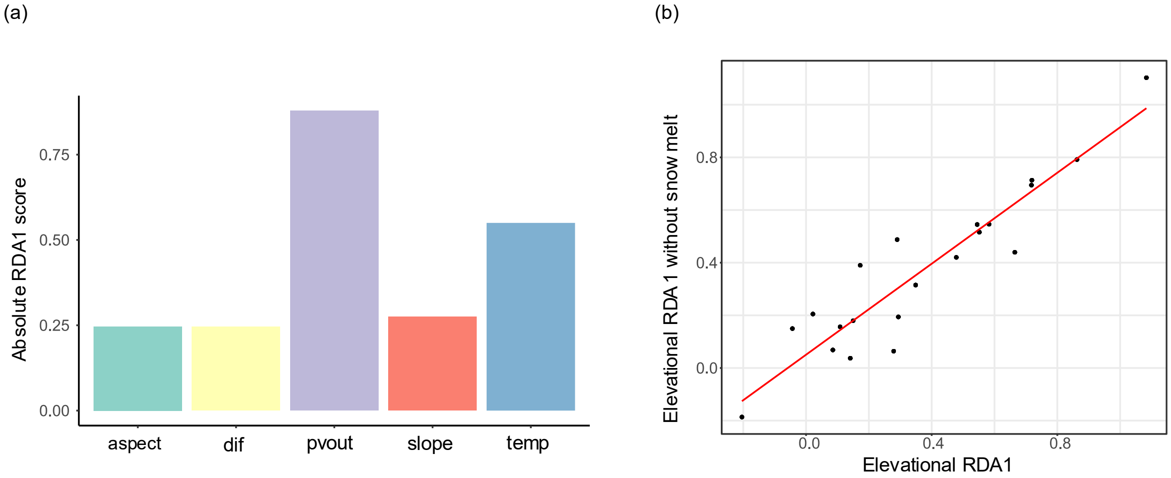

**Figure S17.** RDA without snow melt day. (a) RDA-based contribution of the selected environmental variables in explaining h-index variation in the elevational dataset after excluding snow melt day. (b) Correlation between the elevational dataset RDA1 computed with and without the snow melt day variable (Pearson's: 0.93; Kendall's tau: 0.76).

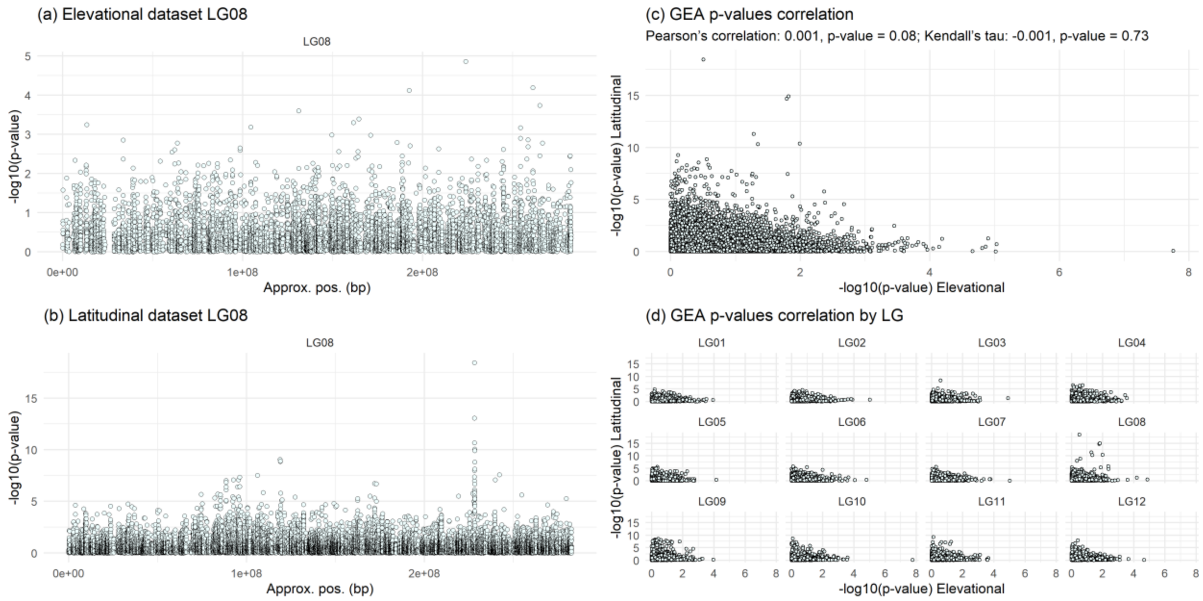

**Figure S18.** LG08 Manhattan plots of p-values, derived from log-likelihood ratio test-based GEAs incorporating population structure correction for the (a) elevational and (b) latitudinal datasets. (c) Correlation between the elevational and latitudinal datasets p-values, derived from log-likelihood ratio test-based GEAs incorporating population structure correction. (d) Correlation between the elevational and latitudinal datasets p-values by linkage group, derived from log-likelihood ratio test-based GEAs incorporating population structure correction.

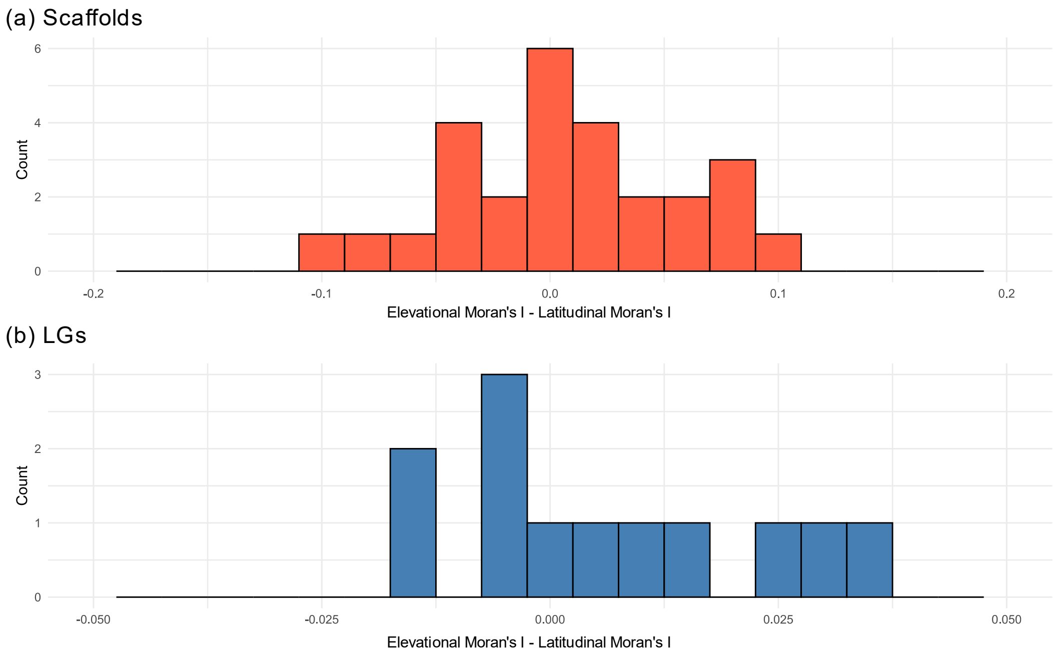

**Figure S19.** Distribution of the difference in $F_{ST}$ Moran's I by (a) scaffolds and (b) LGs between the elevational and latitudinal datasets. If there were no difference in spatial autocorrelation, the histogram would be symmetric around zero. However, if autocorrelation were stronger in the elevational dataset, the distribution would be enriched for positive values.

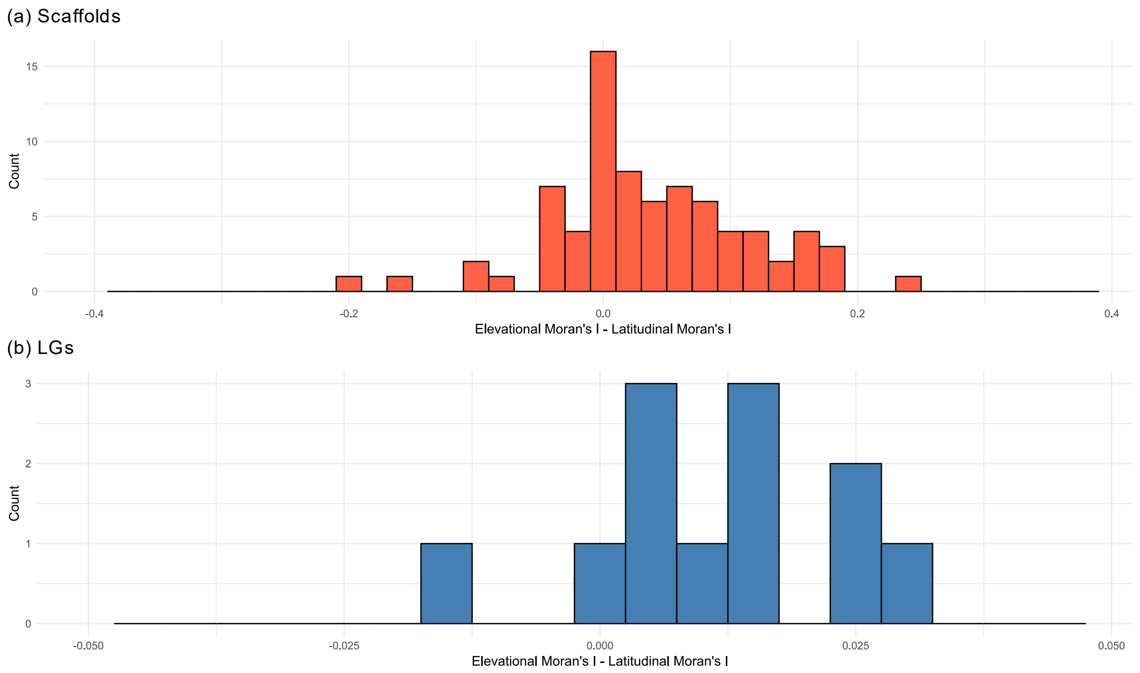

**Figure S20.** Distribution of the difference in GEA p-values Moran's I by (a) scaffolds and (b) LGs between the elevational and latitudinal datasets. If there were no difference in spatial autocorrelation, the histogram would be symmetric around zero. However, if autocorrelation were stronger in the elevational dataset, the distribution would be enriched for positive values.

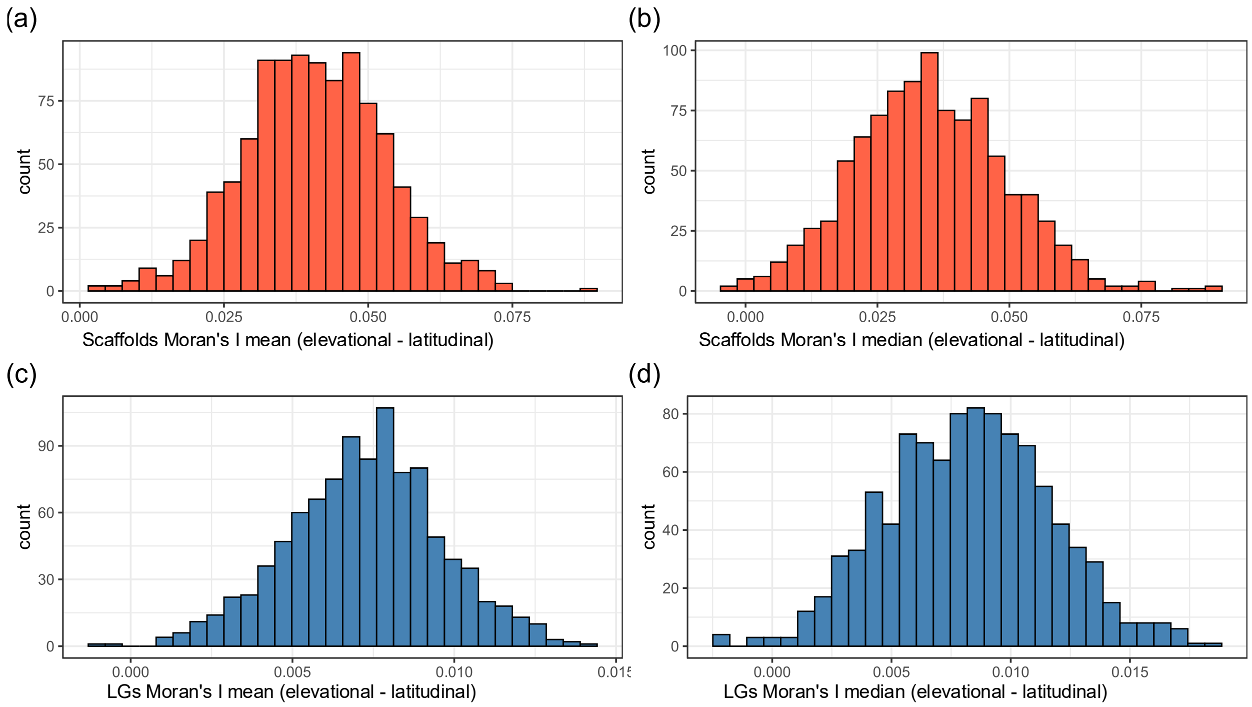

**Figure S21.** Distribution of the mean and median of the difference between GEA p-values Moran's I of each of 1,000 subsampled elevational datasets (subsampled to two individuals per population) and GEA p-values Moran's I of the latitudinal dataset, assessed by (a and b) scaffolds and (c and d) LGs. The mean and median differences remain consistently positive even under subsampling, confirming that the stronger spatial autocorrelation in GEA p-values observed in the fully sampled elevational dataset is not driven by differences in sampling strategies between the elevational and latitudinal datasets.

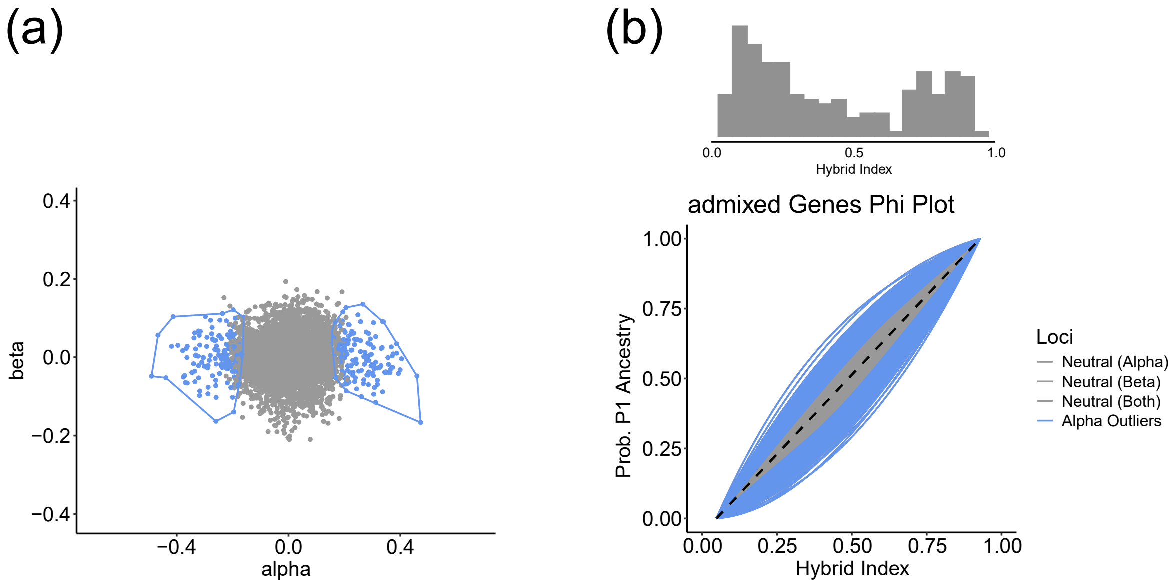

**Figure S22.** Elevational dataset *BGC* results. (a) *BGC* α vs. β plot. Each point is a SNP, and the polygons group the outlier loci. Alpha outliers are blue and neutral loci are grey. (b) *BGC* ancestry probability plot.

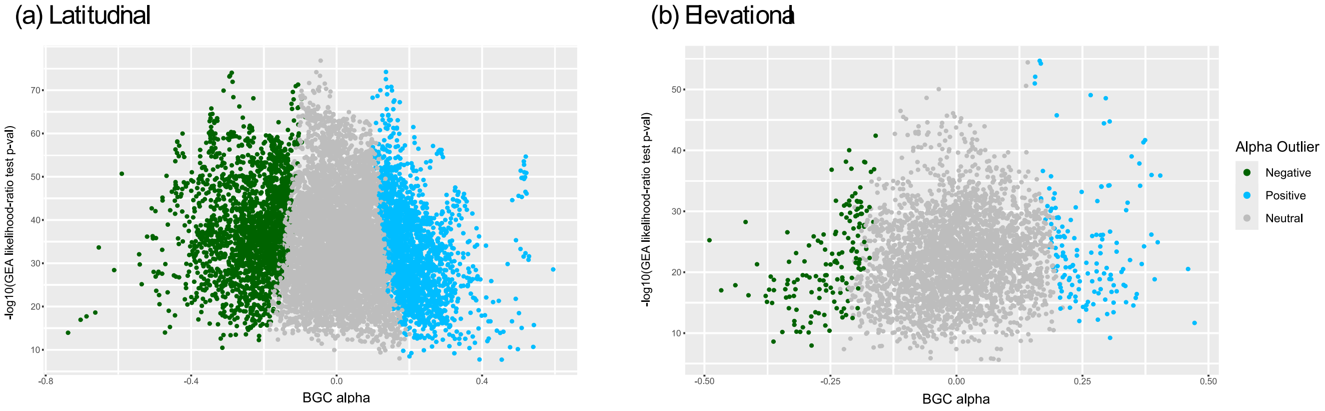

**Figure S23.** Comparison of *BGC* α-values with the GEA p-values for the (a) Latitudinal and (b) Elevational datasets.

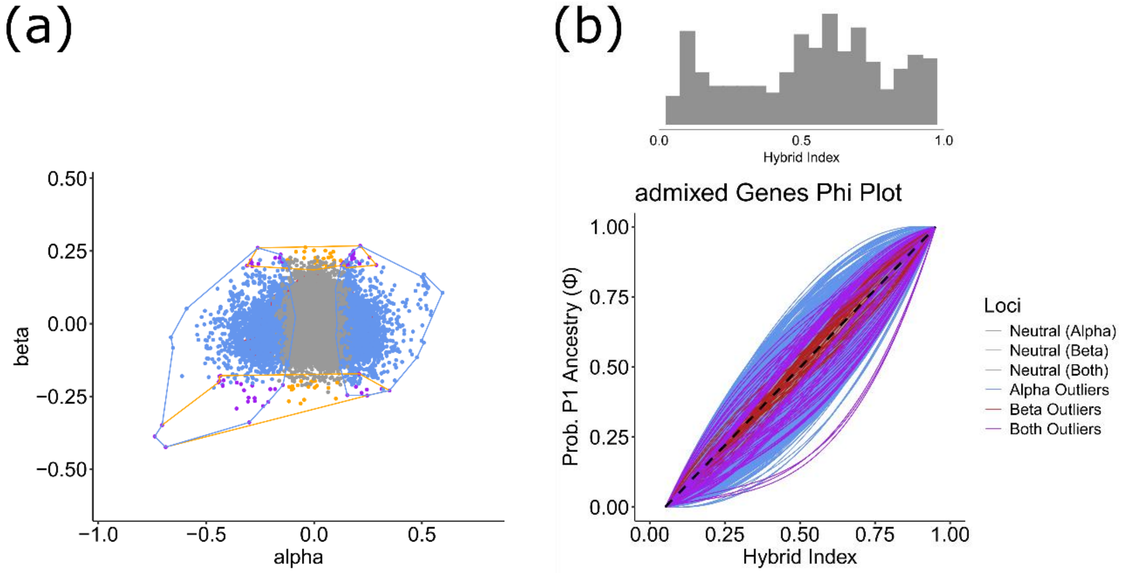

**Figure S24.** Latitudinal dataset *BGC* results. (a) *BGC* α vs. β plot. Each point is a SNP, and the polygons group the outlier loci. Alpha outliers are blue, beta outliers are orange, neutral loci are grey and outliers for both alpha and beta are purple. (b) *BGC* ancestry probability plot.

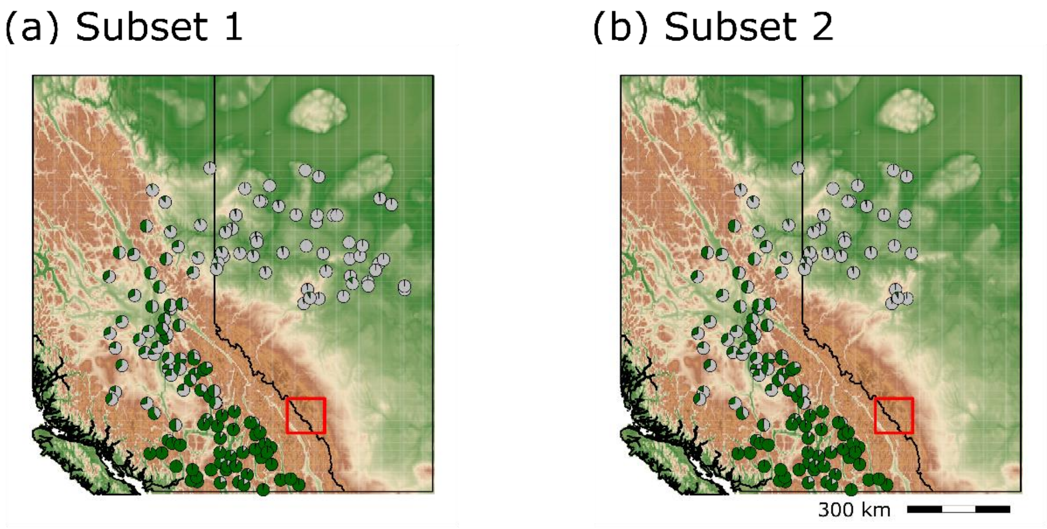

**Figure S25.** Sampling and ancestry distribution of the two subsets from the latitudinal dataset used for the follow-up *BGC* analyses: (a) Subset 1 and (b) Subset 2. Each pie shows the ancestry composition of populations, with its size proportional to the number of individuals sampled in each population. Dark green represents *P. engelmannii* ancestry and gray represents *P. glauca* ancestry.

LG01

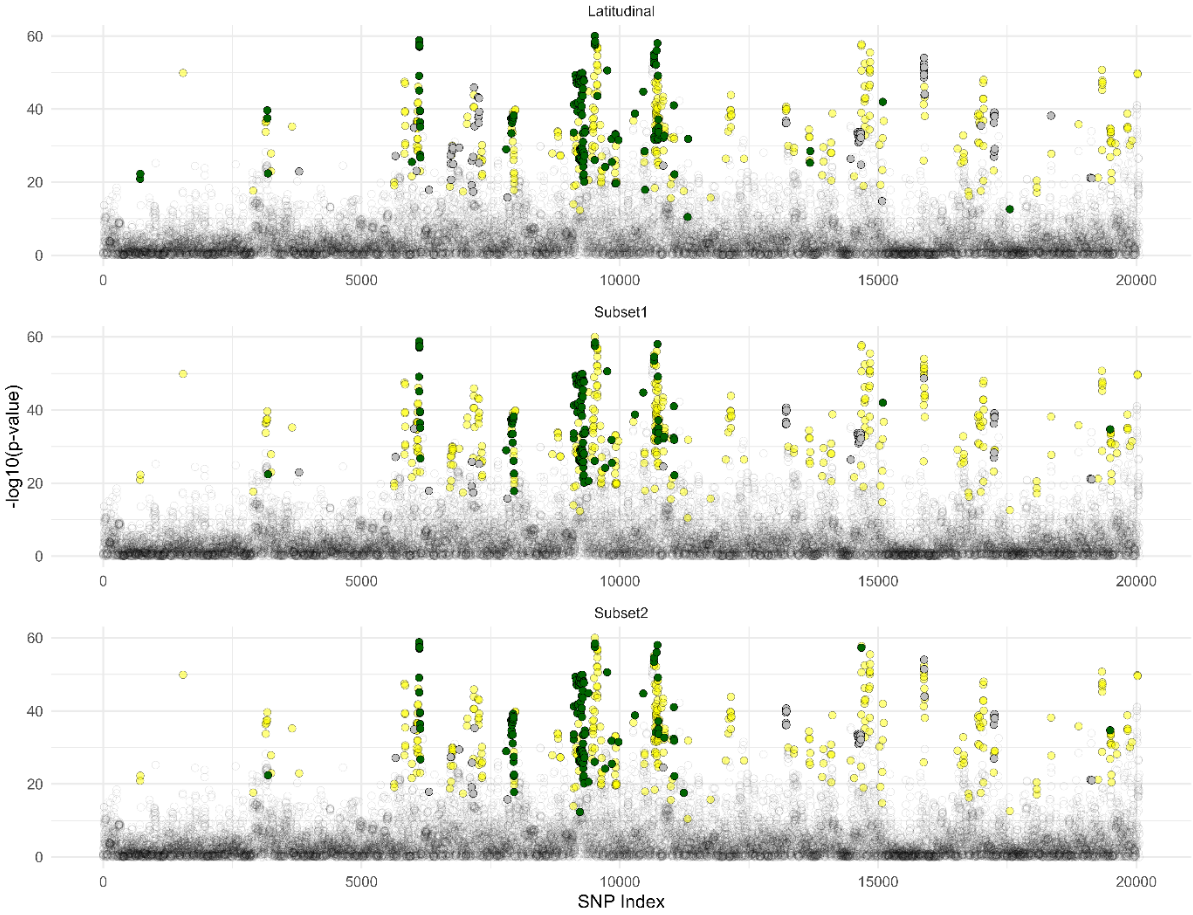

Latitudinal
Subset1
Subset2
-log10(p-value)
SNP Index
0
20
40
60
0
5000
10000
15000
20000

LG02

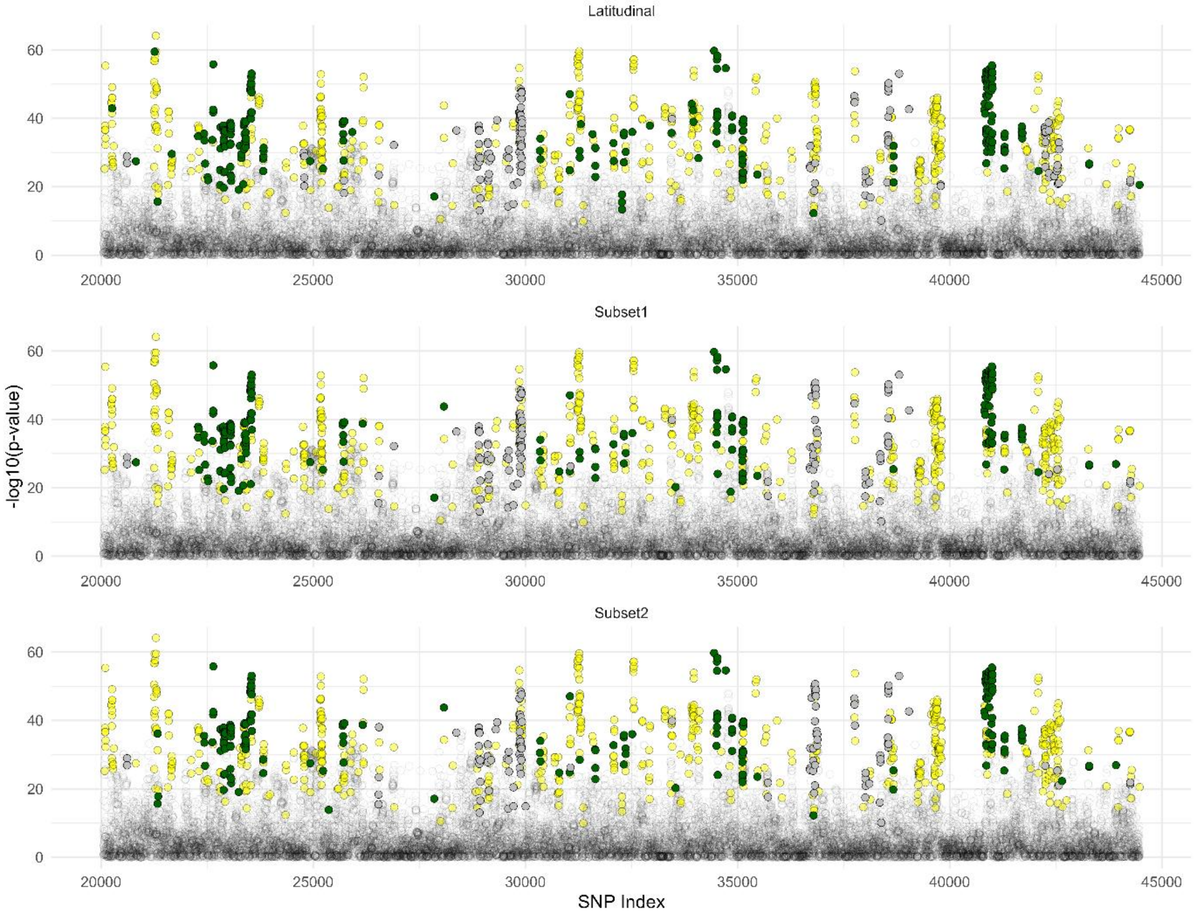

Latitudinal
Subset1
Subset2
-log10(p-value)
SNP Index
0
20
40
60
20000
25000
30000
35000
40000
45000

LG03

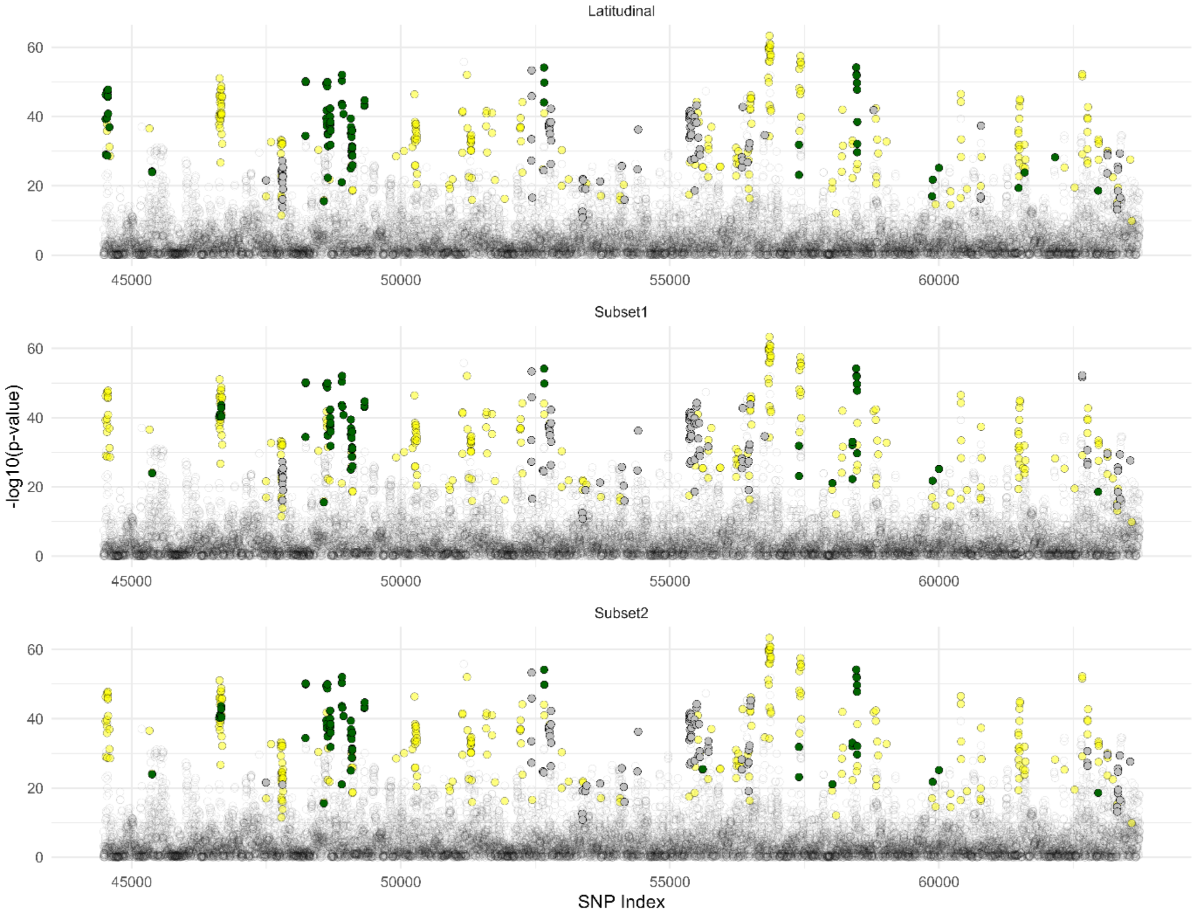

Latitudinal
Subset1
Subset2
-log10(p-value)
SNP Index
0
20
40
60
45000
50000
55000
60000

LG04

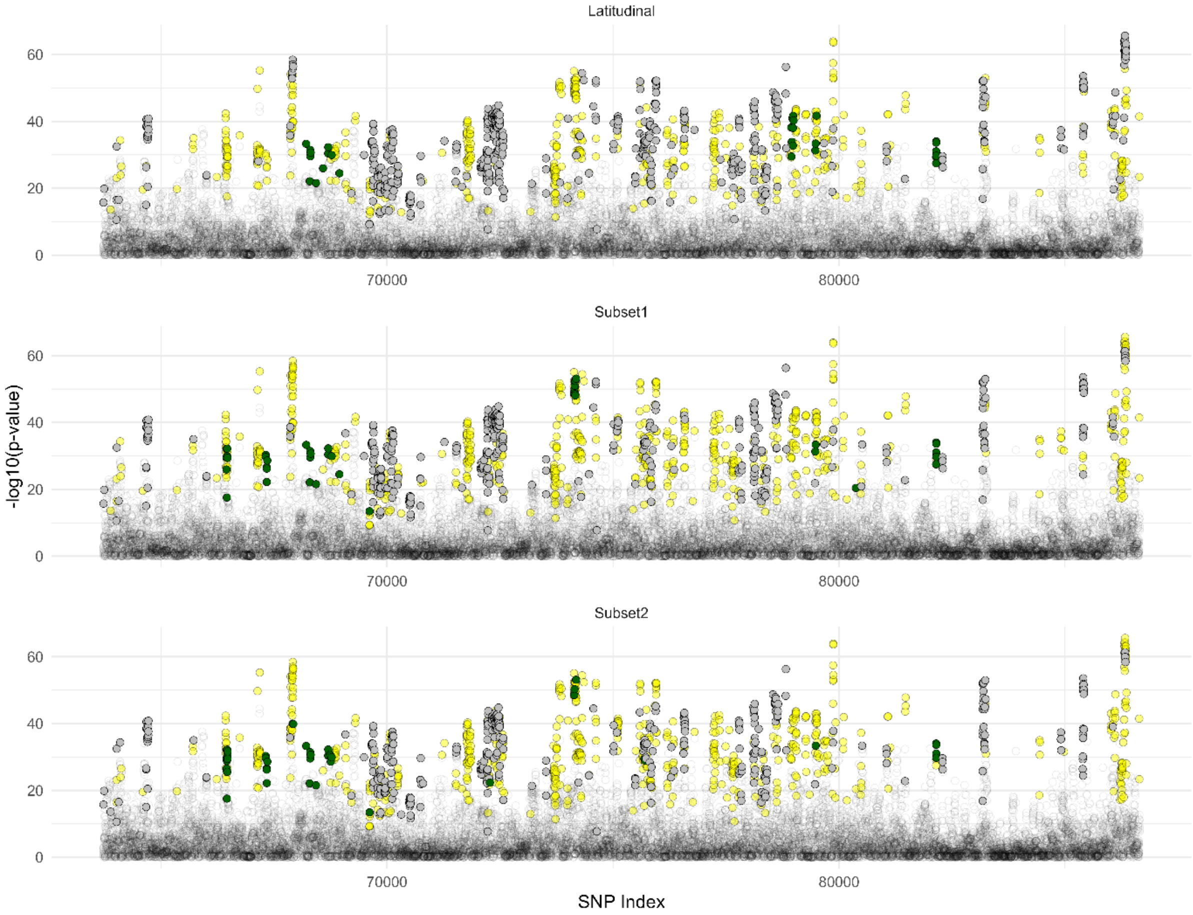
Latitudinal
Subset1
Subset2
-log10(p-value)
0
20
40
60
70000
80000
SNP Index

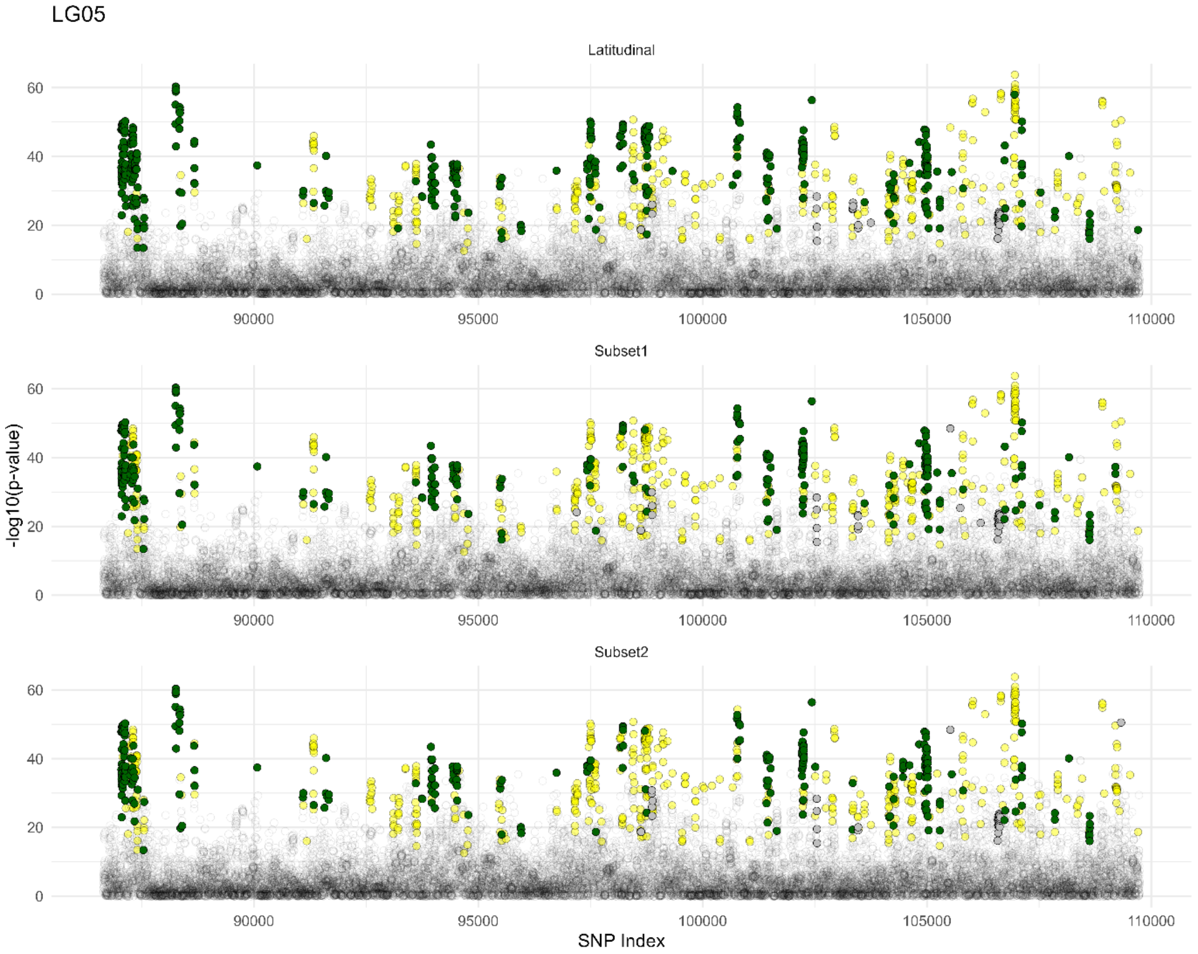
LG05
Latitudinal
Subset1
Subset2
-log10(p-value)
SNP Index
0
20
40
60
90000
95000
100000
105000
110000

LG06

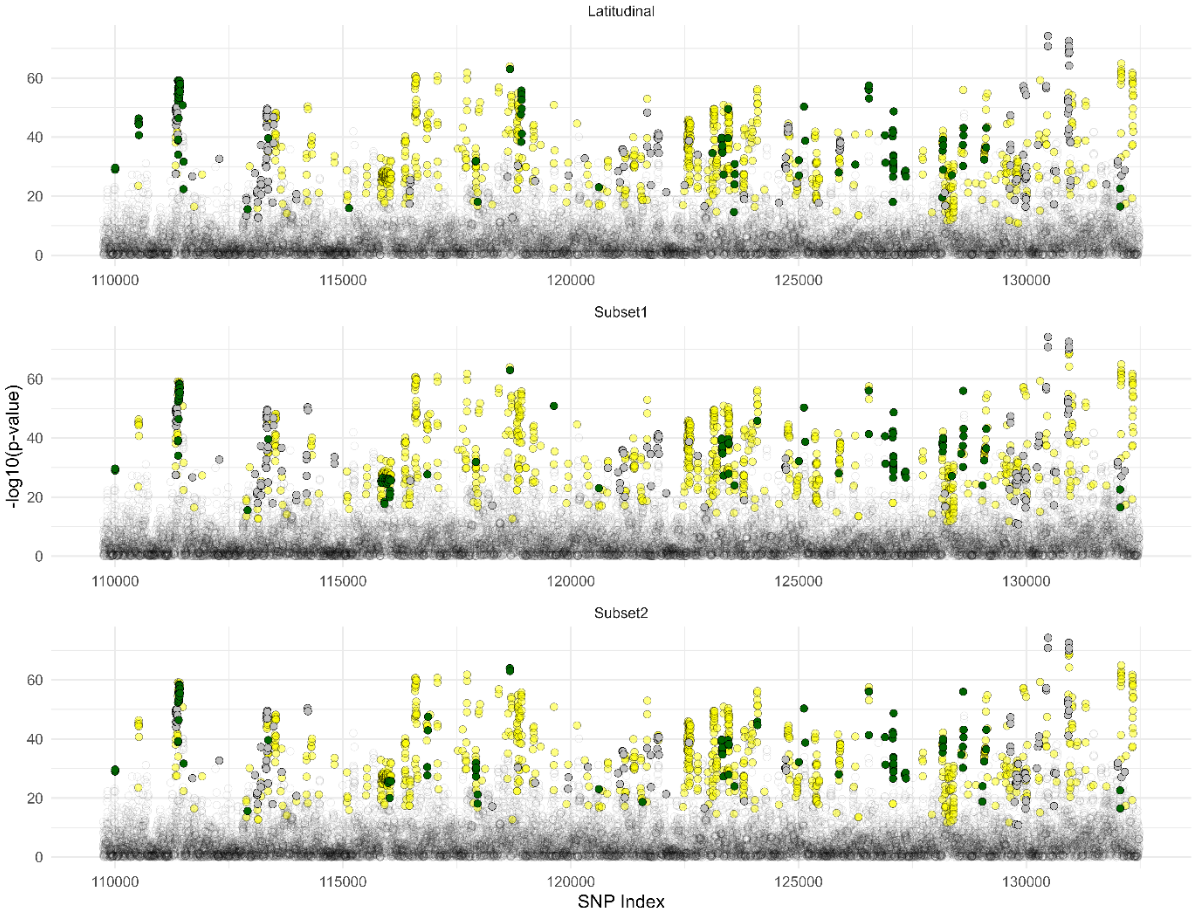
Latitudinal
Subset1
Subset2
-log10(p-value)
0
20
40
60
110000
115000
120000
125000
130000
SNP Index

LG07

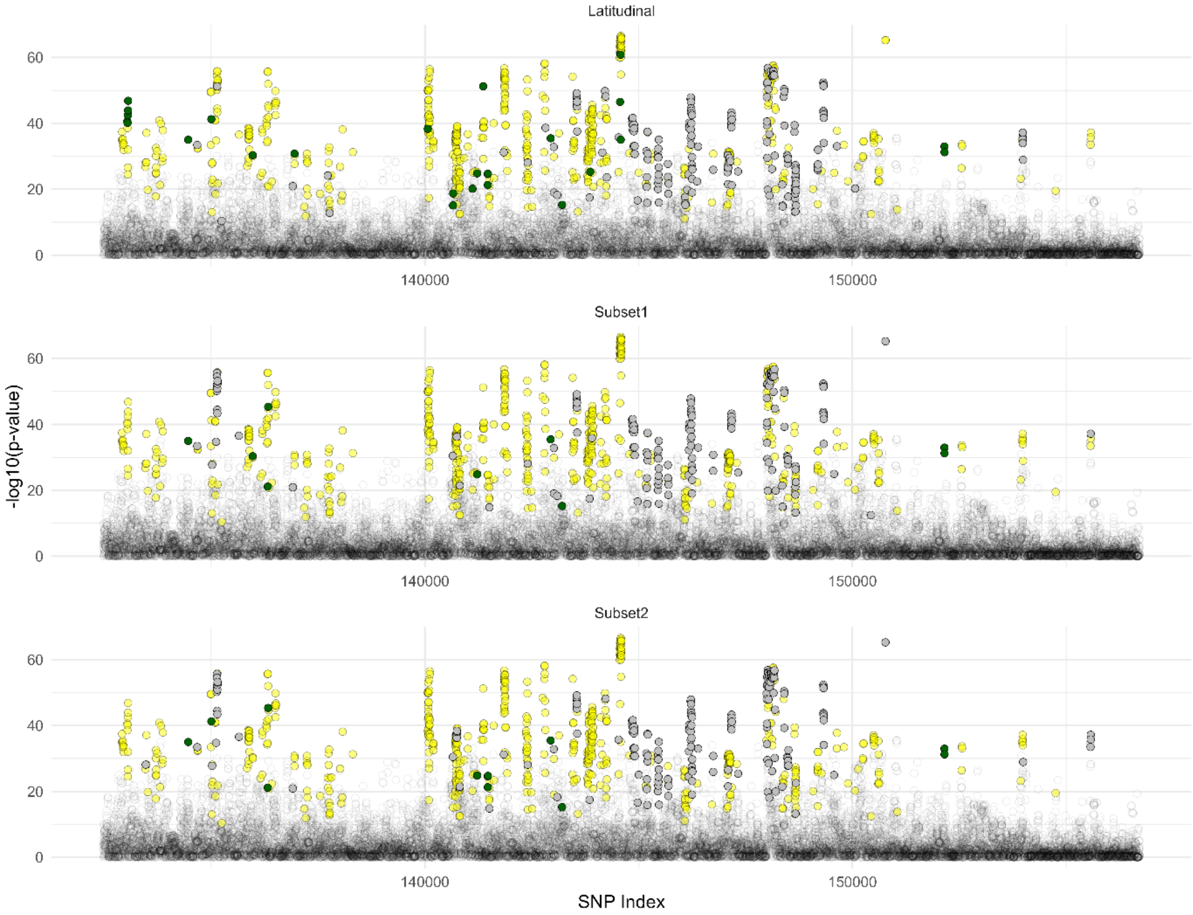
Latitudinal
Subset1
Subset2
-log10(p-value)
SNP Index
0
20
40
60
140000
150000

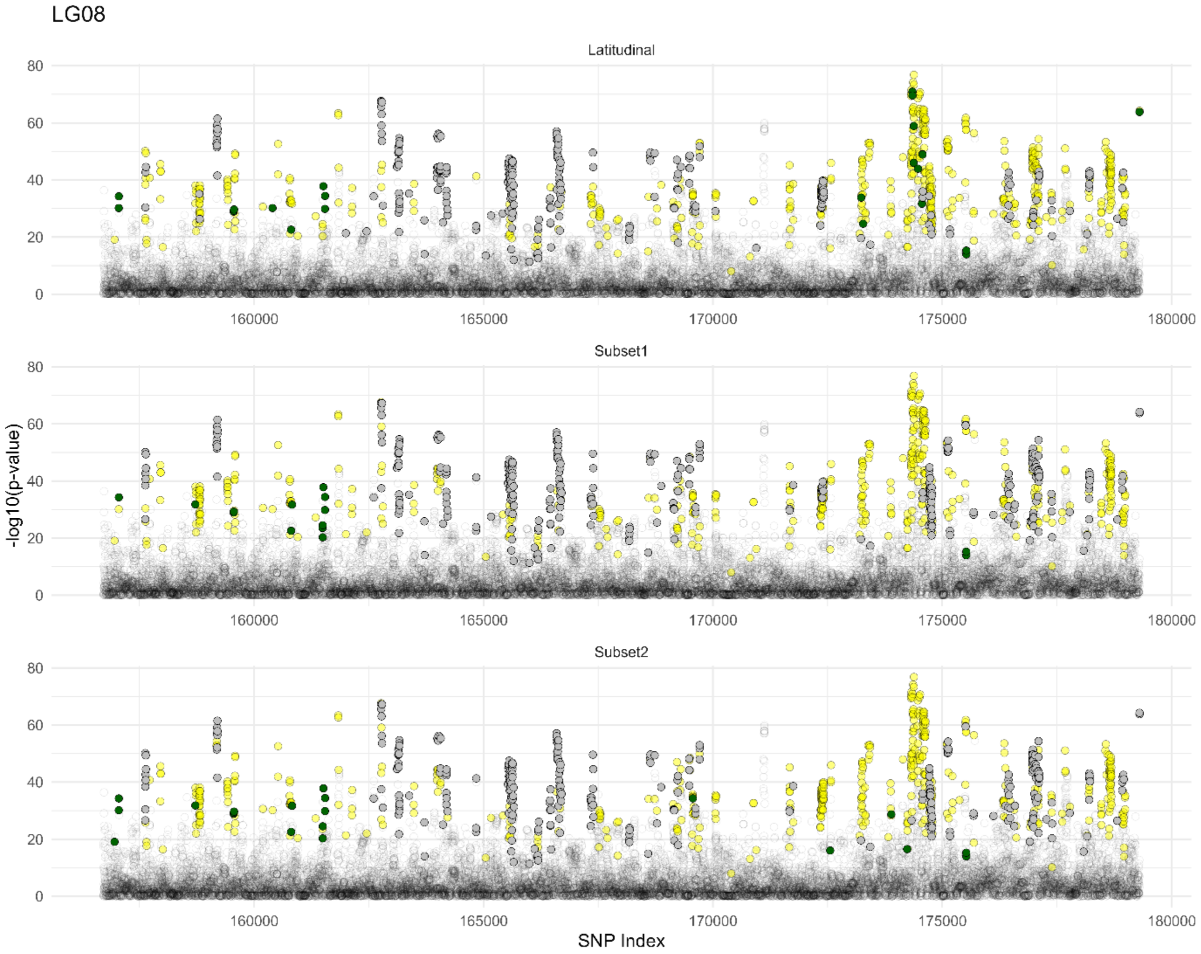
LG08
Latitudinal
Subset1
Subset2
-log10(p-value)
SNP Index
0
20
40
60
80
160000
165000
170000
175000
180000

LG09

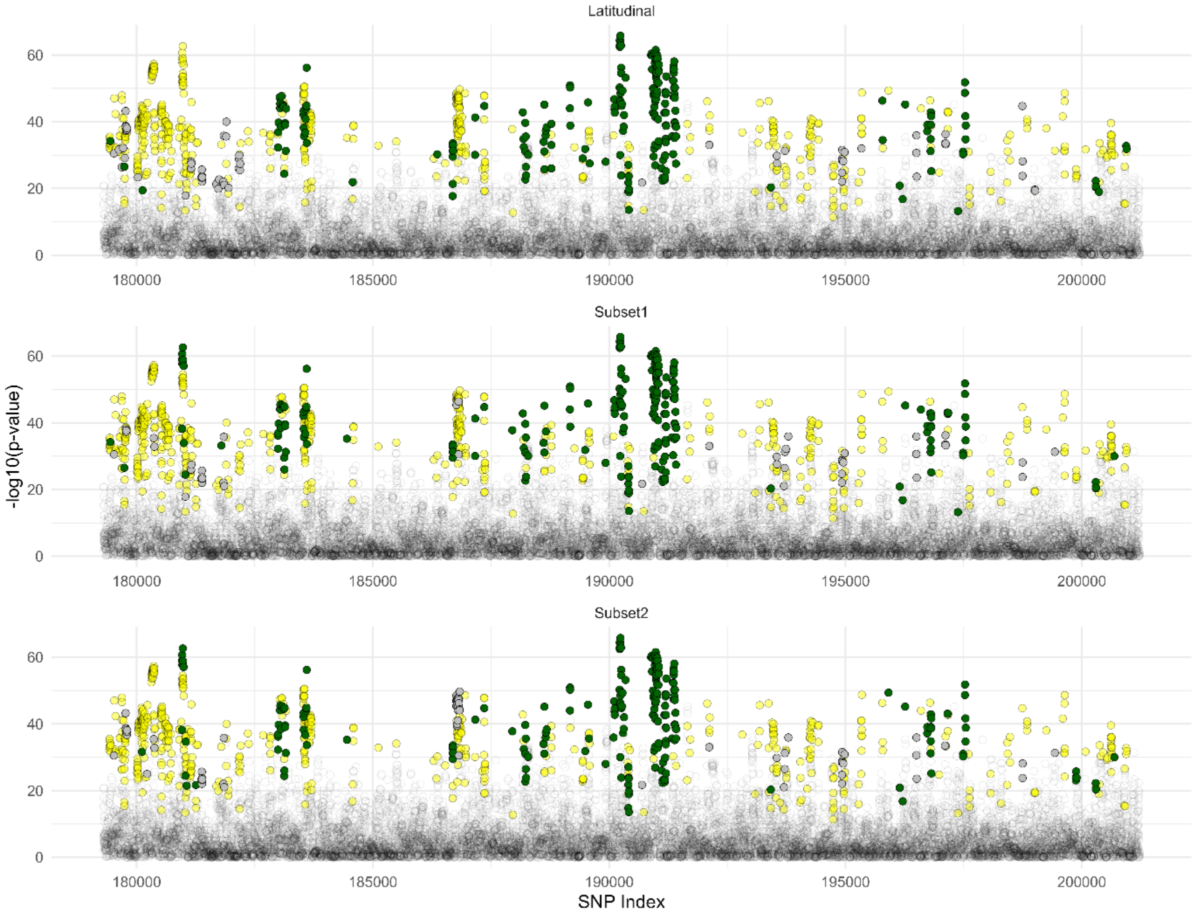
Latitudinal
Subset1
Subset2
-log10(p-value)
0
20
40
60
180000
185000
190000
195000
200000
SNP Index

LG10

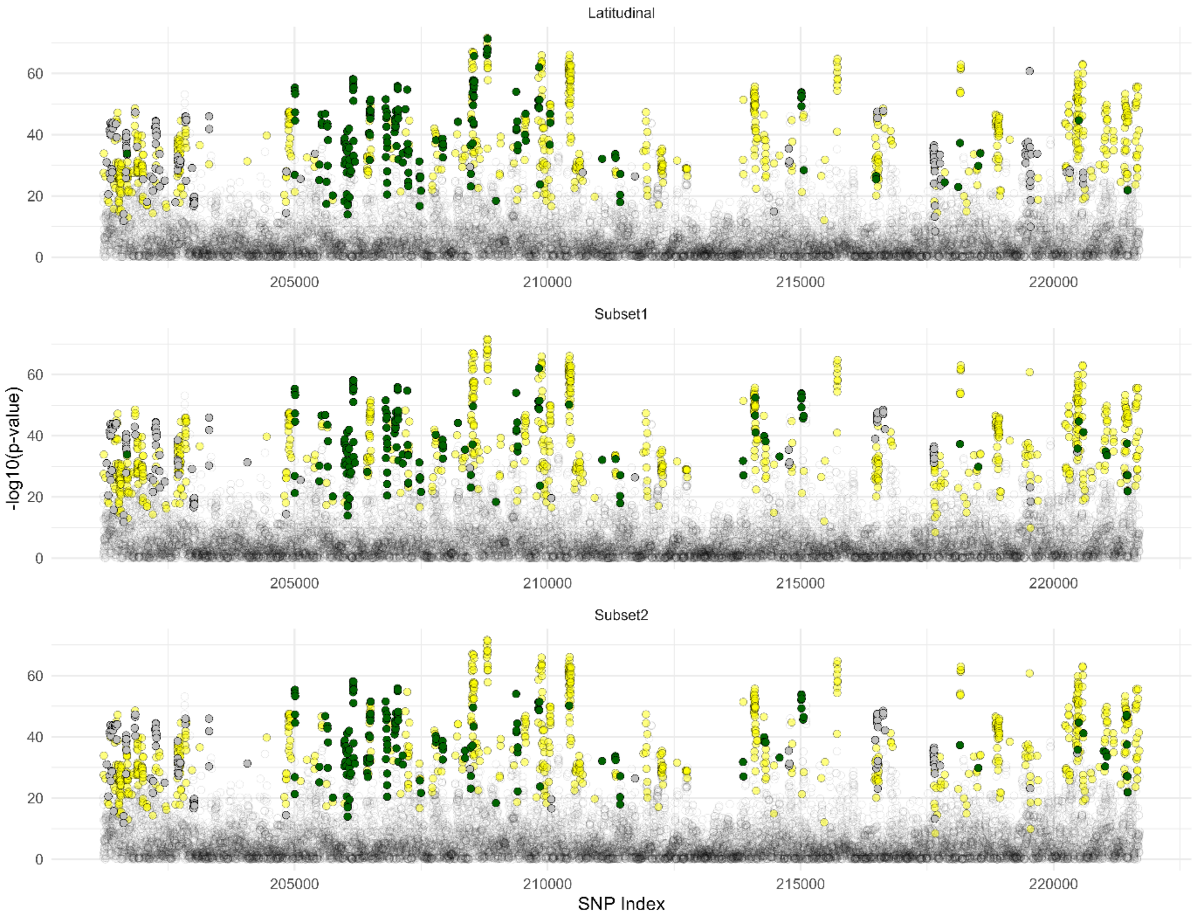
Latitudinal
Subset1
Subset2
-log10(p-value)
0
20
40
60
205000
210000
215000
220000
SNP Index

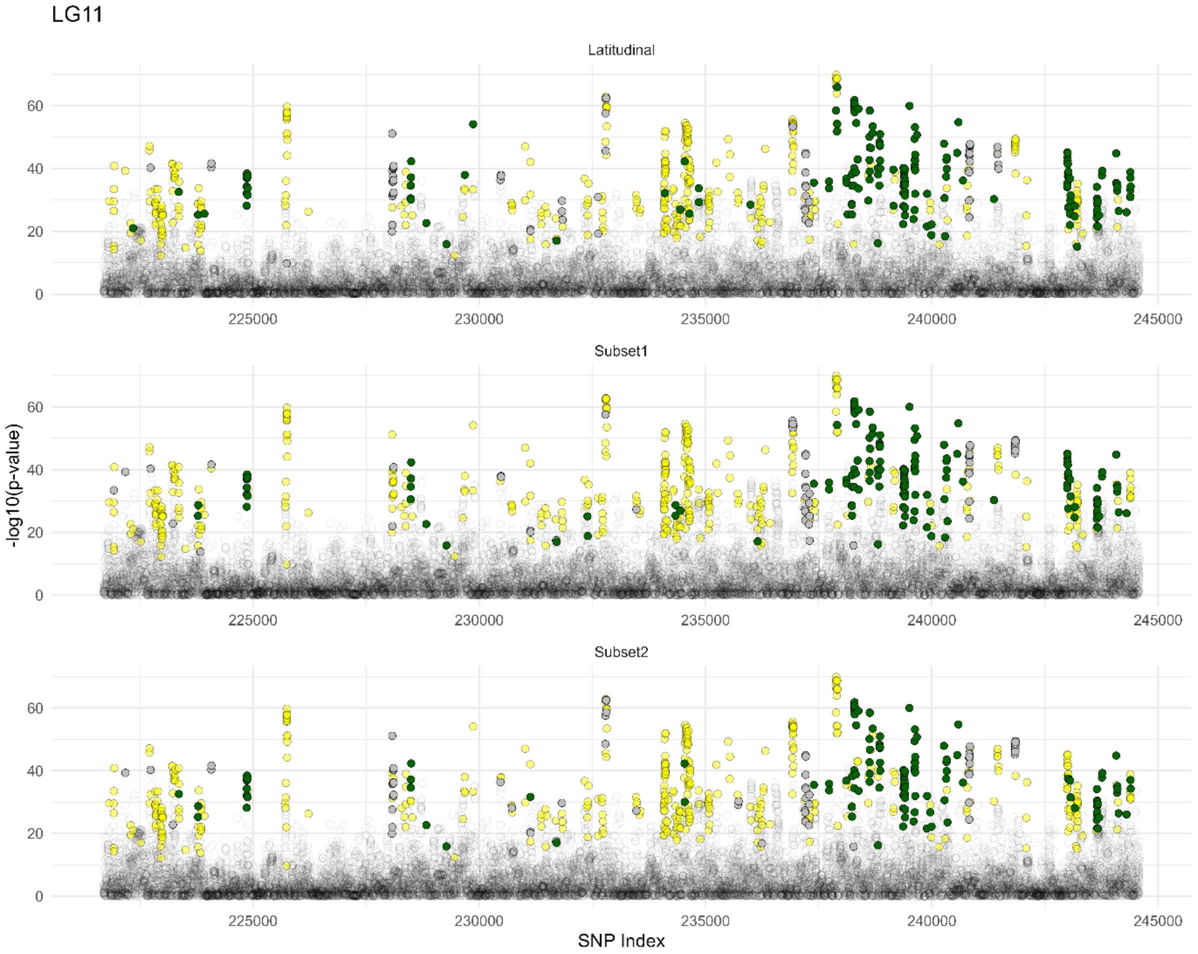
LG11
Latitudinal
Subset1
Subset2
-log10(p-value)
SNP Index
0
20
40
60
225000
230000
235000
240000
245000

LG12

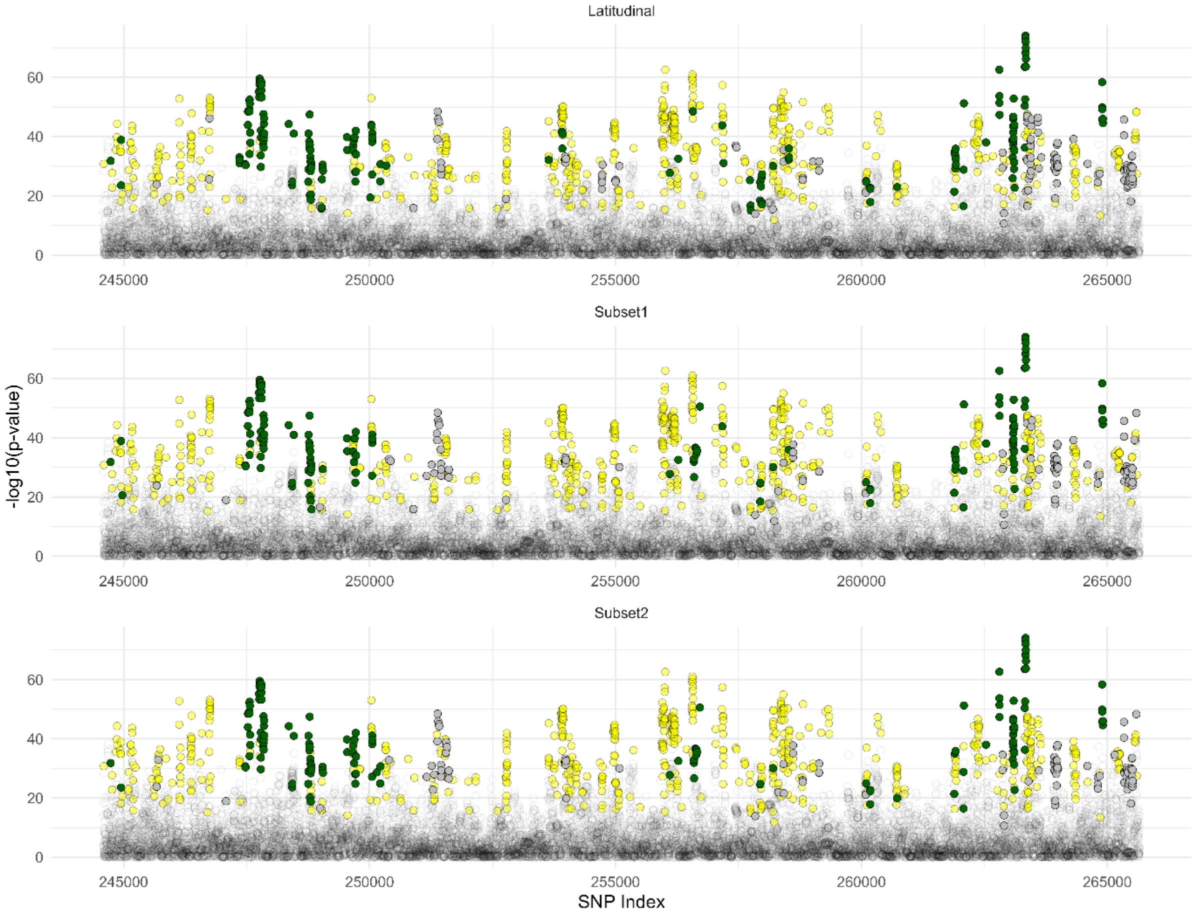

**Figure S26.** Manhattan plots of GEA p-values for the latitudinal datasets analysis and the two additional latitudinal subsets, aligned by genomic position according to the super-scaffold assembly. SNPs showing ancestry excess in the genomic clines (*BGC*) analyses are represented by triangles, coloured as follows: dark green indicates *P. engelmannii* ancestry excess, and grey indicates *P. glauca* ancestry excess. Yellow circles represent SNPs tested in the *BGC* analysis (parental frequency differential > 0.6) but not showing ancestry excess. Small black and transparent circles denote SNPs included only in the GEA analysis but not in the *BGC* analysis.

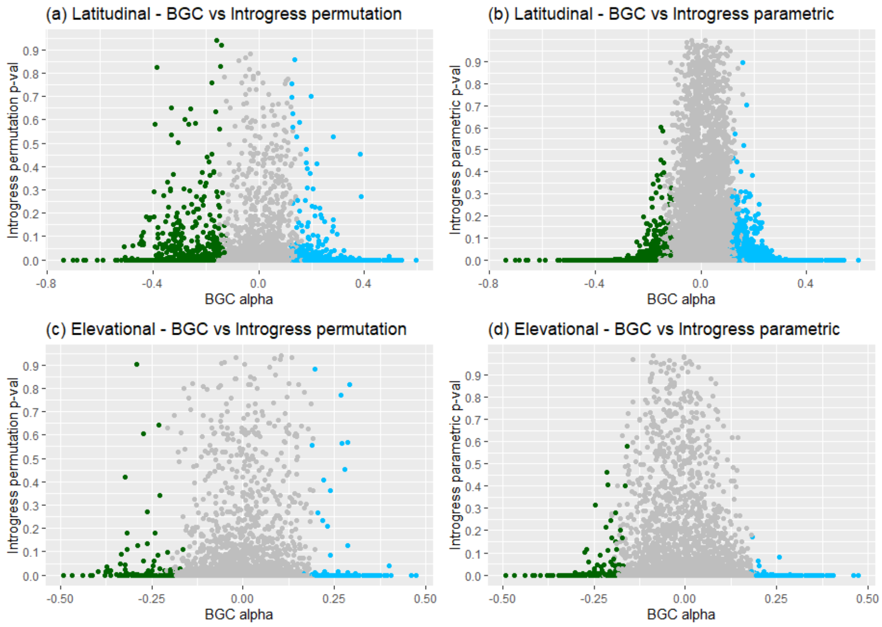

**Figure S27.** Comparison of *BGC* α-values with p-values obtained using *Introgress*, based on both permutation tests (panels a and c) and parametric tests (panels b and d).

Elevational dataset LG01

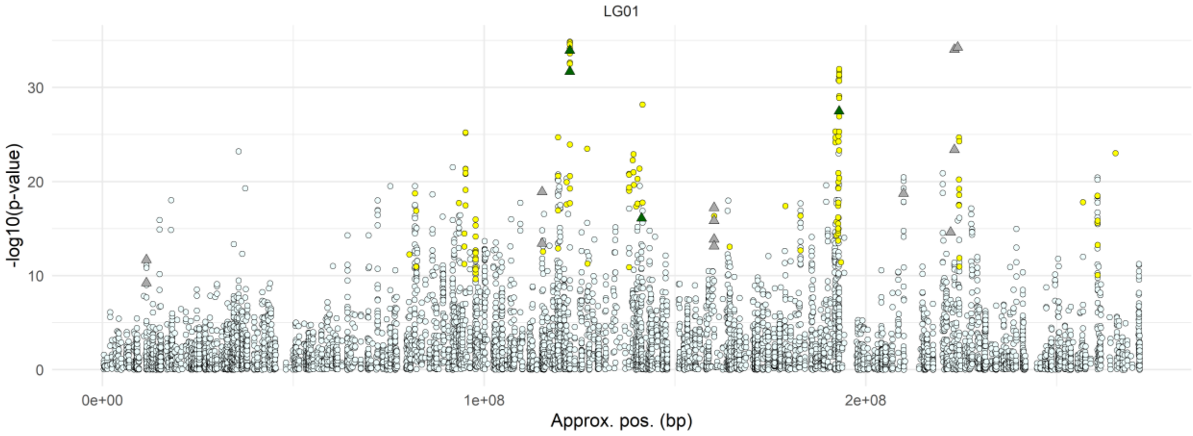

Latitudinal dataset LG01

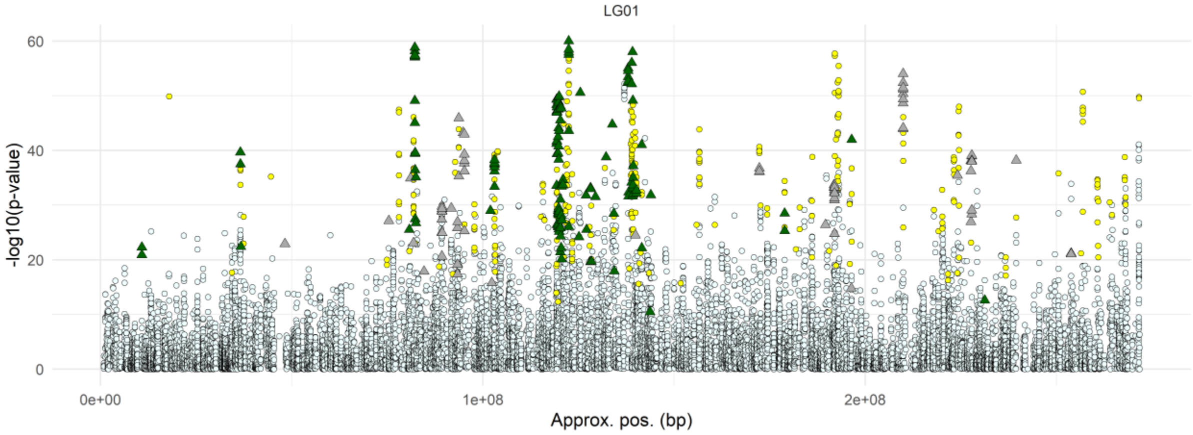

Elevational dataset LG02

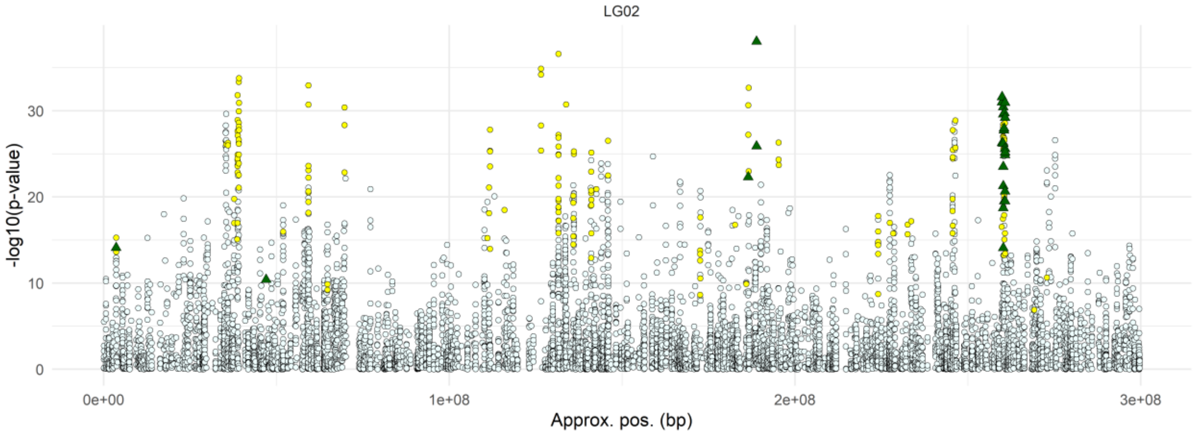

Latitudinal dataset LG02

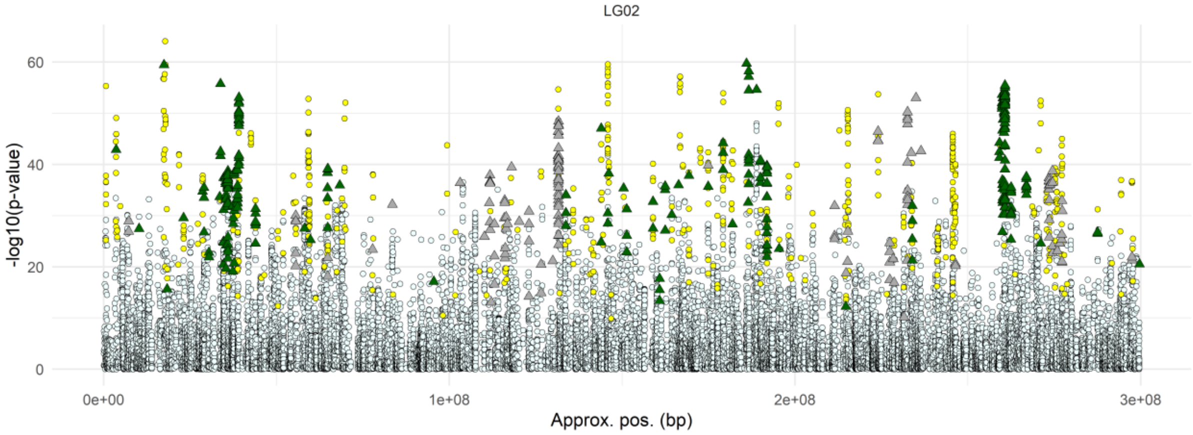

Elevational dataset LG03

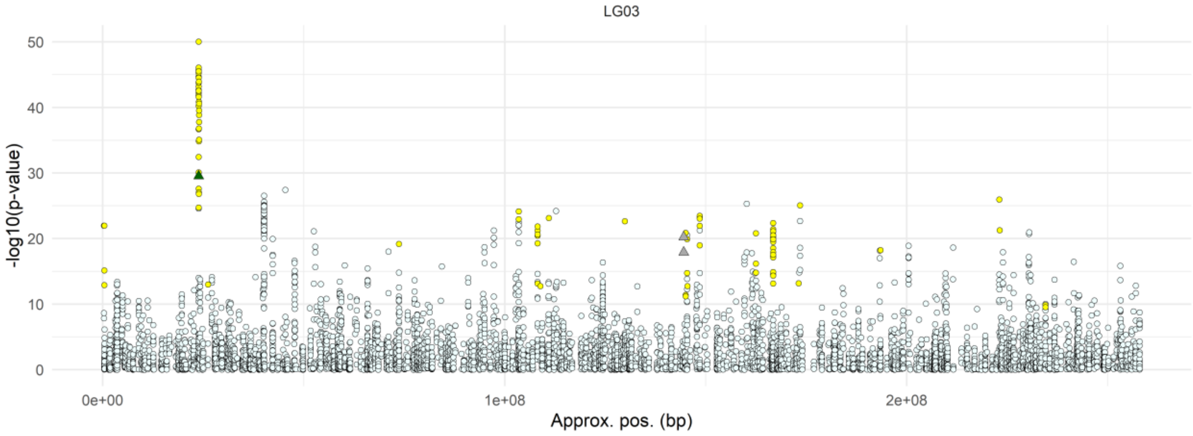

Latitudinal dataset LG03

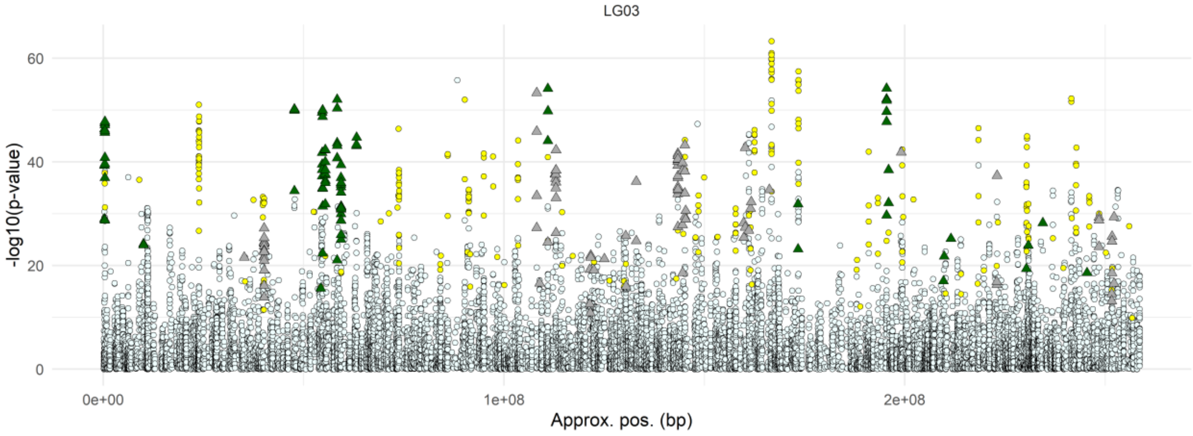

Elevational dataset LG04

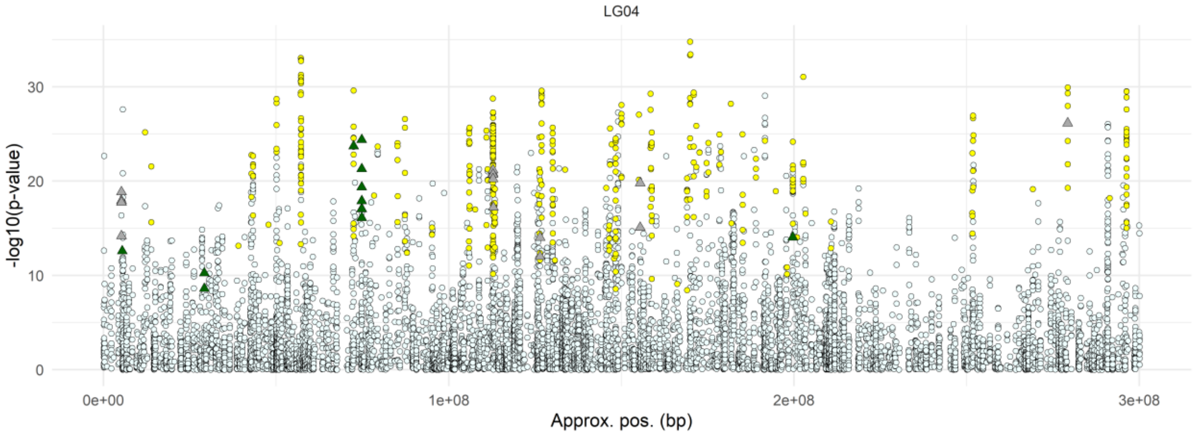

Latitudinal dataset LG04

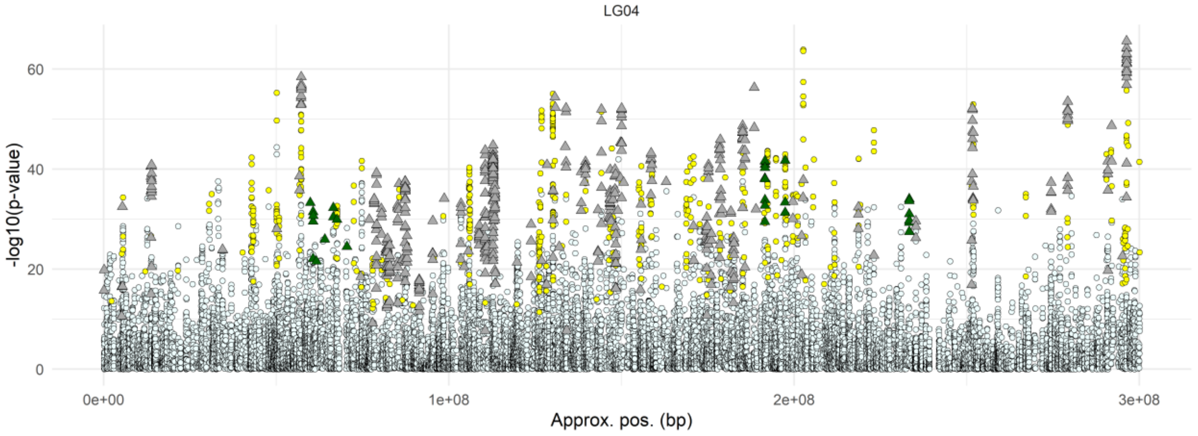

Elevational dataset LG05

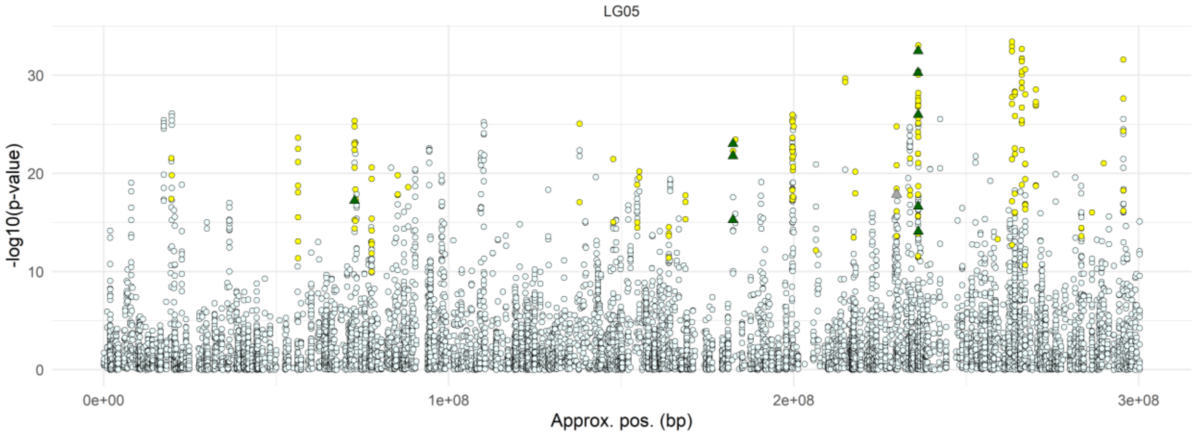

Latitudinal dataset LG05

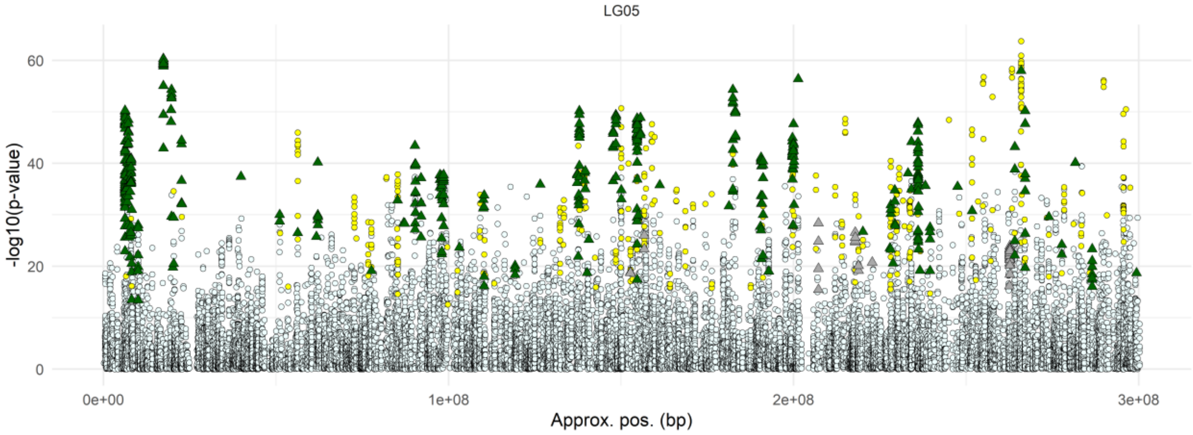

Elevational dataset LG06

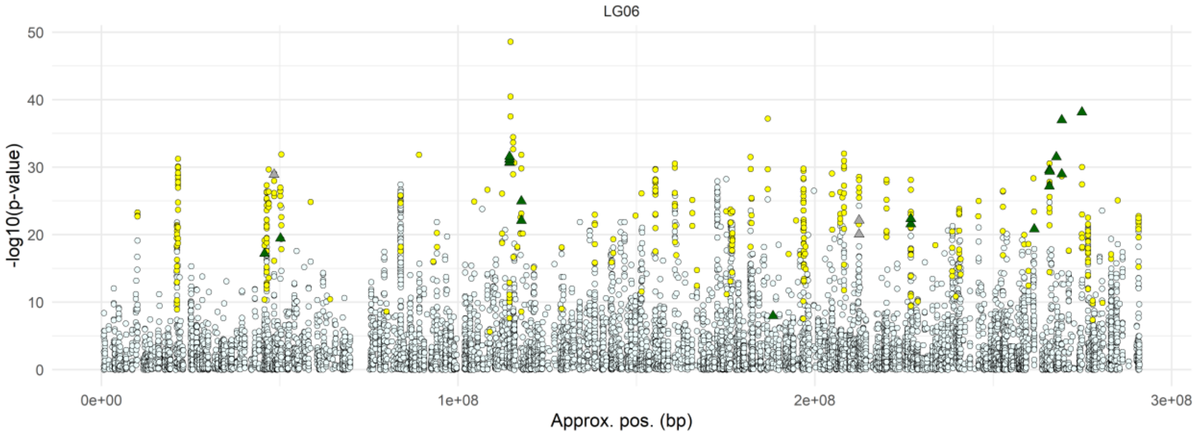

Latitudinal dataset LG06

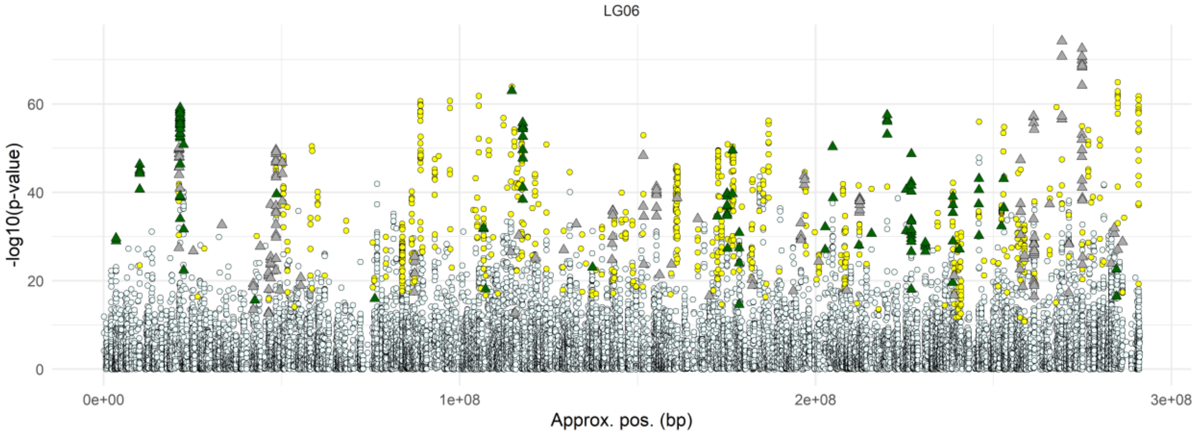

Elevational dataset LG07

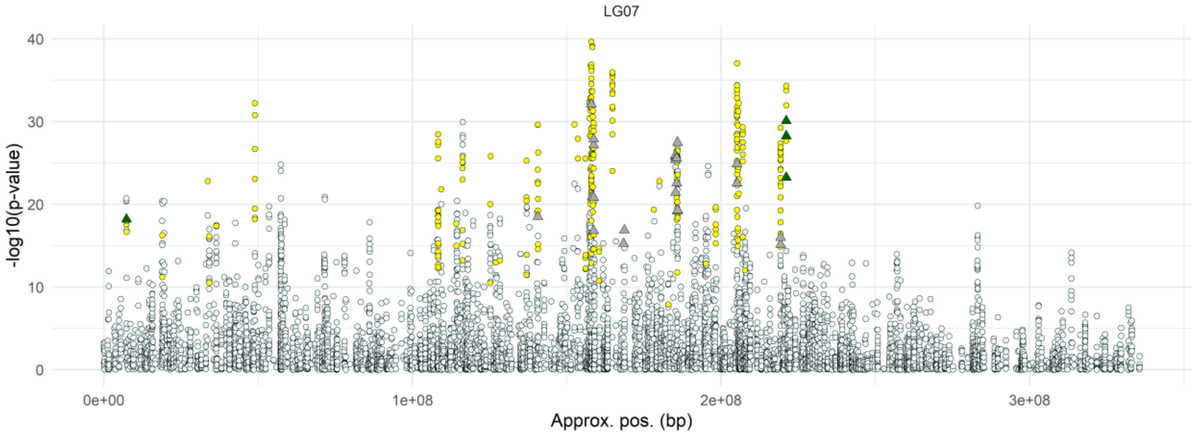

Latitudinal dataset LG07

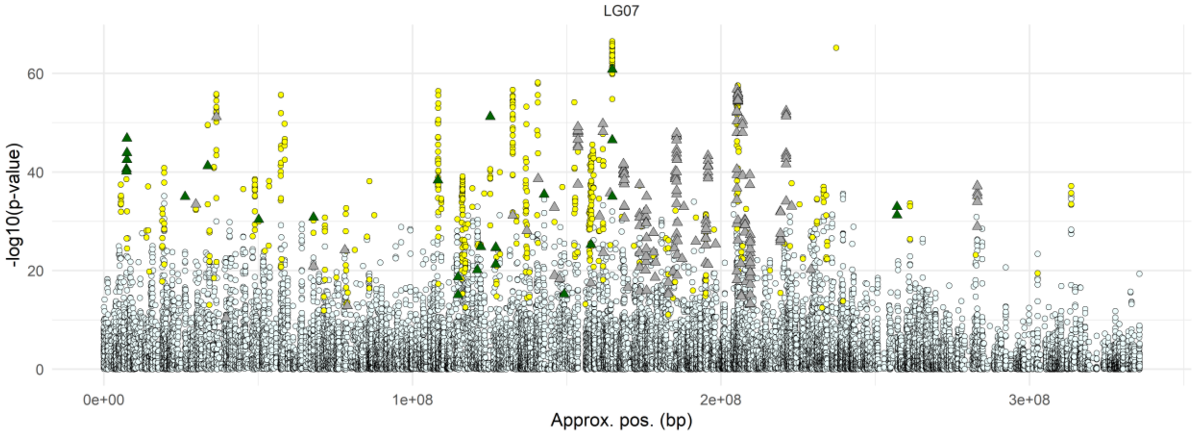

Elevational dataset LG08

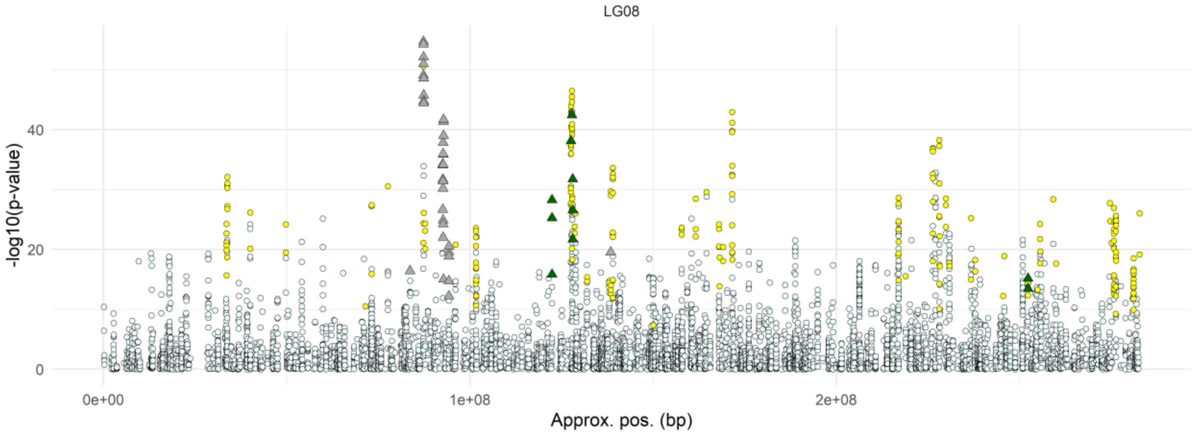

Latitudinal dataset LG08

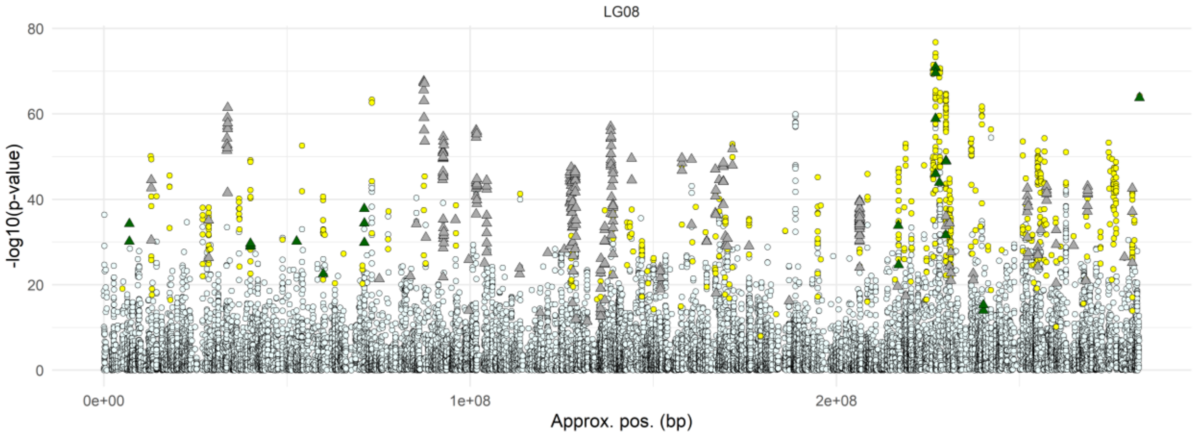

Elevational dataset LG09

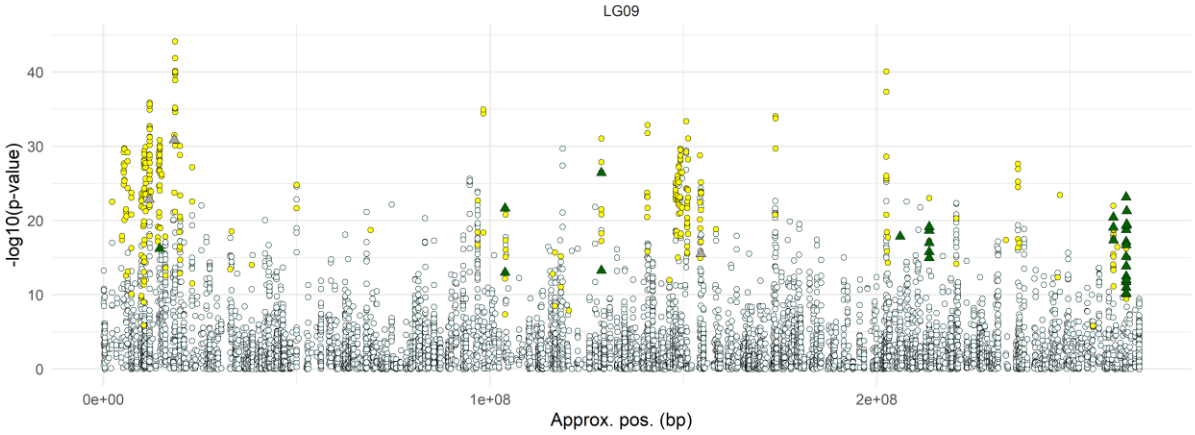

Latitudinal dataset LG09

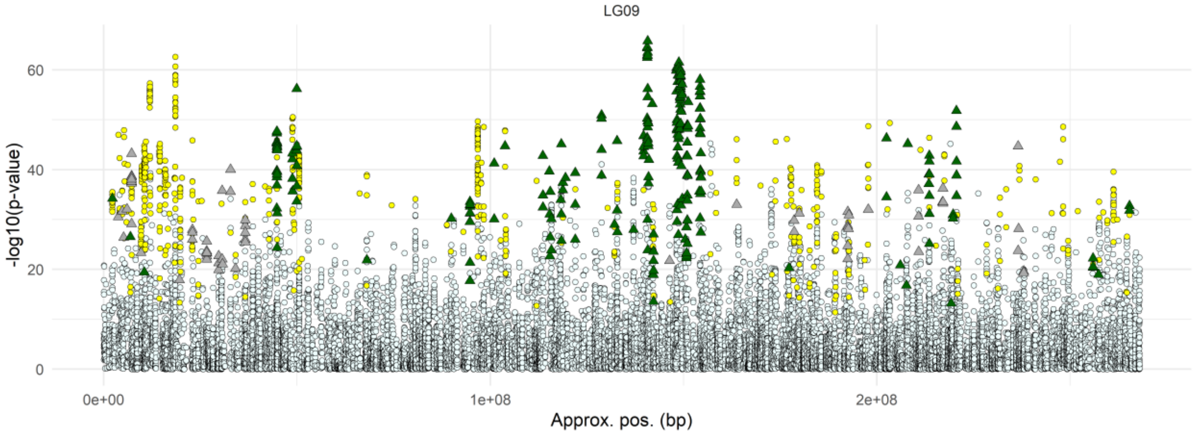

Elevational dataset LG10

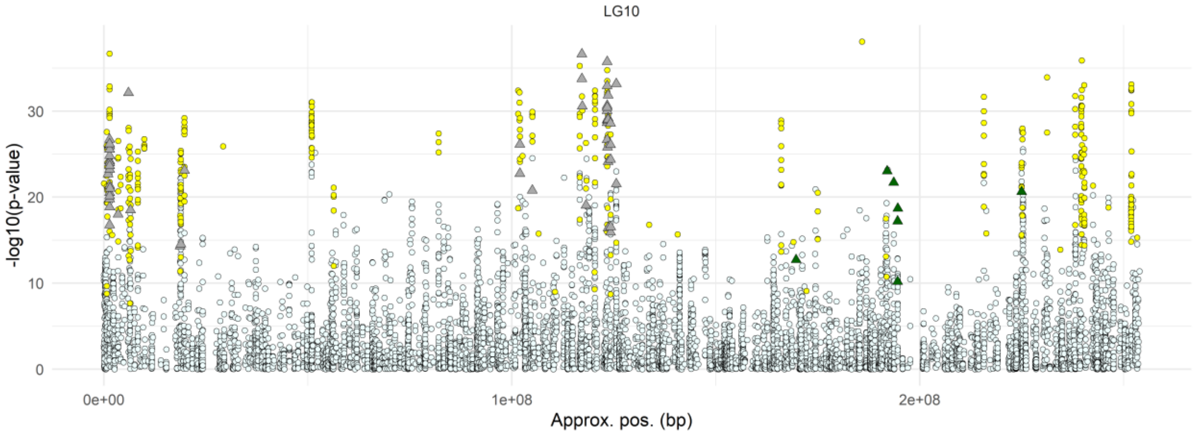

Latitudinal dataset LG10

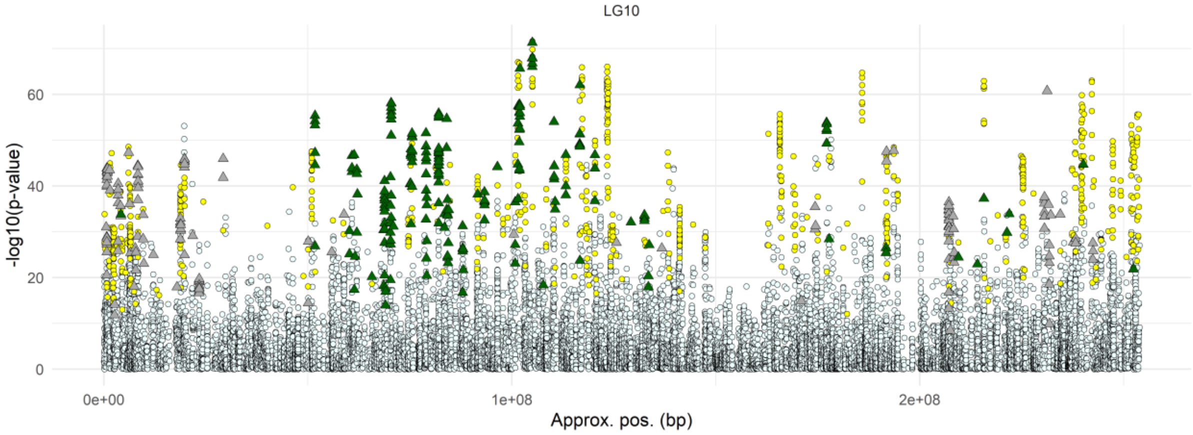

Elevational dataset LG11

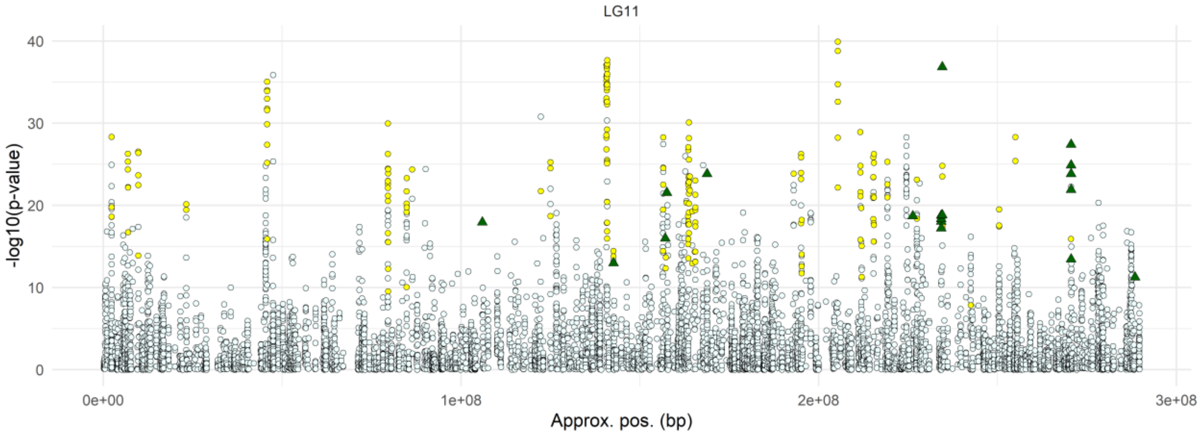

Latitudinal dataset LG11

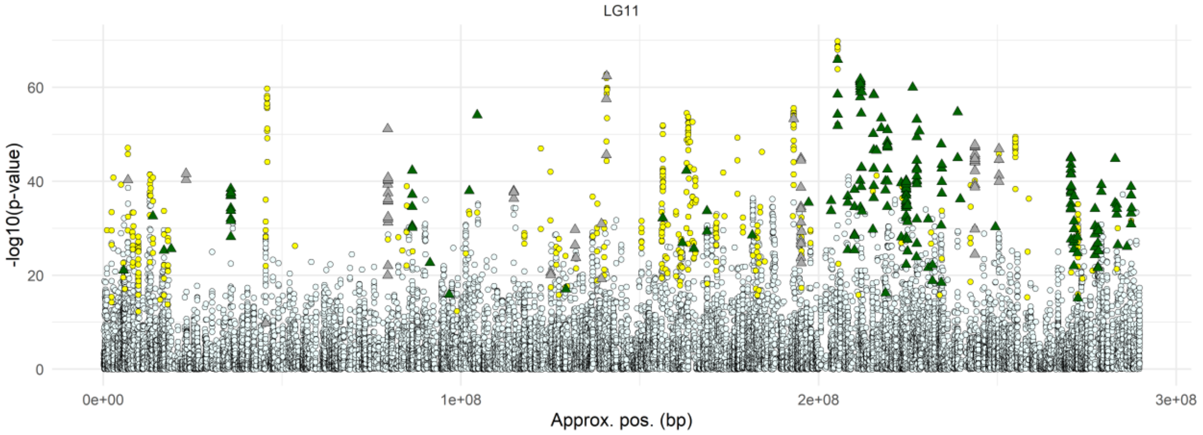

Elevational dataset LG12

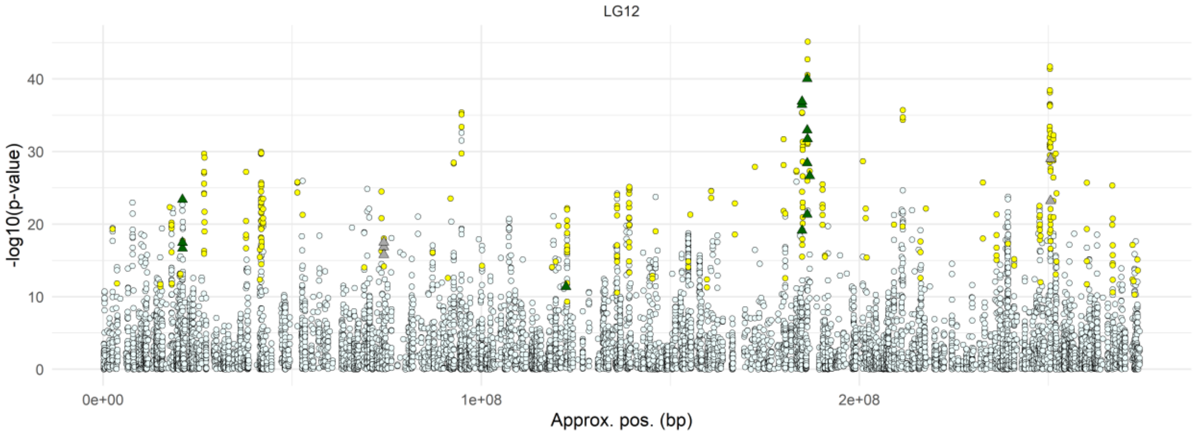

Latitudinal dataset LG12

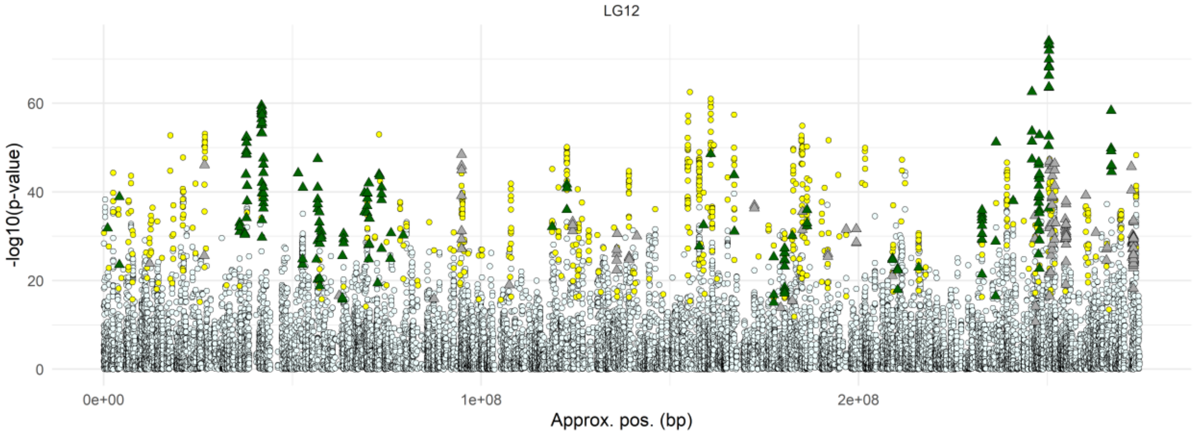

**Figure S28.** Manhattan plots of GEA p-values for the elevational and latitudinal datasets, aligned by genomic position according to our super-scaffold assembly. SNPs showing ancestry excess in the genomic clines (*BGC*) analyses are represented by triangles, coloured as follows: dark green indicates *P. engelmannii* ancestry excess, and grey indicates *P. glauca* ancestry excess. Yellow circles represent SNPs tested in the *BGC* analysis (parental frequency differential > 0.6) but not showing ancestry excess. Small pale-azure circles denote SNPs included only in the GEA analysis but not in the *BGC* analysis.

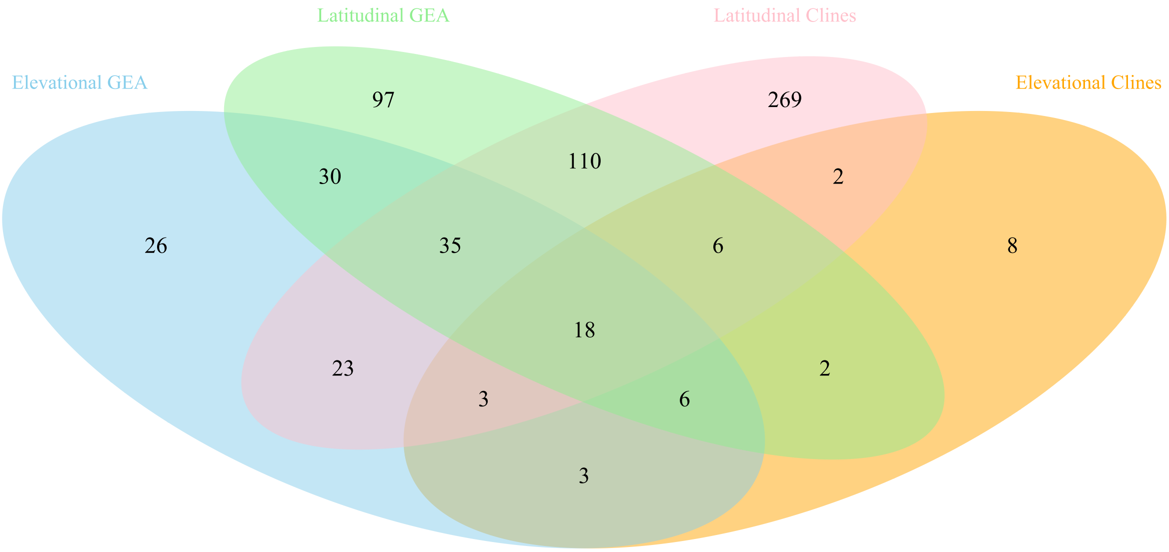

**Figure S29.** Venn diagram showing the overlap of candidate adaptive genes identified across the four analyses: Elevational GEA, Elevational Genomic Clines (*BGC*), Latitudinal GEA and Latitudinal Genomic Clines (*BGC*). Note that the elevational GEA tested 2,901 genes, the latitudinal GEA 6,186 genes, the elevational genomic clines 296 genes, and the latitudinal genomic clines 992 genes. 2,734 genes were shared between the two GEAs, while 257 genes were shared between the two genomic clines analyses.

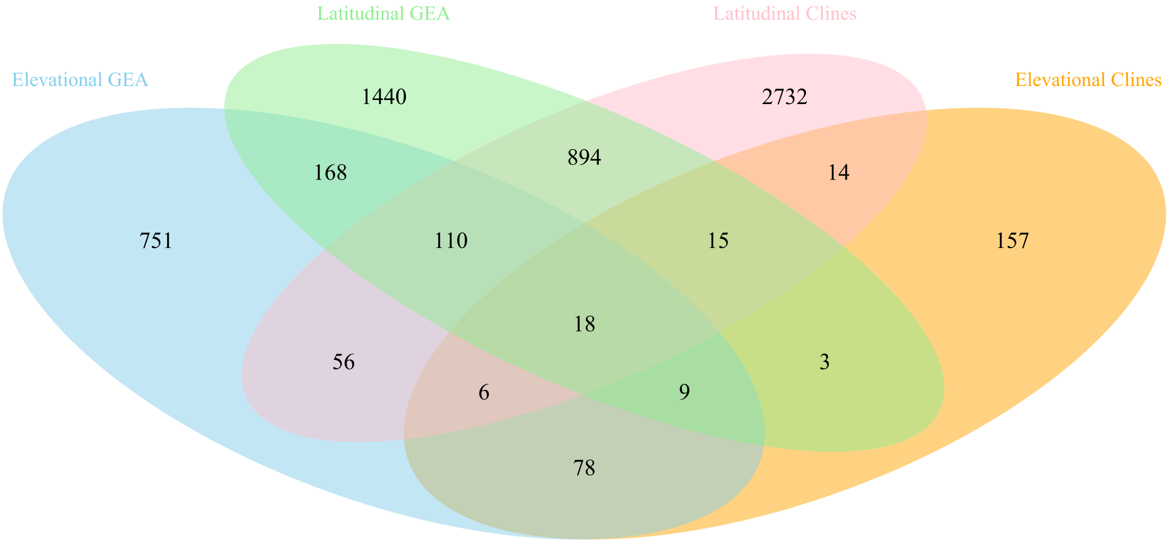

**Figure S30.** Venn diagram showing the overlap of candidate SNPs identified across the four analyses: Elevational GEA, Elevational Genomic Clines (*BGC*), Latitudinal GEA and Latitudinal Genomic Clines (*BGC*). Note that the elevational GEA tested 119,664 SNPs, the latitudinal GEA 265,655 SNPs, the elevational genomic clines 3,256 SNPs, and the latitudinal genomic clines 10,154 SNPs. 31,667 SNPs were shared between the two GEAs, while 982 were shared between the two genomic clines analyses.

**Table S1.** Annotation of the 18 candidate genes identified by all four analyses.

| Gene ID | Description (Pfam/InterPro) |
|---|---|
| DB47_00005558 | RNA recognition Motif |
| DB47_00005560 | Ribosomal protein L30p/L7e |
| DB47_00045432 | Protein tyrosine and serine/threonine kinase Leucine rich repeat N-terminal domain |
| DB47_00046462 | NA |
| DB47_00024128 | Cytochrome P450 |
| DB47_00066514 | U-box domain |
| DB47_00079589 | Kelch repeat domain, E3 ligase activity |
| DB47_00088464 | Myb/SANT-like DNA-binding domain |
| DB47_00035975 | Agenet domain |
| DB47_00029886 | HhH-GPD superfamily base excision DNA repair protein |
| DB47_00025922 | Oxysterol-binding protein |
| DB47_00095825 | C2 domain |
| DB47_00051791 | Protein kinase |
| DB47_00078734 | This entry represents a family of uncharacterised proteins, including chloroplastic UV-B-induced protein At3g17800 from Arabidopsis. |
| DB47_00085790 | ATP synthase delta (OSCP) subunit |
| DB47_00052926 | Leucine rich repeat N-terminal domain |
| DB47_00047065 | Protein kinase |
| DB47_00054068 | Inorganic H+ pyrophosphatase |

**Table S2.** Genes within our broader set of 638 candidates (identified by either analysis) also reported to contribute to local adaptation in other conifer studies.

| *P. glauca* gene ID | Gene ID in compared species | Description /Function (Pfam/InterPro) |
|---|---|---|
| | ***P. glauca* in eastern Canada – Hornoy et al. (2015).** | |
| No GFF Annotation | GQ02903_I19 (temperature) | Unknown |
| DB47_00029799 | GQ04011_G20 (temperature) | AP2/ERF domain |
| DB47_00016142 | GQ02829_F03 (temperature) | Ubiquitin-conjugating-enzyme |
| DB47_00000346 | GQ03211_G05 (temperature) | Rieske Fe-S proteins of Cytochrome_B6 |
| No GFF Annotation | GQ03919_G08 (temperature) | Unknown |
| DB47_00023396 | GQ03507_B21 (temperature) | Leucin-rich repeat protein kinase |
| DB47_00065457 | GQ03114_J11 (precipitation) | PAP2 superfamily |
| | ***Picea rubens* Sarg. in northeastern North America - Capblancq et al. (2022)** | |
| DB47_00034473 | MA_102606g0010 | Chromatin remodelling 4-like isoform X1 |
| DB47_00059053 | MA_10432806g0020 | LAG1 longevity assurance homolog 3-like |
| DB47_00064461 | MA_10434569g0010 | Glycerol-3-phosphate dehydrogenase (NAD+) cytosolic-like |
| DB47_00052451 | MA_10435314g0010 | Flowering time control (FPA) |
| DB47_00035208 | MA_10437228g0010 | Probable cyclic nucleotide-gated ion channel chloroplastic |
| DB47_00055335 | MA_132866g0010 | Probable ubiquitin-conjugating enzyme E2 18 |
| DB47_00043788 | MA_14836g0010 | Transport inhibitor response 1-like |
| | ***Pinus contorta* in western North America – Yeaman et al. (2016)** | |
| DB47_00054460 | comp10479_c0_seq1.p1 | Zinc finger, C3HC4 type (ring-type finger) |
| DB47_00005245 | comp11554_c0_seq1.p1 | TCP family transcription factor |
| DB47_00034340 | comp12027_c0_seq1.p1 | Exostosin GT47 domain |
| DB47_00013080 DB47_00013080 | comp1438_c0_seq1.p1 | Unknown |

| | | |
|---|---|---|
| DB47_00011918 | comp20671_c0_seq1.p1 | Protein kinase domain |
| DB47_00047312 | comp2224_c0_seq1.p1 | Ring-type zinc finger |
| DB47_00008815 | comp26675_c0_seq1.p1 | Poly (ADP-ribose) polymerase catalytic domain |
| DB47_00099276 | comp3285_c0_seq1.p1 | Zinc-finger |
| DB47_00026687 | comp365_c0_seq1.p1 | Glyceraldehyde 3-phosphate dehydrogenase, NAD binding domain |
| DB47_00015009 | comp6567_c0_seq1.p1 | ATP-dependent Clp protease adaptor protein ClpS |
| DB47_00066488 | comp7786_c0_seq1.p1 | AP2/ERF domain |
| DB47_00089623 | comp9858_c0_seq1.p1 | Zinc-finger of the FCS-type, C2-C2 |
| | ***P. glauca* in Quebec – Depardieu et al. 2021** | |
| DB47_00099276 | GQ03707_G19 (summer soil moisture index) | Zinc finger protein |
